# Orthogonal Inducible Transcriptional Control Systems for Investigation of Gene-Gene Interactions

**DOI:** 10.1101/2025.10.28.685021

**Authors:** Aslı Azizoğlu

## Abstract

Genes do not act in isolation but through complex networks of interactions. While many double and triple gene interactions have been uncovered in *Saccharomyces cerevisiae*, most have been identified through overexpression or loss-of-function studies, leaving dosage-dependent interactions largely unexplored. Multi-gene control systems suffer from various drawbacks, including limited range and high cell-to-cell variation of expression across the population. To address this gap, we extend a recently developed inducible expression system that enables precise, tunable control of gene expression to additional repressor-inducer pairs: LacI/IPTG and LexA-hER/β-estradiol. We demonstrate that both new systems can control endogenous genes on their own by placing various low, medium and high expressed endogenous genes under their control. Using all three WTC systems, we show simultaneous and orthogonal double and triple gene control that confirms known gene-gene interactions within the anaphase signaling network and reveals new dosage-dependent phenotypes. Given their precision, orthogonality and compatibility with all commonly used growth media, this collection of WTC systems represents a new tool suitable for precise investigation of dosage-dependent gene-gene interactions in yeast.

## Introduction

The yeast genome is a highly connected network^1^. *S.cerevisiae* can tolerate deletions of about 80% of its Open Reading Frames (ORF) without lethal effects. These genes are non-essential for life presumably because there are other genes with similar or redundant functions. Underlining this redundancy and interconnectivity, synthetic Gene Array screens (SGA) have revealed many cases where deletion or overexpression of one gene is well-tolerated until another gene is perturbed^2,3^. There are also genes whose overexpression is detrimental to the cell, but overexpression of another gene can alleviate this effect^4,5^. As high throughput methods develop further, trigenic interactions and redundancies are also being discovered^6,7^. Even genes that are considered essential can have “bypass” partners; genes which upon acquiring a mutation can compensate for the deletion of an essential gene^8,9^.

This abundance of genetic interactions means that understanding the function of certain genes and phenotypes emerging from gene-gene interactions requires experimental perturbation of more than simply a single gene. For example, the role of phosphoinositide-dependent kinases in heat shock response was only discovered once the products of the three paralogs PDK1, PDK2 and PDK3 were all simultaneously depleted^10^. Additionally, questions about dosage sensitivity and suppression require simultaneous and quantitative perturbation of multiple genes. For example, while yeast is sensitive to overexpression of the mitotic phosphatase Cdc14, this effect can be suppressed by simultaneous overexpression of the Cdc14 inhibitor Net1^5^. Nevertheless, the exact dosage balance required to maintain normal cellular function, and the threshold beyond which this balance becomes harmful, remain undetermined. Such dosage-dependent interactions are relevant to understanding varying penetrance of not just yeast but also human traits, including disease alleles^11^.

Thus far, most screens of genetic interactions were done using deletion or overexpression collections and temperature sensitive alleles. However, temperature sensitive (*ts*) alleles might not impair protein function completely and they alter the physiological state of the cell due to temperature shifts. And neither *ts* alleles, nor deletion or overexpression libraries lend themselves to investigating dosage effects precisely. There are methods, such as the Tug-of-War method, that can determine whether a dosage imbalance between two proteins is tolerated or not^12,13^. In Tug-of-War, an endogenous gene is expressed on a plasmid, and the cell is incentivized to increase the copy number of this plasmid. If there is a dosage-dependent interaction between two genes expressed on two separate plasmids, their copy number is expected to increase in tandem. While this method can provide information on which genes are interacting in a dosage-dependent manner, questions about where the threshold of dosage imbalance between two interacting proteins like Cdc14 and Net1 might lie would require precise titration of these gene products.

Inducible promoters or CRISPR interference^14^ (CRISPRi) based methods are better suited for such applications. Recently a genome-scale library based on β-estradiol inducible promoters and another based on CRISPRi have been built, demonstrating the potential of titratable gene expression in elucidating genome-wide dosage-dependent effects^15^. Hormone inducible systems have even been expanded to build multi-gene control, demonstrating simultaneous control of multiple enzymes in a pathway ^16^. But all these methods have significant drawbacks. They rely on eukaryotic transcriptional activators fused to DNA binding domains, which can have off-target effects^17^. They also struggle to shut off expression of a gene of interest (GOI) completely, therefore precluding analysis of naturally low-expressed genes. Crucially, they show high cell-to-cell variation and therefore cannot titrate protein levels homogenously across a cell population. This is especially detrimental for applications where only a small number of cells can be observed, such as real-time microscopy. While noise control systems have been engineered to address this issue^18^, they suffer from the same drawbacks detailed above.

To address these drawbacks, we previously developed an inducible system, WTC_846_, that uses prokaryotic repressors rather than eukaryotic transcriptional activators for control, is induced by addition of exogenous anhydrotetracycline (aTc), is completely off in absence of the inducer, and has a very high maximum expression level when fully induced^19^. We also showed that this system is active under different growth conditions without detrimental effects to the cell. Most importantly, WTC_846_ can titrate protein expression much more precisely than all other commonly used inducible systems such as the GAL1 promoter or hormone induced systems. To achieve this, WTC_846_ expresses the GOI from a modified endogenous yeast promoter that is repressible by TetR (P_7tet.1_) within a complex autorepression (cAR) architecture. This architecture is composed of two elements: a simple repression (SR) component where constitutively expressed TetR-Tup1 protein represses P_7tet.1_, and an autorepression component (AR) where TetR is also expressed from P_7tet.1_ and therefore represses its own expression. TetR-Tup1 is a fusion protein between the bacterial TetR and the yeast transcriptional repressor Tup1 and eliminates all basal expression in absence of the inducer aTc. It is weakly expressed, such that upon addition of a small amount of inducer, almost all TetR-Tup1 is inactivated by aTc and the AR component takes over control of GOI expression. The advantage of the AR component is that it expands the titratable range of the promoter, flattening the slope of the typically very steep, sigmoid dose response curves. Therefore, with WTC_846_, intermediate concentrations of protein expression are easily accessible to the researcher. Additionally, the AR component reduces cell-to-cell variation at any given expression level, allowing precise and uniform protein expression across the cell population.

Here, we extend the WTC system to two additional orthogonal bacterial DNA binding proteins and cognate inducers, such that precise and orthogonal control of three GOIs could be achieved in yeast. We extensively characterized these systems, WTC_847_ and WTC_848_, using dose response experiments and by using them to control various essential and non-essential endogenous genes including low-abundance cell cycle genes, medium-to-high-abundance metabolic enzymes and one highly expressed membrane protein. Finally, to demonstrate the usefulness of controlling multiple GOIs simultaneously, we placed two sets of genes within the same cell cycle signaling pathway under the control of orthogonal WTC systems. We demonstrated that we can precisely and simultaneously control the expression level of up to three genes in yeast, which revealed fine-grained dosage-dependent interactions.

## Results

### Engineering new repressor-promoter pairs as building blocks for Well-Tempered Controller systems

The promoter (P_7tet.1_) driving the expression of the GOI in WTC_846_ was based on the endogenous yeast promoter, P_TDH3_^19^. Placing *tetO_1_* sites within P_TDH3_ had allowed P_7tet.1_ to be regulated by binding and unbinding of the bacterial repressor TetR. We replaced all seven *tetO_1_* sites in P_7tet.1_ with *SulA* operator sites, one of the highest affinity natural binding sites for LexA^20^ (Figure 1A). This promoter, P_7SulA.1_, showed 91% of P_TDH3_ activity (Figure 1B) and could be repressed (strongly but not fully) by the fusion protein LexA-human estrogen receptor hormone binding domain (LexA-hER) ^21^ in presence of β-estradiol (Figure S1).

**Figure 1.**
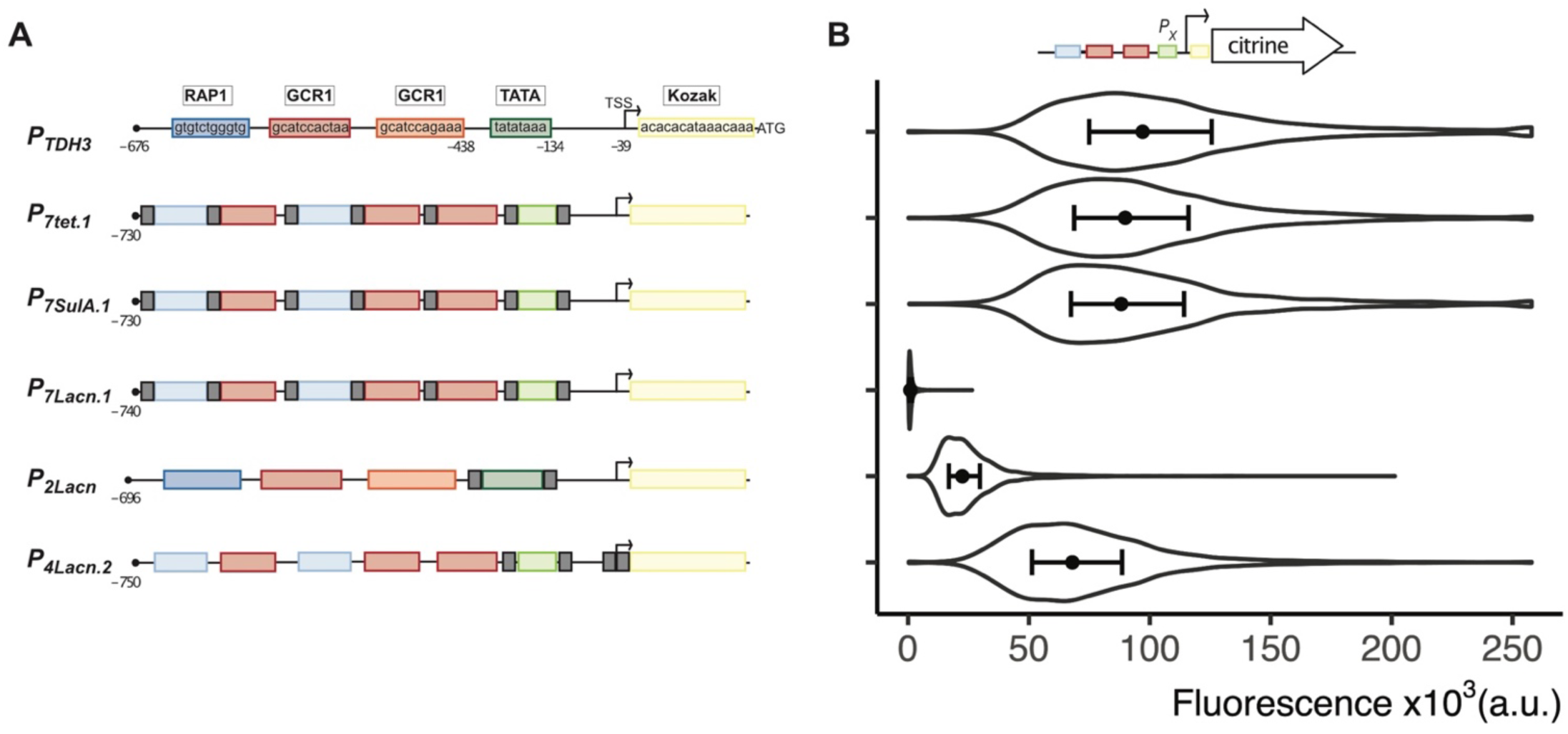
Comparison of maximum activity between the endogenous PTDH3 and derivatives repressed by bacterial proteins. **A)** Schemes depict the promoter designs based on the endogenous PTDH3. Endogenous transcription binding sites, the TATA-box and the Kozak sequence (last 15bp before the start codon) are indicated by colored boxes. The Rap1 binding site and the TATA box have been optimized for increased promoter activity as previously described^19^. These optimized sequences are indicated by lighter color boxes. The grey boxes indicate binding sites for the bacterial protein TetR (in P7tet.1), LexA (in P7SulA.1) or LacI (in P7Lacn.1, P2Lacn and P4Lacn.2). Numbers below the schemes indicate position within the promoter, the first base before the start codon being −1. (TSS= Transcription start site. ATG= start codon). **B)** Maximum promoter activity measured in strains where the promoters depicted in (A) were genomically integrated and drove Citrine expression. The violin plots depict the entire population of exponentially growing cells measured by flow cytometry (n=5000). The dots indicate the median of the population and the error bars span from the first to the third quartile. The strains used were Y3668,Y3728,Y3391,Y3464,Y3412,Y3542, top to bottom.

Various strategies to improve repression of P_7SulA.1_ by LexA-hER, including increasing the size of the repressor, increasing the number of repressor binding sites, and mutating LexA to bind DNA with higher affinity, were unsuccessful. We believe this is due to the low intrinsic transcriptional activation capability of hER in yeast, which could be driving a basal level of promoter activity from P_7SulA.1_ (See Appendix 1) ^22–24^. While repression by LexA-hER is incomplete, repressed activity of P_7SulA.1_ is still extremely low at only 1.3-fold above autofluorescence and 0.54% of P_TDH3_ activity.

Next, we replaced the 7 tetR binding sites in P_7tet.1_ with 7 Lac natural binding sites (*Lacn)* to create P_7Lacn.1_ (Figure 1A). However, unlike P_7tet.1_ and P_7SulA.1_, this promoter had very low activity (Figure 1B). This is likely because *Lacn* sites are longer (21bp) than both *tetO* (15bp) and *SulA* (16bp) sites. This changes the distance between the endogenous binding sites, potentially interfering with the way they interact on the promoter. An alternative design, where there were only 2 *Lacn* sites flanking the TATA-box (P_2Lacn_) and none in the upstream activating region (UAS), showed better maximum activity (Figure 1B).

Repression of this promoter was weak (7.1-fold above autofluorescence control). Replacing *Lacn* sites with higher affinity *Lacs* sites slightly improved repression, but completely abolished induction by IPTG (Figure S2). Further optimization of P_2Lacn_ yielded P_4Lacn.2_, which has two additional *Lacn* sites flanking the Transcription Start Site (TSS), and in which the 2bp on both sides of the TATA-box were reverted to the endogenous P_TDH3_ sequence to increase maximum activity (Figure 1A). This promoter shows a maximum activity that is 70% of the endogenous P_TDH3_ promoter, and a basal level of 4.1-fold above autofluorescence (Figure 1B & Appendix 2). We also further optimized LacI (Apendix 3) for increased repression using six individual amino acid mutations and by adding a Nuclear Localization Sequence (NLS) to create LacI^m^. This optimized LacI repressed P_4Lacn.2_ activity fully and was still inducible to 93% of unrepressed promoter activity in presence of 10 µM IPTG.

### Building the β-estradiol-regulated WTC_847_ and IPTG-regulated WTC_848_

The original WTC_846_ was built with a Complex Autorepression architecture (cAR), which is a combination of Autorepression (AR) to increase titratable range and reduce cell-to-cell variation in expression, and Simple Repression (SR) to reduce basal expression (Figure S3)^19^. For the effects of the AR to be observable, its repressor needs to respond to the inducer at a different concentration range than the SR repressor. Therefore, to implement this architecture with LexA-hER, we needed to shift the dose response curve of the constitutively expressed SR repressor to the right, such that it yielded repression only at the high β-estradiol concentrations where the AR repressor reaches its repression limit. To achieve this, we introduced single amino acid mutations into hER that had been described to reduce its affinity to β-estradiol (A350M,L346I,M388Q,G521S,F461L,V560M,E353Q,F461I,L387M)^25–27^ (Figure S4A). We picked 5 candidates that shifted the dose response to the right compared to the original LexA-hER and checked their repression strength (Figure S4B). We picked the one with the lowest basal level (LexA-hER(L387M), 1.6-fold above autofluorescence) as the SR repressor.

LexA-based cAR was built using P_7SulA.1__citrine and P_ACT1__LexA-hER(L387M)_P_7SulA.1__LexA-hER-adh1tail. The adh1 tail (amino acids 305-348 of the ADH1 protein^28^) serves as a degradation tag to prevent excessive LexA-hER production, which could result in gratuitous binding to DNA. We quantified input dynamic range and Volume Independent Variation (ViV) as described previously^19^. Briefly, input dynamic range is the range of usable doses (where the slope of the dose response is non-zero) and ViV is akin to a cell-size corrected coefficient of variation, used to quantify cell-to-cell variation in a population. Compared to SR, cAR showed larger dynamic range and lower ViV throughout most of this range (Figure 2). Additionally, compared to AR, cAR had lower basal expression (2.6-fold vs. 14.3-fold over autofluorescence) at full repression with 30µM β-estradiol. We call this final cAR design the Well-tempered Controller 847 (WTC_847_).

**Figure 2.**
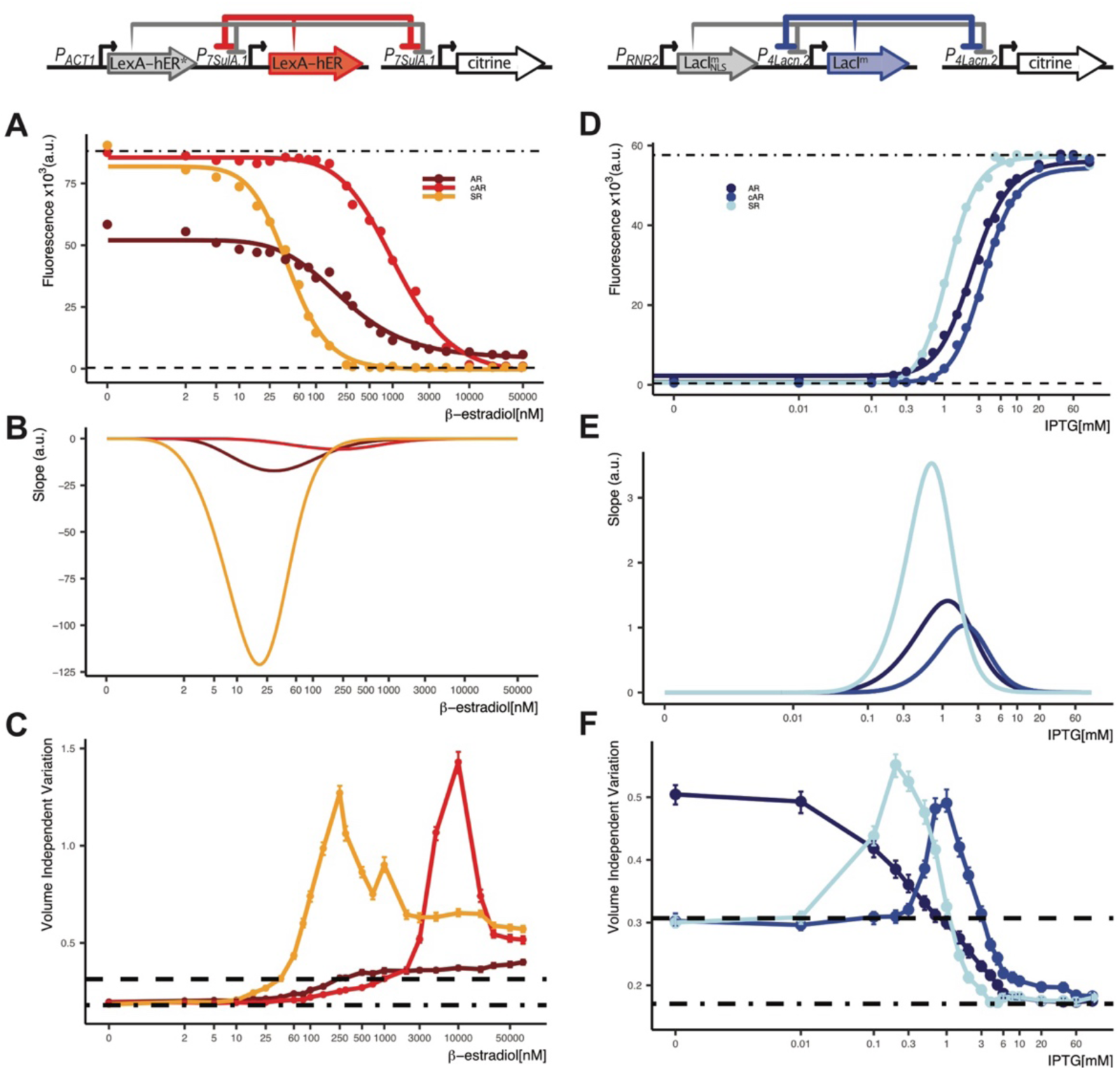
Dose response behavior and cell-to-cell variation of WTC847 and WTC848. **A)** β-estradiol dose response of three strains with varying architectures. A Simple Repression (Y3516), an Autorepression (Y3572) and a strain bearing the WTC847 system (cAR,Y3808) were grown overnight in different β-estradiol concentrations and diluted into the same concentrations in the morning. The scheme above depicts the contents of WTC847. Citrine fluorescence was measured with flow cytometry after 7 hours (n=5000 cells per dose). Symbols indicate the median fluorescence of the measured population at each dose. Lines are fitted using a five-parameter log logistic function as explained in Methods. Dashed line indicates autofluorescence signal measured from the parental strain without Citrine (Y70). Dot-dashed line indicates fluorescence measured from a strain where P7SulA.1 drove Citrine expression (Y3391). **B)** Slopes of the dose response curves shown in (A). **C)** Cell-to-cell variation of expression in the strains shown in (A). Higher Volume Independent Variation values (y axis) mean larger cell-to-cell variation, calculated as explained in the text and previously^19^. Dot-dash line indicates the variation of the strain where Citrine is constitutively expressed from P7SulA.1 and dashed line indicates variation of autofluorescence in the parent strain without Citrine. Error bars indicate 95% confidence interval calculated using bootstrapping (n=1000) as described in Materials and methods. **D)** IPTG dose response of three strains with varying architectures. A Simple Repression (Y3831), an Autorepression (Y3674) and a strain bearing the WTC848 system (cAR, Y3856) were grown overnight in different IPTG concentrations and diluted into the same concentrations in the morning. The scheme above depicts the contents of WTC848. Citrine fluorescence was measured with flow cytometry after 7 hours (n=5000 cells per dose). Symbols and lines as in (A). Dashed line indicates autofluorescence signal measured from the parental strain without Citrine (Y3316). Dot-dashed line indicates fluorescence measured from a strain where P4Lacn.2 drove Citrine expression (Y3542). **E)** Slopes of the dose response curves shown in (D). **F)** Cell-to-cell variation of expression in the strains shown in (D). Higher Volume Independent Variation values (y axis) correspond to greater variation in expression, calculated as explained in the text and previously^19^. Dot-dash line indicates the variation of the strain where Citrine is constitutively expressed from P4Lacn.2 and dashed line indicates variation of autofluorescence in the parent strain without Citrine (Y3316). Error bars indicate 95% confidence interval calculated using bootstrapping (n=1000) as described in Materials and methods.

LacI-based cAR was built using P_ACT1_ expressed Lac12 (for importing IPTG into the cell), P_RNR2_ expressed LacI^m^, P_4Lacn.2_ expressed LacI^m^ and P_4Lacn.2_ expressed Citrine. It showed a wider dynamic range compared to the SR strain (600 vs. 100-fold), and lower cell-to-cell variation (Figure 2). However, the reduction in cell-to-cell variation was not as pronounced as in the other two WTC systems, likely due to the larger overlap between the dose response curve of the SR and AR components. Additionally, expression of the SR and AR constructs from the same loci resulted in a very small but detectable basal expression in the WTC_848_ system of 1.2-fold above autofluorescence. Nevertheless, given the low level of basal expression in this strain, these constructs were chosen to finalize the WTC_848_ system.

### Detailed characterization and orthogonality of WTC systems

We performed detailed characterizations of the new transcriptional control systems. Time dependent dose responses showed that both systems are activated and can be shutoff within 30 minutes of inducer addition/removal (Figure S5). Steady state in expression is reached within 6 to 7 hours for both systems. Both systems are functional and show the same qualitative cell-to-cell variation behavior when various carbon and nitrogen sources are used for growth, though the required inducer concentration changes based on the condition (Figure S6). In terms of cell physiology, we observed that cells bearing WTC_847_ or WTC_848_ grew just as well as the parent control in all media, with or without inducer, except when ethanol was the sole carbon source, where some perturbance to physiology was observed (Figure S7). Overall, like the original WTC_846_, both systems are largely functional under conditions of various carbon and nitrogen sources, making them ideal systems for exploring gene functionality.

We then checked whether all three WTC systems could function in the same cell orthogonally using dose responses (aTc, IPTG, β-estradiol) (Figure S8). All three systems had the same maximum and basal expression levels, and a dose response behavior consistent with previous characterization done in Figure 2 and ^19^. We also confirmed that simultaneous operation of all three systems does not impair growth of cells, compared to the parent strain (Figure S9). This is true for when all three systems are in the OFF state (no aTc, high β-estradiol and no IPTG) or when all three are in the ON state and are expressing as much protein as they can (high aTc, no β-estradiol, high IPTG). Overall, we confirmed that the three WTC systems can perform orthogonal gene expression control without impairing cell physiology.

### Orthogonal two-gene control using the WTC systems recapitulates known dosage interactions

We first confirmed that the two new systems, WTC_847_ and WTC_848_, could be used to control the expression level of endogenous yeast genes on their own (Appendix 4). We explain how to generate such strains in detail in Appendix 5. Since these systems showed the expected behavior when controlling various endogenous genes individually, we moved on to multi-gene control.

We focused on cell cycle cascades centered around the phosphatase Cdc14 (Figure 3A). Cdc14 is localized in the nucleolus throughout most of the cell cycle and is kept there through its interaction with its inhibitor Net1. It is released from this inhibition in two waves triggered by two complementary, successive pathways: first by the FEAR (Cdc14 early anaphase release) and later the MEN (Mitotic Exit Network)^29,30^. Two important inhibition interactions play a central role in these cascades: that of Net1-Cdc14 and Pds1-Esp1. It has been shown that a dosage imbalance between Net1 and Cdc14 is not tolerated by the yeast cell, leading to reduced growth rates^5^. Whereas an imbalance between Pds1-Esp1 is well-tolerated because the dosage imbalance in case of the latter pair could be masked by Cdh1.

**Figure 3:**
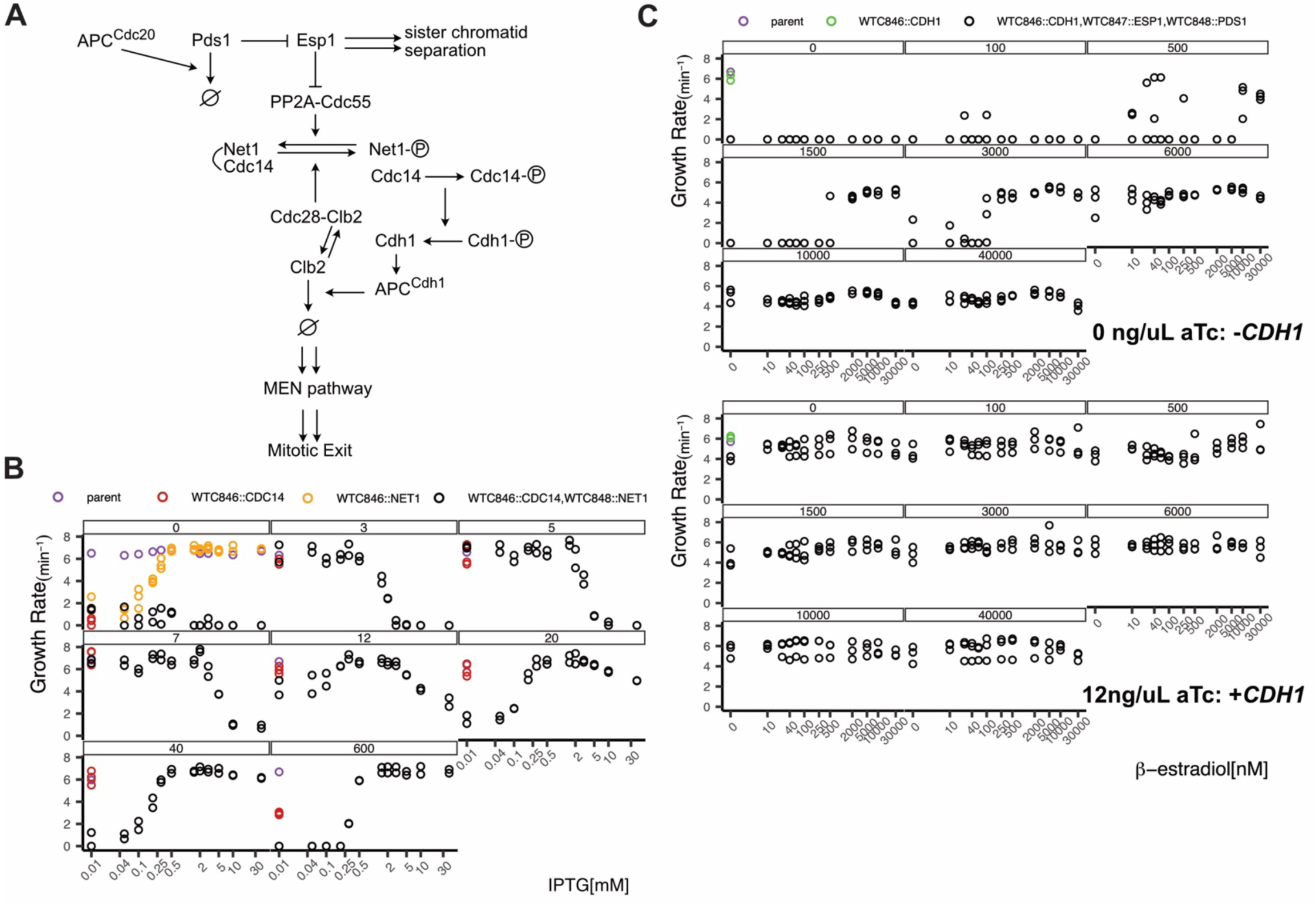
Dosage sensitive interactions within the anaphase regulatory networks. **A)** The FEAR cascade is initiated at anaphase onset when APC^Cdc^^20^ (anaphase promoting complex) marks the Esp1 inhibitor Pds1 (also called securin) for destruction. This allows Esp1 (also called separase) to separate sister chromatids, and to inhibit PP2A-Cdc55 phosphatase activity. This allows increased Net1 phosphorylation, releasing its inhibition on Cdc14. Once released, Cdc14 phosphatase dephosphorylates Chd1. Cdh1 then complexes with APC and degrades mitotic cyclins like Clb2 to lower Cdc28 activity. Lowered Cdc28 activity allows the MEN pathway to be triggered, causing sustained release of Cdc14. Overall, Cdc14 activity throughout late M phase drives correct cell cycle progression, and the two separate waves of release ensure that the sister chromatid separation occurs before mitotic exit. **B)** All cells (Y3878,3925,3932,3945) were grown overnight with low (5ng/mL) aTc. *CDC14^846^NET1^8^*^48^ cells were additionally provided with 10µM IPTG. In the morning cells were spun down and washed twice with YPD, followed by 6 hours of growth in YPD without inducers. Cells were then diluted again into plate reader plates with varying amounts of inducer. Each facet corresponds to a different aTc concentration in ng/µL. Maximum growth rate observed during the exponential phase is used as a proxy for fitness. Each dose is in duplicates except for the parent strain, and each circle represents an independently growing culture. **C)** Same as in B, except cells (Y3878,3928,3979) were grown with 12ng/µL aTc overnight and *CDH1^846^ESP1^847^PDS1^848^*cells were additionally given 30 µM β-estradiol. Each facet corresponds to one IPTG concentration in µM. Top panel has no aTc and thus Cdh1 expression is turned off. Lower panel has aTc and thus expressed Cdh1 at a level that allows normal cell growth. Each dose is in triplicates except for the parent strain, and each circle represents an independently growing culture.

We placed each gene of interest under the control of an orthogonal WTC system and recorded the growth of these cells in liquid culture under different inducer concentrations in rich media, and quantified maximum growth rate observed during exponential phase as a proxy for the fitness of the cell. As expected, we saw that cells did not grow in absence of the essential gene *CDC14* (Figure 3B and S10A). And in the double gene control strain (*CDC14*^846^*NET1*^848^) we observed that when Cdc14 expression was low, high levels of Net1 led to a reduction in fitness, and vice versa, confirming the requirement for balanced dosage of these two proteins for optimal cell growth.

### High lethality threshold of CDH1 overexpression

To investigate dosage masking by Cdh1, we first titrated Cdh1 expression alone to determine the expression level that allowed for optimal growth in a *CDH1*^846^ cell (Figure 4A and S11A). We observed that extreme overexpression of *CDH1*, which was previously shown to cause cell cycle arrest in M phase and lethality^31^, prevents growth. Interestingly, we noticed that significant overexpression of Cdh1 is well tolerated up until a certain point, and no decrease in fitness is observed even with 50% activity (100ng/µL aTc) of WTC_846_ (which corresponds to roughly 400.000 to 500.000 molecules per cell, and endogenous Cdh1 levels are estimated to be <1000 ^32^). At about 83% of WTC_846_ activity (200ng/mL, ∼ 800.000 molecules per cell), fitness is minimally reduced and only an increased lag phase is observable (Figure S11). At full WTC_846_ activity no growth was observed. One potential explanation for this overexpression tolerance followed by sudden lethality is competition between Cdh1 and Clb2. It is known that Clb2 cyclin can induce its own expression, therefore activating a positive feedback loop that drives cells through the G2/M phase^33^. If Cdh1 levels are too high, this positive feedback loop would not be activated. But if there is just enough Clb2, the positive feedback loop would amplify its expression enough to allow the cell to progress through the cell cycle.

**Figure 4:**
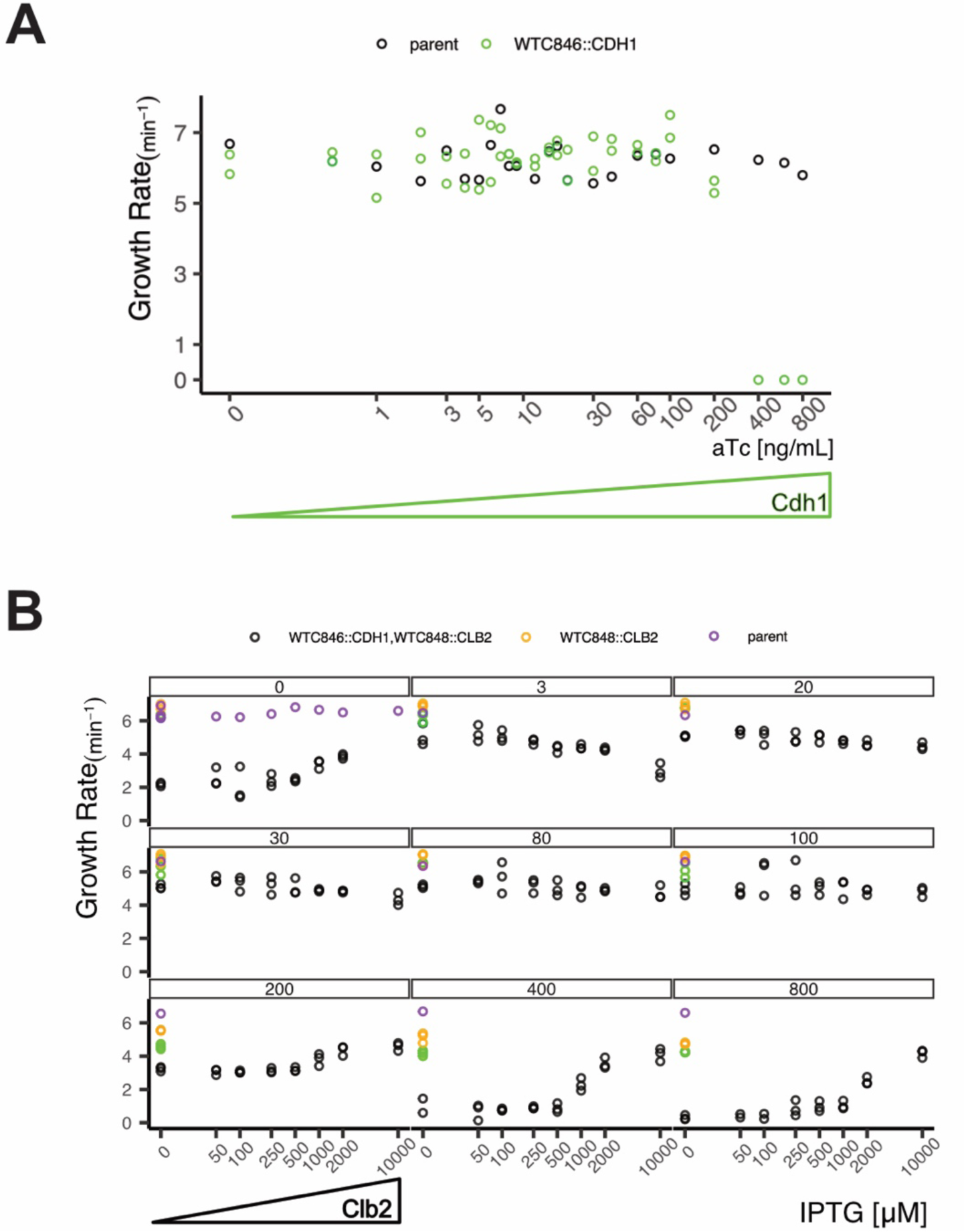
Cdh1 interacts with Clb2 in a dosage dependent manner. **A)** *CDH1^846^* (Y3928) strain was used to titrate increasing amounts of Cdh1. Growth rates were observed using a plate reader. Maximum slopes during exponential phase were quantified as a proxy for fitness. Each dose is in duplicates for the *CDH1^846^* strain and each circle represents an independently growing culture. **B)** Same as in (A), performed with parent (Y3878), *CDH1^846^* (Y3928, in duplicates), *CLB2^848^* (3923, in duplicates), *CDH1^846^CLB2^848^* (4043, in triplicates) strains. Each facet represents one aTc concentration in ng/µL (controlling Cdh1 expression) and IPTG concentration corresponds to Clb2 expression level. Each circle represents an independently growing culture.

As a preliminary test of this hypothesis, we titrated Cdh1 and Clb2 levels the same cell (*CDH1^8^*^46^ *CLB2^848^*). Overexpression of Clb2 could rescue the lethality associated with Cdh1 overexpression (Figure 4B and S11B), and we also noticed that *CDH1^846^* cells in which Cdh1 was overexpressed eventually grew at around 2/3 the speed of parent cells, albeit with an incredibly long lag phase (Figure S11B). This could indicate adaptation, perhaps through accumulation of Clb2 and subsequent activation of the positive feedback loop. On the other hand, *CDH1^846^ CLB2^848^* cells where Clb2 expression was suppressed failed to show this adaptation. Therefore, these results suggest that Cdh1 and Clb2 interact in a dose-dependent manner, and Cdh1 overexpression lethality is linked to Clb2. Although further investigation, including quantification of actual protein levels would be needed in the future to confirm this hypothesis, this serves as a demonstration that precise titration of gene products with WTC systems can uncover previously unknown dosage-dependent gene-gene interactions.

### Orthogonal three-gene control with WTC systems

Finally, we measured fitness of the triple control strain (*CDH1^846^ESP1^847^PDS1^848^*) to determine whether we observed dosage imbalance masking by *CDH1*. We varied Esp1 and Pds1 levels when Cdh1 was absent (no aTc) or present (12 ng/mL aTc) (Figure 3C). We observed that when Cdh1 was present, fitness was stable despite varying Pds1 or Esp1 levels. On the other hand, when Cdh1 expression was completely turned off, cells did not grow at all when Esp1 expression was high and Pds1 level was low. Only with increasing Pds1 concentrations could cells expressing high Esp1 grow, supporting the idea that a dosage imbalance between Esp1 and its inhibitor Pds1 can be masked by Cdh1. As explained by the authors of the original Tug-of-War study^13^, one potential mechanism for this masking is through Cdc20 stabilization. Since Cdc20 is a potential target of the APC^Cdh1^ complex for degradation, and APC^Cdc20^ activity can lead to degradation of Pds1, lack of Cdh1 might reduce the amount of Pds1 available, widening the gap between Esp1 and Pds1 expression levels.

Interestingly, the dosage imbalance was seen only when Esp1 was overexpressed in the triple control strain (*CDH1^846^ESP1^847^PDS1^848^*), and not when Pds1 was overexpressed. We therefore checked the fitness of the double control strain *CDH1^846^PDS1^848^*, where Esp1 is expressed under endogenous control. In this case, overexpressing Pds1 did cause a loss of viability without Cdh1 (Figure S12). This was probably not observed in the *CDH1^846^ESP1^847^PDS1^848^* strain due to basal leakiness of the WTC_847_ system combined with the lower maximum activity of the WTC_848_ system, highlighting the importance of precise titration of expression over large ranges for uncovering gene-gene interactions. Although protein levels would need to be quantified to confirm this, these results indicate that Cdh1 masks the dosage imbalance between Esp1 and Pds1 both when Esp1 is overexpressed and when Pds1 is overexpressed. However, the APC^Cdc20^ inactivation theory explained above would not explain why Pds1 overexpression can be masked by Cdh1. It is possible that Pds1 overexpression delays sister chromatin separation, while lack of Cdh1 leads to easier accumulation of Clb2 and thus higher Cdc28 activity, causing early activation of the MEN pathway. This temporal disruption (MEN activation before proper sister chromatid separation) could be leading to reduced fitness in the cell. While this was not further explored here, these results demonstrate clearly that precise multi-gene control using the WTC systems can be a powerful tool for hypothesis generation when investigating gene-gene interactions.

## Discussion

Inducible control of endogenous genes is a fundamental technique used in biological experimentation. Changing the expression level of proteins provides information on their function, interactions with other proteins and potential regulatory relationships between them. Previously we had developed a new inducible system called WTC_846_, which we showed was more precise and capable of a larger expression range than any other inducible system. The key components were a promoter repressible by TetR and a complex autorepression (cAR) architecture. Here we demonstrated that the promoter design and the cAR architecture used for WTC_846_ are in principle generalizable but requires optimization for each new repressor. We then demonstrated that these new systems are functional both alone and orthogonally together within the same cell. We generated strains where two or three interacting genes were orthogonally controlled and reproduced previously observed dosage-dependent interactions. We also made observations suggesting that the Cdh1 overexpression lethality can be rescued by overexpression of Clb2. Overall, we showed the utility of precisely titrating more than one gene with WTC systems to gain insight into gene and pathway functions in *S. cerevisiae*.

The three systems show differences that might make one system more suitable for a certain application than others. The original WTC_846_ system can fully suppress expression of any gene placed under its control, whereas the other two systems have a certain level of basal expression even under conditions where gene expression is uninduced. WTC_848_ has a lower basal expression than WTC_847_, evidenced by the fact that WTC_848_ suppression of genes such as *CDC20* lead to a larger growth impairment compared to WTC_847_ based suppression. On the other hand, both WTC_846_ and WTC_847_ can reach maximum expression levels on par with the *TDH3* promoter, estimated to be one of the most active promoters in yeast^32^. At full induction, WTC_848_ reaches about only 60% of this level. Therefore, WTC_846_ is most suitable for experiments where gene expression needs to be reversibly turned off. If complete lack of the gene product is not vital, WTC_848_ is also suitable for expressing endogenously low-expressed genes. While all systems do reach a high maximum level, overexpression studies would benefit most from using the WTC_846_ and WTC_847_ systems. Additionally, it is advisable to use WTC_846_ and WTC_847_ in cases where precise and uniform expression of a gene is desired within the cell population. An example might be microscopy setups, where only a small number of cells can be observed over time, and a homogenous population would be conducive to drawing strong conclusions.

Thus far in the first three iterations of the WTC design, we used very well-characterized prokaryotic repressors. TetR, LexA and LacI have been studied for decades, but there are many other prokaryotic repressors that are similarly well-characterized. For example, other TetR family repressors and their cognate ligands have been shown to be orthogonal in an activation-based system in yeast^34^. Recently, 12 different repressor-inducer pairs have been characterized and optimized for low basal leakiness, high activity and low cross-talk in E.coli^35^. For at least some of these repressors, there are also detailed studies on how the placement of their binding sites within yeast promoters affect their repression efficiency^36^. These all represent potential starting points for future WTC developments to expand multi-gene control.

Given their orthogonality, the WTC systems are likely to find use in multi-gene studies where pathway interactions are being investigated. For example, investigating the reasons behind the lethality of *CDH1* overexpression might benefit from simultaneous titration of Cdh1 and a non-degradable Clb2 version^37^, or Cdh1 and Cdc20 to see if the competition for the APC complex between them plays a role. Additionally, there are further 114 genes originally determined to be dosage sensitive by the same Tug-of-War study that identified the Net1-Cdc14 dosage sensitivity^12^. Some of these have been shown to have partners that compensate for their overexpression. These gene pairs would represent a good starting point for further study using the WTC systems. Additional readouts, such as quantification of protein levels, can be used to determine the exact threshold of balance to imbalance, and might shed light on compensation mechanisms that prevent dosage imbalance. Furthermore, large-scale inducible gene libraries could also be constructed with WTC systems to hunt for further interacting partners. While technically challenging, crossing of two or even three such libraries would create a very detailed map of dosage-dependent gene interactions and yield hitherto inaccessible knowledge on gene networks that could potentially be extrapolated to mammalian systems.

## APPENDICES

### Appendix 1: Strategies to increase repression of P7SulA.1 by LexA-hER

We had four strategies to address the basal leakiness issue, based on different hypotheses of why repression of P_7SulA.1_ by LexA-hER was incomplete. Initially, we hypothesized that LexA-hER was occluding endogenous transcription factor binding through steric hindrance but was either not large enough or wasn’t binding strongly enough to the *SulA* sites to compete effectively with the endogenous factors. We increased repressor size by fusing it to the *E.coli* Maltose Binding Protein (MBP) and we increased DNA binding affinity by using mutant LexA versions (V59I, E71K) known to bind operator DNA with higher affinity^38^. We created strains where the LexA repressible promoter drove Citrine expression and the repressors being tested were driven by P_ACT1_ (Y3402, 3438,3441,3442). Compared to the original strain with P_7SulA.1_ and LexA-hER (Y3408), none of these repressors showed better repression (Figure Appendix 1A).

**Figure Appendix 1.**
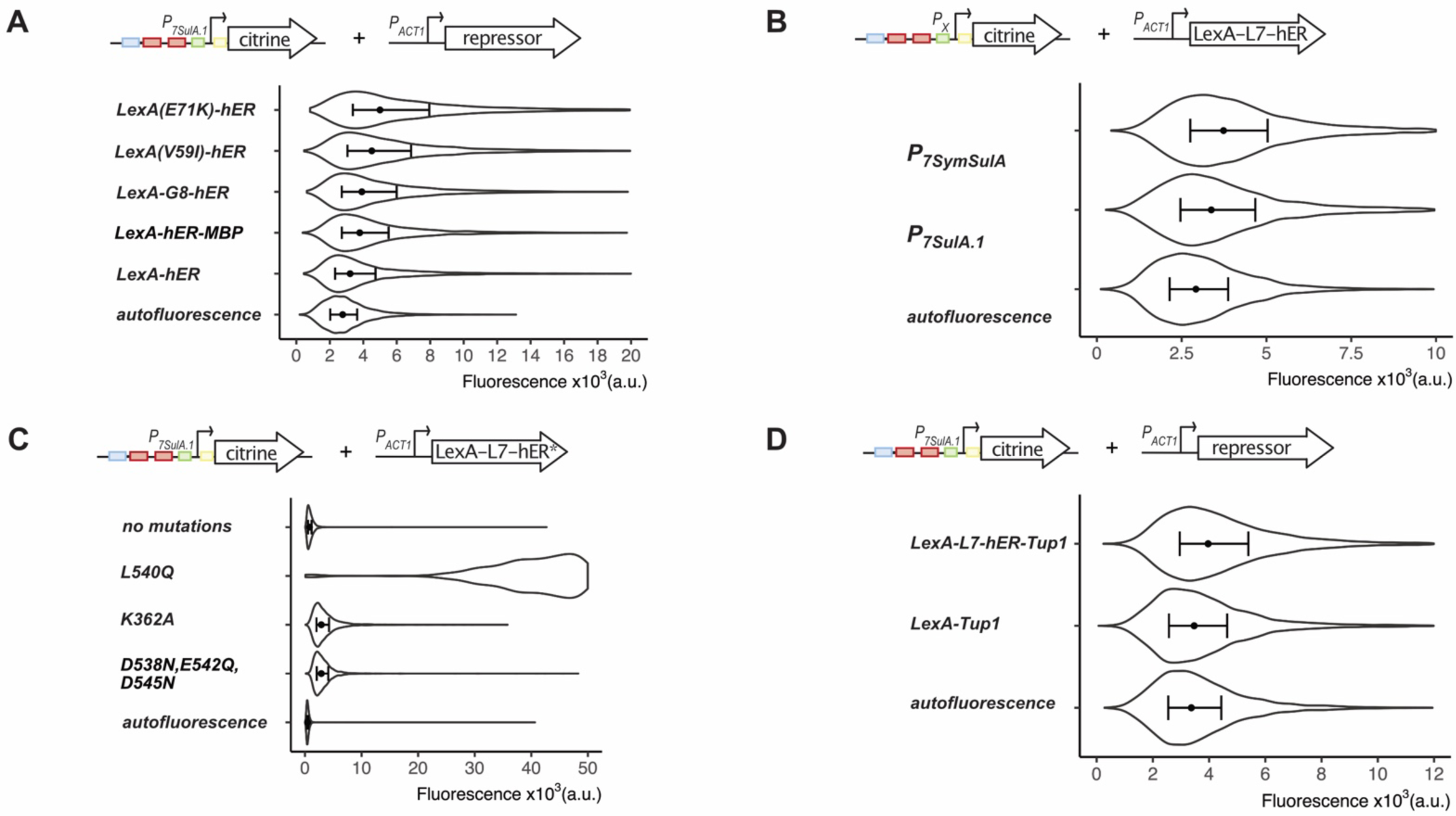
Strategies to increase repression by LexA-hER. **A)** Strains where different versions of the repressor LexA-hER and the construct P7SulA.1_Citrine were genomically integrated were grown overnight in presence or absence of 10µM β-estradiol (Y3402,3408,3438,3441,3442). Strains were diluted into the same conditions and Citrine fluorescence was measured using flow cytometry during exponential phase (n=5000 cells). Fluorescence was compared to the parent strain (Y70) as autofluorescence control to determine basal level. G8= 8 glycine residues, MBP= Maltose Binding Protein. Symbols inside the violin plots indicate the median of the population, and the error bars span Q1-Q3. **B)** Two versions of the LexA-hER repressible promoter (P7Sym and P7SulA.1) were genomically integrated and used to drive Citrine expression in a strain where PACT1 drove the expression of LexA-L7-hER (Y3516,3620). Citrine fluorescence was measured and plotted as in (A). **C)** Different mutations were introduced into the hER portion of the repressor LexA-hER. These repressor versions were integrated into a strain where P7SulA.1 drove Citrine expression (Y3516,3664,3651,3642). Citrine fluorescence was measured and plotted as in (A), but with 30µM β-estradiol instead of 10µM. The strain with the mutation L540Q had a very high expression level, and therefore the violin plot is cut off. **D)** Two strains where either LexA-Tup1 or LexA-L7-hER-Tup1 were expressed from PACT1 and Citrine from P7SulA.1 were created (Y3719,3849). Citrine fluorescence was measured and plotted as in (A).

Next, we hypothesized that the short homology (CATCCA) between the endogenous transcription factor Gcr1 binding site and the *SulA* operator might be causing competition at the *SulA* site between Gcr1 and LexA-hER binding. We therefore changed *SulA* operators in the UAS of P_7SulA.1_ with *SymLexA*, which are symmetrical consensus LexA binding sites, and tested this promoter (P_7SymSulA_) in the same fashion as above. However, this modification led to a decrease in repression strength instead, likely because LexA has a higher affinity towards *SulA* than *SymLexA* sites (Figure Appendix 1B, Y3620, 3622, 3516).

Finally, we hypothesized that the ligand binding domain of hER, containing transcriptional activation domains functional in human cells, could be driving a small amount of transcription in yeast from P_7SulA.1_. The full size hER had been shown to drive transcription in yeast from synthetic promoters with hER binding sites inserted^22–24^. To reduce this intrinsic activity, we created 3 different versions of LexA-hER, where we incorporated one or more mutations into hER that were described to abolish its transcriptional activation ability (K362A,L540Q, or a combination of D538N,E542Q and D545N) ^24,39–42^. Tested in the same fashion described above, these mutations again decreased repression strength instead of increasing it (Figure Appendix 1C). However, being within the hormone binding domain, the mutations might have affected transcriptional activation ability and hormone binding ability simultaneously. Therefore, we created two other constructs to check whether the promoter could be fully repressed. We fused LexA-L7-hER and only LexA to the yeast transcriptional repressor Tup1, which when fused to TetR could suppress all activity from the analogous promoter^19^. Whereas the LexA-Tup1 construct could suppress all expression of P_7SulA.1_, the LexA-L7-hER-Tup1 construct could not, potentially indicating weak but detectible transcriptional activation by hER (Figure Appendix 1D, Y3651,3642, 3664, 3719). Overall, despite multiple attempts at increasing repression, the best repressor was still the original LexA-hER fusion protein.

### Appendix 2: Optimization of the LacI repressible P2Lacn

As P_7Lacn.1_ had shown, *Lacn* sites in the Upstream Activating Sequence (UAS) were detrimental to functioning of the promoter. Therefore, we chose to place any additional *Lacn* sites to improve repression within the core promoter. We placed one right after putative Transcription Start Site (TSS), which is at the −39^th^ nucleotide in the P_TDH3_^43^. When driving Citrine expression and repressed by P_ACT1_ driven LacI, P_3Lacn.2_ (Y3470) reduced basal level 2.3-fold compared to P_2Lacn_(Y3420) and brought it down to a level of 4.6-fold above autofluorescence (Figure Appendix 2). Before attempting further decreases in the basal level, we added one additional Rap1 and one additional Gcr1 binding site within the UAS to increase maximum level, because this modification had successfully increased the maximum level of the TetR repressible promoter^19^. The resulting promoter, P_3Lacn.3_, had a slightly higher maximum expression level (1.1-fold increase) and a repression level of 4.4-fold above autofluorescence (Y3518,Y3765). To improve repression of this promoter, we added a second *Lacn* site before the Kozak sequence, to generate P_4Lacn.1_. While this modification did reduce basal level to 2.1-fold above autofluorescence, it also decreased maximum level once again. As a final step, we performed three modifications to increase maximum level: we removed either the (i)upstream or the (ii)downstream TATA-flanking *Lacn* site, or (iii) we reverted the 2 upstream and 2 downstream base pairs directly adjacent to the TATA-box back to the endogenous sequence. We call these promoters P_3Lacn.4_,P_3Lacn.5_, and P_4Lacn.2_. The last modification was performed because the sequence of these 2 base pairs was shown to dramatically effect expression level of yeast promoters^44^. While this modification reduced the length of the *Lacn* sites flanking the TATA-box to 19bp, it still kept all the bases shown to be critical for LacI binding^45^. We measured maximal and basal levels of these promoters using flow cytometry. P_4Lacn.2_ showed the second highest maximal level and the lowest repressed level (4.1-fold above autofluorescence) of the three candidate promoters and was therefore chosen as the promoter for the LacI-based WTC_848_ system.

**Figure Appendix 2.**
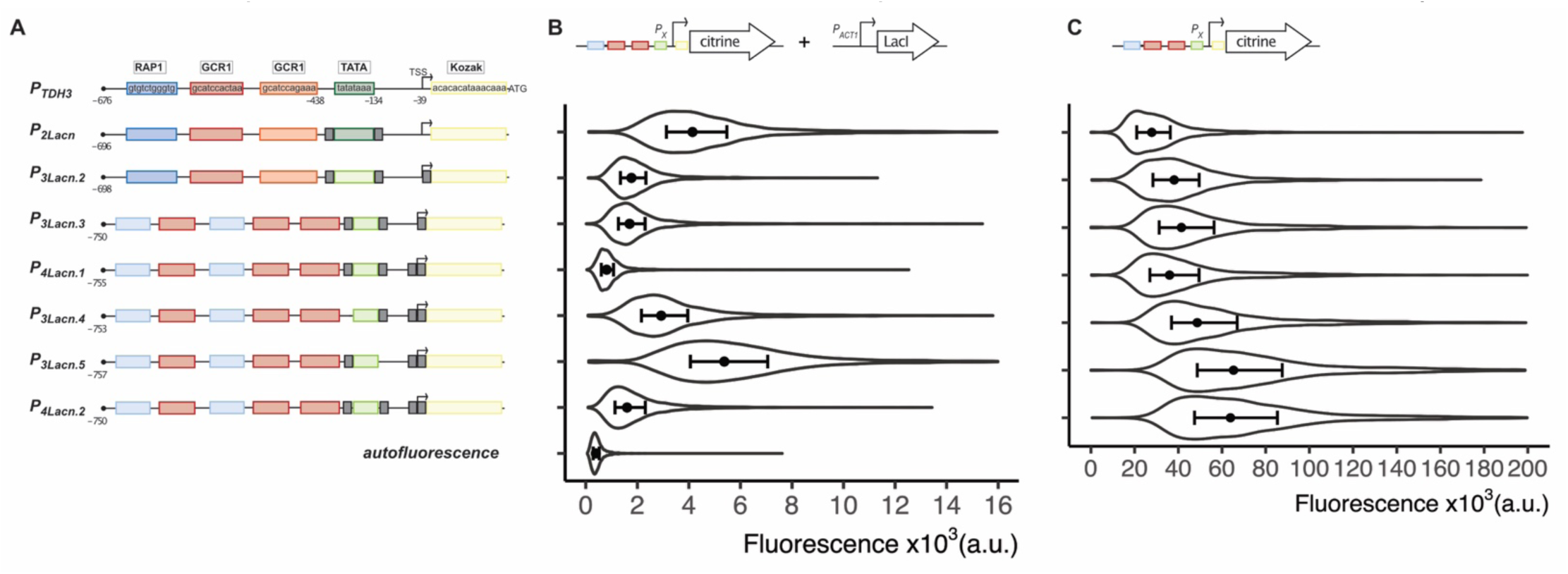
Maximum expression and repressibility of different LacI repressible promoter designs. **A)** Schemes depict the promoter designs based on the endogenous PTDH3. Endogenous transcription binding sites, the TATA-box and the Kozak sequence (last 15bp before the start codon) are indicated by colored boxes. In some designs, the Rap1 binding site and the TATA box have been optimized for increased promoter activity as described in ^19^. These optimized sequences are indicated by lighter color boxes. The grey boxes indicate binding sites for the bacterial protein LacI. Numbers below the schemes indicate position within the promoter, the first base before the start codon being −1. (TSS= Transcription start site. ATG= start codon). **B)** Repressed activity measured in strains where the promoters depicted in (A) were genomically integrated and drove Citrine expression, and the *ACT1* promoter drove expression of LacI. The violin plots depict the entire population of exponentially growing cells measured by flow cytometry (n=5000). The dots indicate the median of the population, and the error bars span from the first to the third quartile. The strains used were Y3420,Y3470,Y3765,Y3498,Y3445,Y3444,Y3547,Y3316, top to bottom. **C)** Maximum promoter activity measured in strains where the promoters depicted in (A) were genomically integrated and drove Citrine expression. Plots as in (B). The strains used were Y3412,Y3463,Y3518, Y3497,Y3540,Y3539,Y3542, top to bottom.

### Appendix 3: Optimization of LacI for better repression of P4Lacn.2

The activity of the optimized promoter, P_4Lacn.2_, could not be fully repressed by wild type LacI. Therefore, the next step was to optimize the repressor for tighter repression, while maintaining inducibility by IPTG. To this end, we initially fused LacI to the yeast endogenous transcription repressor Tup1. This fusion had completely abolished basal expression in the context of the TetR-based system^19^. Here also the P_ACT1_ expressed LacI-Tup1 fusion protein reduced basal expression substantially to 1.1-fold over autofluorescence, when repressing Citrine expression from P_4Lacn.2_(Y3581). However, it also reduced how well IPTG could induce expression from this promoter, such that at full induction with 10mM IPTG, promoter activity was only 20% of the unrepressed P_4Lacn.2_ (Y3542) (Figure Appendix 3.1A). To solve this issue, we introduced single amino acid mutations in the fusion protein LacI-Tup1 that had been described to improve inducibility of LacI (K33E,M242L,T167A,V313A,M254L) (Y3554,3583,3582,3584,3591)^46^. These modifications led to induced activity from the P_4Lacn.2_ promoter that was 87,21,30,21,30% of unrepressed promoter (Figure Appendix 3.1A). However, we noticed a slight (∼1.2 fold compared to unmutated LacI-Tup1) increase in basal level with all these mutations, except for T167A which showed no increase and K33E, which increased basal level by around 30-fold.

**Figure Appendix 3.1.**
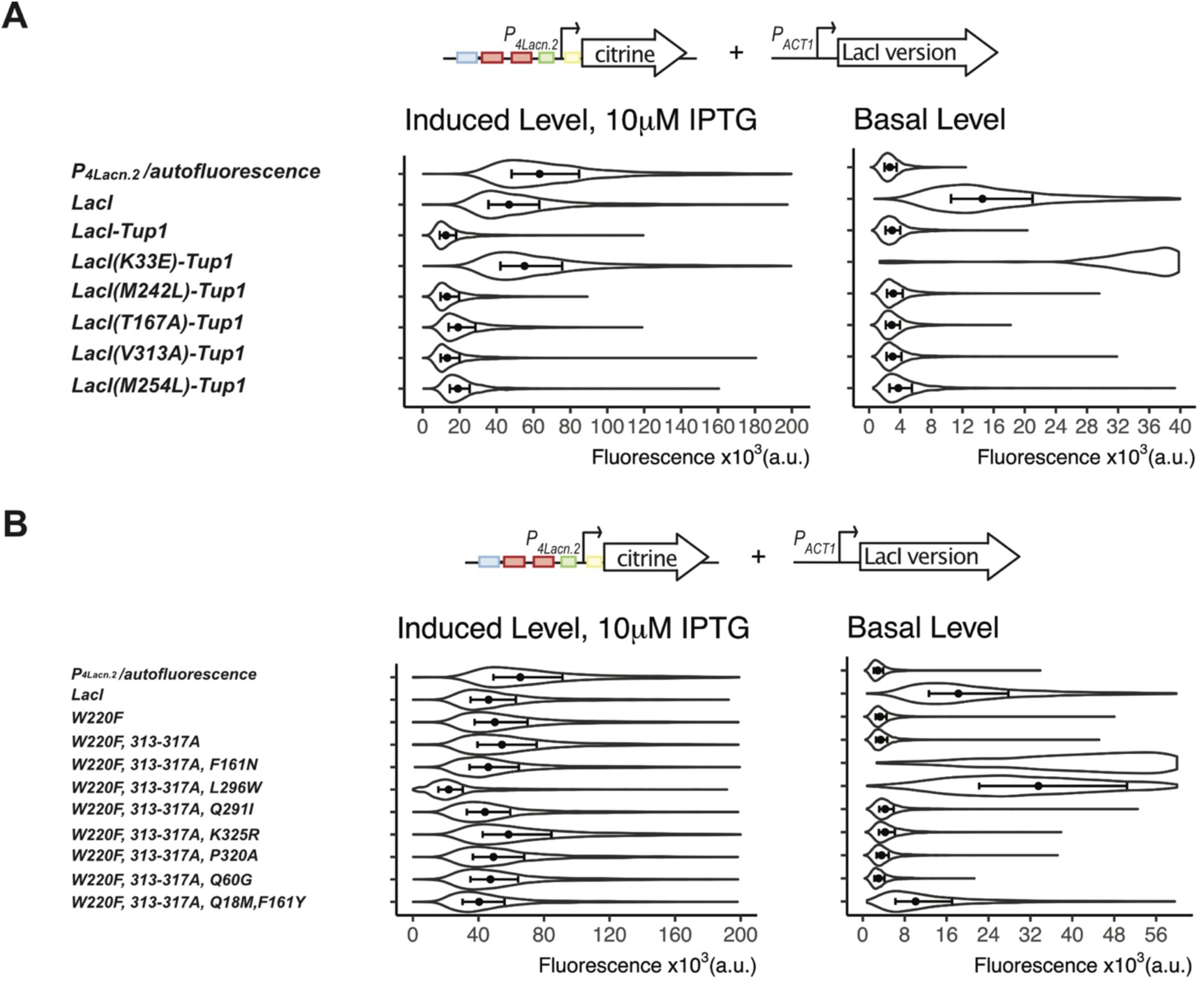
Optimization of LacI for increased repression strength and better inducibility by IPTG. **A)** LacI was fused to the yeast endogenous transcription factor Tup1 and single mutations were introduced into LacI. These various versions of the repressor were genomically integrated and expressed from PACT1. In these strains, the LacI repressible P4Lacn.2 expressed Citrine and PACT1 expressed Lac12. The scheme above the plots indicates the elements genomically integrated into each strain. These strains were cultured with or without 10 µM IPTG overnight and diluted into the same conditions to measure basal and induced activity levels of the promoter in presence of the different repressors. Citrine fluorescence was measured using flow cytometry during exponential growth phase (n=5000 cells per strain). Strain Y3542 bearing only P4Lacn.2 expressed Citrine and strain Y70 bearing no integrated constructs were measured as the P4Lacn.2 and autofluorescence controls. The violin plots depict the entire population within the given x-axis range, the symbol is the measured median fluorescence and the error bars span from the first to the third quartile. The repressor with the mutation K33E had a very high basal level, therefore the violin plot is cut off. For basal level measurements, flow cytometer sensitivity was increased to maximum. **B)** Different LacI versions bearing either single or multiple amino acid mutations were constructed and expressed as explained in (A). 313-317A indicates that all amino acids from 313 to 317 inclusive were mutated to Alanine. Culture conditions, measurements and plots same as in (A).

Therefore, in a second step we tested mutations that had been described to reduce leakiness of LacI repression (W220F, F161N,L296W,Q291I,K325R,P320A,Q60G,Q18M+F161Y,313-317A)^35,47–50^. We tested these either as single mutations or as combinations, in strains where P_ACT1_ expressed LacI variants and P_4Lacn.2_ expressed Citrine (Y3574,3606,3613,3607,3605,3604,3603, 3612,3593). We observed that only two mutations reduced basal leakiness in our context, which were Q60G (1.3-fold reduction) and W220F (5.5-fold reduction) (Figure Appendix 3.1B).

We then combined these two mutations (Q60G+W220F) with some selected inducibility mutations described above (T167A+M242L+M254L) within LacI-Tup1. We later discovered that an additional mutation, V119I, was introduced into this repressor, likely due to an error during PCR-mediated cloning. This repressor with 6 aa mutations was called LacI^m^-Tup1. It was expressed from the *RNR2* promoter, which was the weak promoter we had used for the original WTC_846_ system. We also added two NLS sequences to LacI-Tup1^m^ to increase its nuclear concentration. However, when expressed in this manner, LacI-Tup1^m^ did not repress basal level of P_4Lacn.2_ fully (1.2-fold above autofluorescence), and was only inducible to 64% of unrepressed P_4Lacn.2_ (Figure Appendix 3.2). We then incorporated all 6 mutations described above and an NLS into LacI alone to create LacI^m^, and P_4Lacn.2_ activity was repressed fully in the absence of IPTG, and was still inducible to 93% of unrepressed promoter activity in presence of 10 µM IPTG. Therefore, LacI^m^ with NLS (LacI^m^) was chosen as the final repressor.

**Figure Appendix 3.2.**
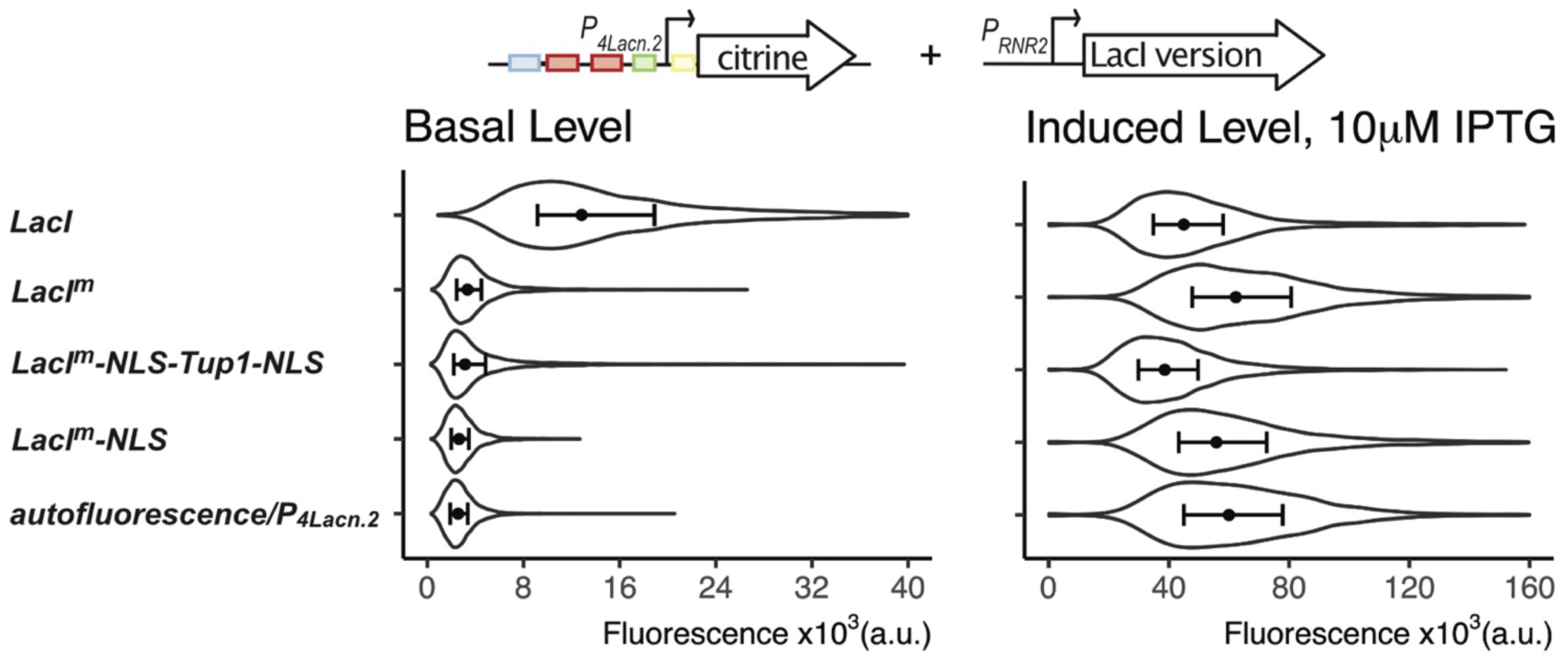
Repression strength and inducibility of the optimized LacI. Several modifications, including amino acid mutations and fusion to Tup1, were performed on the repressor LacI. These repressor versions were genomically integrated and expressed from PRNR2. In these strains, the LacI repressible P4Lacn.2 expressed Citrine and PACT1 expressed Lac12. The scheme above the plots indicates the elements genomically integrated into each strain. LacI^m^ indicates LacI with 6 amino acid mutations (Q60G+W220F+T167A+M242L+M254L+V119I). NLS is nuclear localization signal. These strains were cultured with or without 10 µM IPTG overnight and diluted into the same conditions to measure basal and induced activity levels of the promoter in presence of the different repressors (Y3547,3832,3807,3831). Citrine fluorescence was measured using flow cytometry during exponential growth phase (n=5000 cells per strain). Strain Y3542 bearing only P4Lacn.2 expressed Citrine and strain Y70 bearing no integrated constructs were measured as the P4Lacn.2 and autofluorescence controls. The violin plots depict the entire population, the symbol is the measured median fluorescence and the error bars span from the first to the third quartile.

#### Appendix 4: New WTC systems can regulate gene expression on their own

We placed various low, medium and high expressed genes under the control of P_7SulA.1_ in a strain where WTC_847_ components were integrated (Y3809), or P_4Lacn.2_ in a strain where WTC_848_ components were integrated (Y3870) (Figure Appendix 4). We chose these genes to represent different endogenous expression levels and various functions within the cell, including cell cycle regulators (CDC28,CDC42,CDC20), metabolic enzymes (FBA1,PGK1,TP1), proteins that function in cell wall production (KRE5,KRE9), proteins that regulate sister chromatid separation (ESP1, IPL1), an uncharacterized protein (PBR1), a signaling protein (TOR2), and a cell membrane transporter (PMA1). All except KRE9 were essential genes, and disruption of KRE9 was also shown to lead to reduced growth rates^51^. We performed spotting assays and liquid growth assays with different inducer concentrations. Both systems could regulate the expression level of endogenous genes, as evidenced by the different growth rates observed at different inducer concentrations, both in spotting and liquid assays. Neither system could completely abolish function of the endogenously low-expressed essential genes such as *TOR2,* and therefore cells always grew in presence of high β-estradiol/absence of IPTG. This was expected, given that both these WTC systems displayed a low level of basal expression when controlling fluorescent protein expression as well. We also observed reduced growth rates upon overexpression of certain genes, such as *IPL1, CDC20,* and *TOR2* which are among the lowest expressed proteins tested here (∼1000-2000 molecules per cell^32^). We had previously demonstrated these overexpression phenotypes of using the WTC_846_ system as well. Overall, we confirmed that the new WTC systems could be used to regulate the expression level of endogenous genes on their own.

**Figure Appendix 4.**
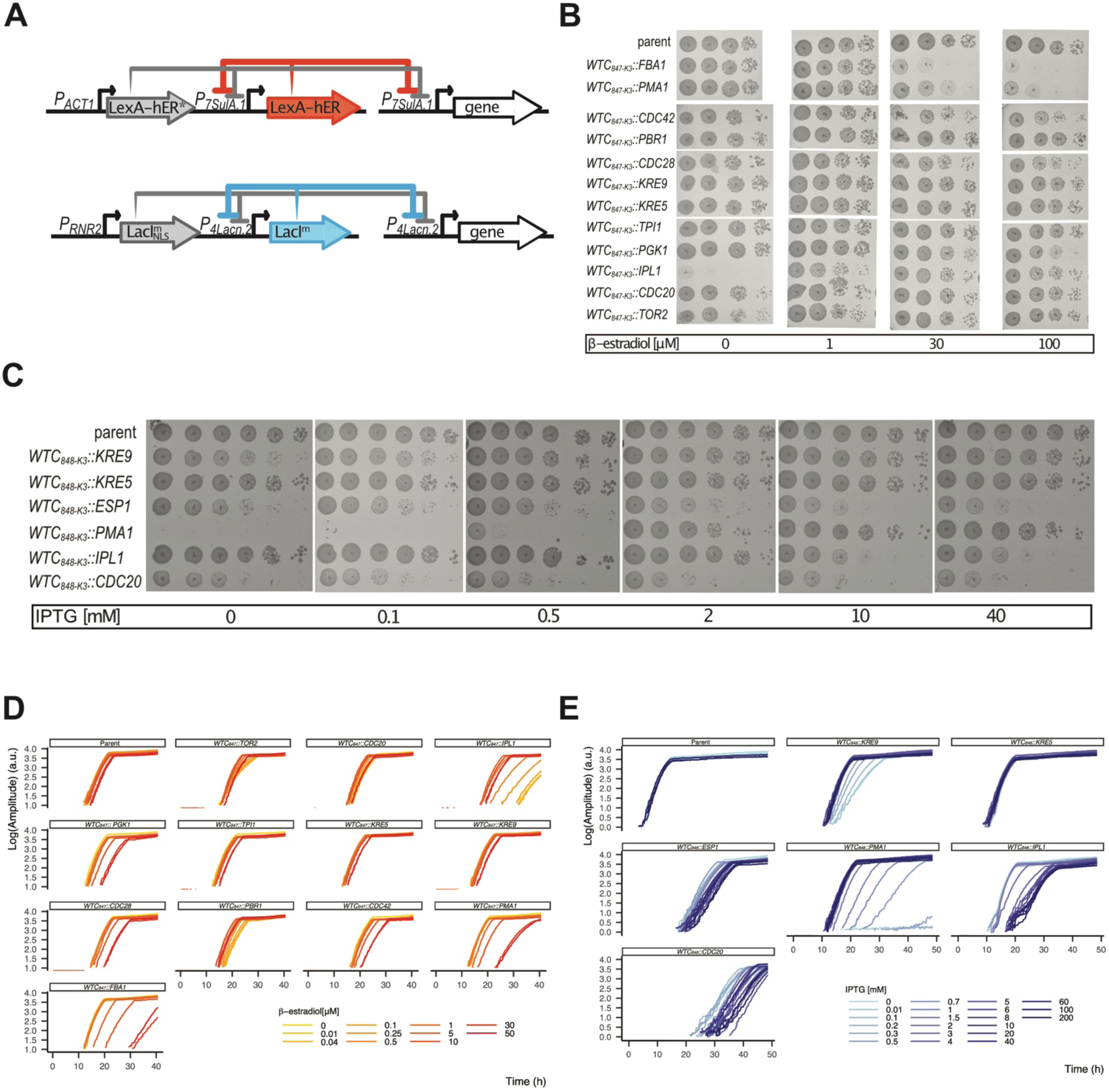
WTC847 and WTC848 alleles of essential endogenous genes show quantitative expression phenotypes. **A)** Schematic representation of the two WTC strains where endogenous genes were brought under WTC control. Top diagram depicts a strain where a gene of interest is under WTC847 control and the bottom diagram depicts a strain where a GOI is under WTC848 control. Grey indicates the SR portion of the circuit and the colours indicate the AR portion. * signifies the L387M mutation. **B)** Strains were created where all WTC847 control components were integrated and the gene indicated on the left was placed under the control of P7SulA.1 (Y3833-3837,3839-3841,3843-3845,3847). These were grown in presence of 30 µM β-estradiol for 8 hours and spotted onto YPD plates with various β-estradiol concentrations. The most concentrated spot on the left had 100.000 cells per spot, and each column is a 1:10 dilution. Parent refers to the strain where all WTC847 components were integrated but no endogenous gene was placed under its control (Y3809). K3 refers to the Kozak sequence AAGGGAAAAGGGAAA. **C)** Strains were created where all WTC848 control components were integrated and the gene indicated on the left was placed under the control of P4Lacn.2 (Y). These were grown in absence of IPTG for 8 hours and spotted onto YPD plates with various IPTG concentrations. The most concentrated spot on the left had 250.000 cells per spot, and each column is a 1:10 dilution. Parent refers to the strain where all WTC848 components were integrated but no endogenous gene was placed under its control (Y3870). **D)** Strains seen in (B) were grown overnight with 1µm β-estradiol, washed twice with YPD and grown without inducer for 6 hours. They were then diluted at a concentration of 100.000 cells per 250µL with various inducer concentrations into a Growth Profiler plate and growth curves were recorded in liquid culture. **E)** Same as in (D) for the strains seen in (C). Cells were grown overnight in 0.1mM IPTG (except for *WTC848::PMA1* strain, grown in 40mM IPTG) and washed and diluted as in (D).

### Appendix 5: Protocol for building strains where endogenous genes are regulated by WTC components

Well-tempered Controller (WTC) strains can be built in two steps: integration of all the components necessary for the operation of the system followed by replacement of the promoter of an endogenous gene with the relevant WTC promoter. We previously published a very detailed protocol on how to do this for WTC ^52^. This protocol can be followed with the modifications mentioned here to create single, double or triple WTC strains. We recommend that all necessary WTC plasmids in Step 1 are integrated (i.e. all three if triple gene control is desired) before moving onto Step 2 for endogenous promoter replacement.

**Step 1:** For generating a parent WTC strain, cut and linearize the following plasmids using AscI, followed by transformation into yeast, as explained in the published protocol:

WTC_846_: One plasmid containing all repressors as previously explained in the published protocol.

WTC_847_: One plasmid containing all repressors*. This plasmid is provided with one selection marker on Addgene P2802 (LYS2).

WTC_848_: Two successive transformations are required to integrate two separate plasmids. One plasmid contains all repressors**. This plasmid is provided with three different selection markers on Addgene: P2814(HIS3MX), P2880 (MET15), and P2881 (LEU2MX). A second plasmid is required, which carries the IPTG importer Lac12***. This plasmid is provided with five different selection markers on Addgene: P2072 (MET15), P2882 (LEU2MX), P2883 (LYS2), P2884 (URA3MX), P2885(HIS3MX).

* PACT1_LexA-L7-hER(L387M)_tCyc1_P7SulA.1_LexA-hER-adh1tail_tCyc1

**PRnr2_LacI(T167A,W220F,M254L,V119I,Q60G,M242L,NLS)_tFum1_P4Lacn.2_LacI(T167A,W220F,M254L,Q60G,M242L)_tCyc1

*** PACT1_Lac12_tCyc1

**Step 2:** For replacing the promoter of an endogenous gene, a PCR product is generated with two homology arms. Everything within the genome that lies in between the homology regions will be excised during homologous repair. 5’ homology arm corresponds to a region upstream of the ATG and the exact sequence determines whether the endogenous promoter is completely deleted or simply displaced by the WTC promoter. The 3’ homology arm corresponds to the ATG followed by ∼30bp of the gene of interest itself. This PCR product can then be used in either Basic homologous repair or the Advanced CRISPR-Cas9 protocol described previously for homologous repair using an antibiotic selection scheme. For more details on how to run this PCR, and more information on an alternative markerless integration, consult the published protocol. To generate the PCR fragment, use the following plasmids provided on Addgene:

WTC_846_: P2375 (Nourseothricin resistance) P2350 (Hygromycin resistance)

WTC_847_: P2754 (Nourseothricin resistance) P2755 (Hygromycin resistance)

WTC_848_: P2765 (Nourseothricin resistance) P2879 (Hygromycin resistance)

Exceptionally for WTC_848_, the annealing portion of the reverse primer differs from the published protocol. Use <aattgttatccgctcacaattaattgttatccgctcacaattggtgttttaaaac> as the annealing portion of this primer (Basic Protocol 2, step 4b in the published protocol).

## MATERIALS AND METHODS

### Plasmids

Details on all plasmids and the sequences therein can be found in tables 1 and 2. Plasmids with auxotrophic markers were based on the pRG vector series backbones^53^. Plasmids used to generate linear PCR products for endogenous gene promoter replacements or integrative transformations of some fluorescent proteins with an antibiotic marker were based on the pFA6 backbone^54^. The oligos used for these PCRs can be found in Table 3. All plasmids were constructed using isothermal assembly. Inserts were either generated from existing plasmids through PCR or synthesized as custom DNA (GeneArt, UK).

**Table 1.**
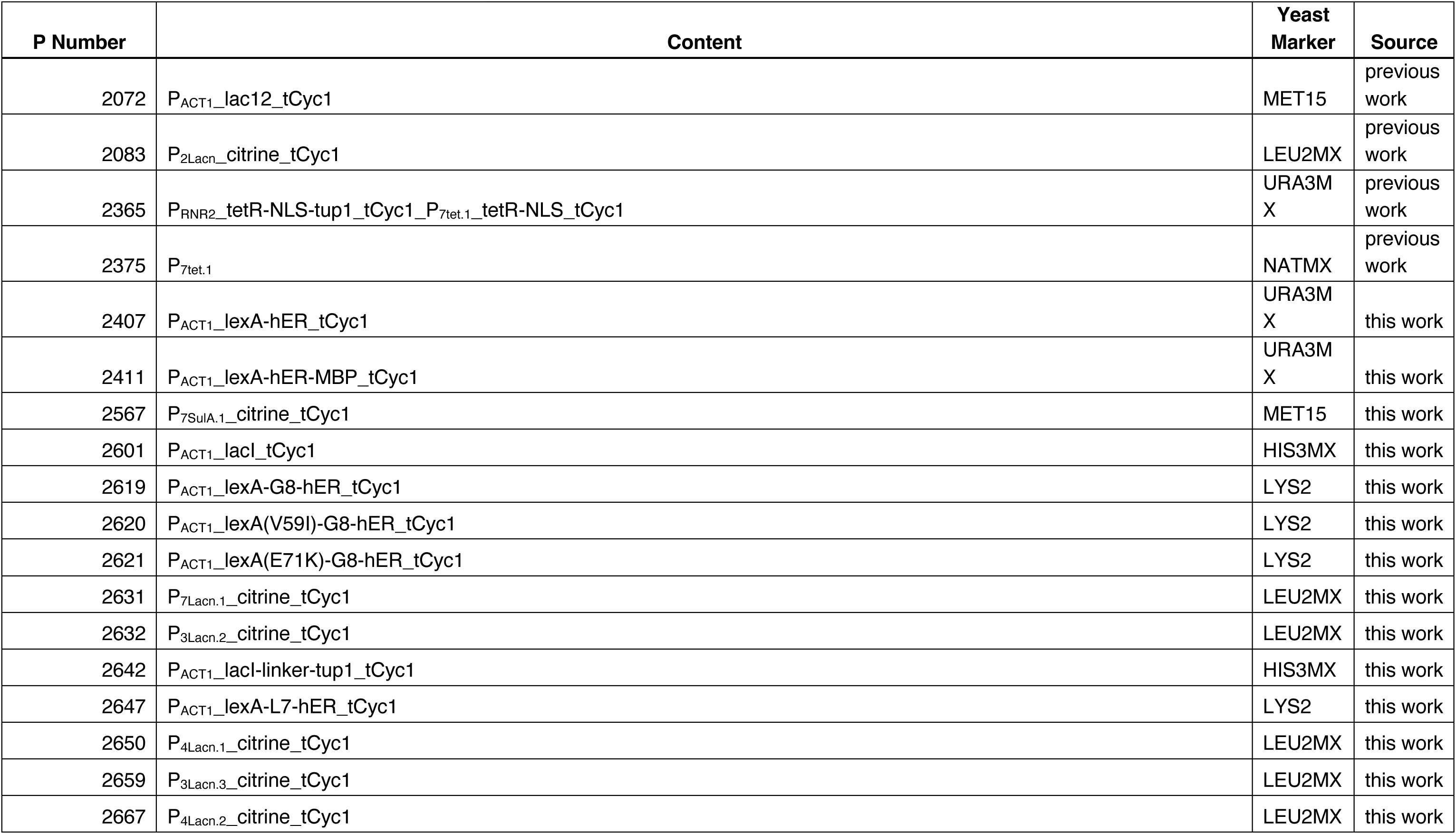
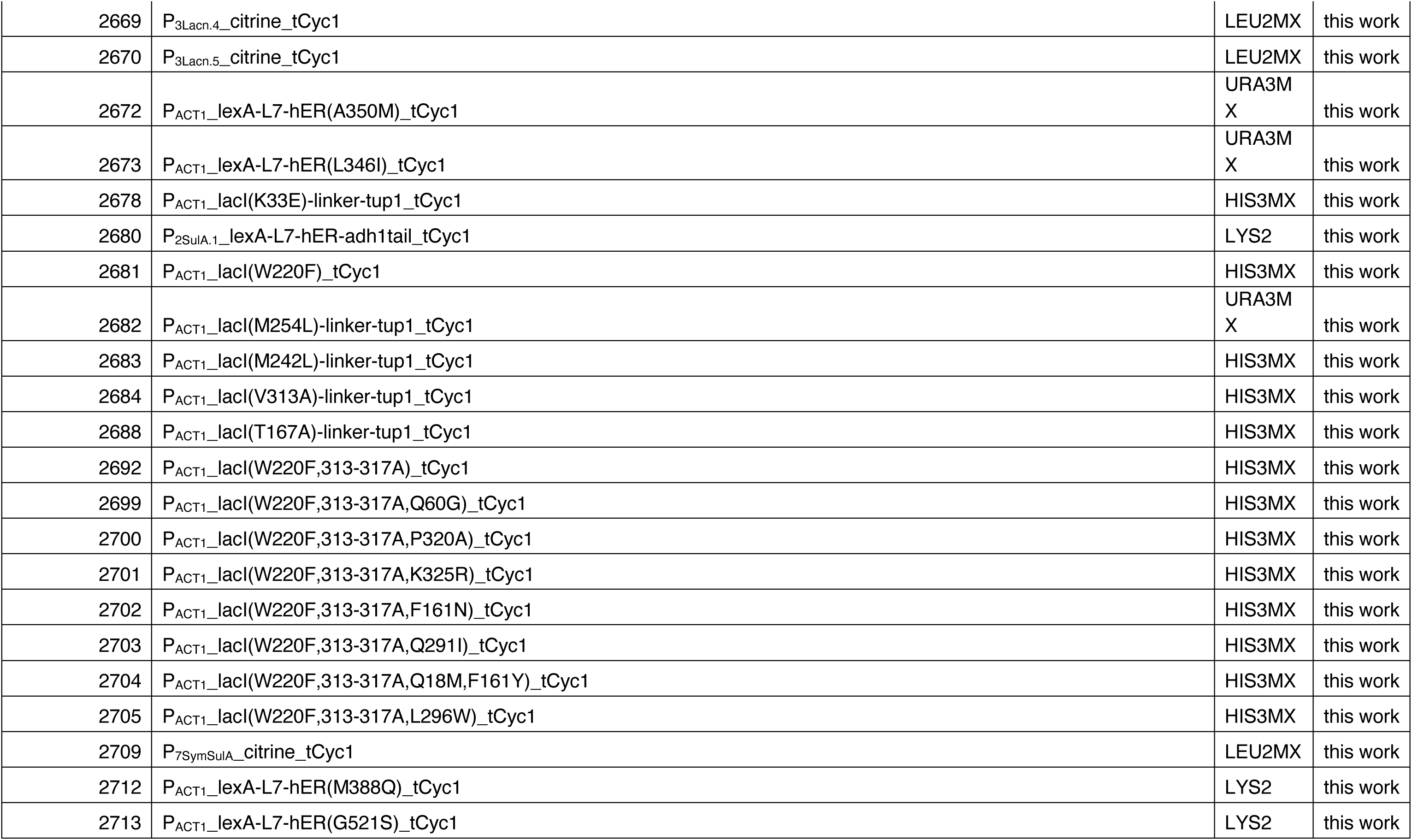
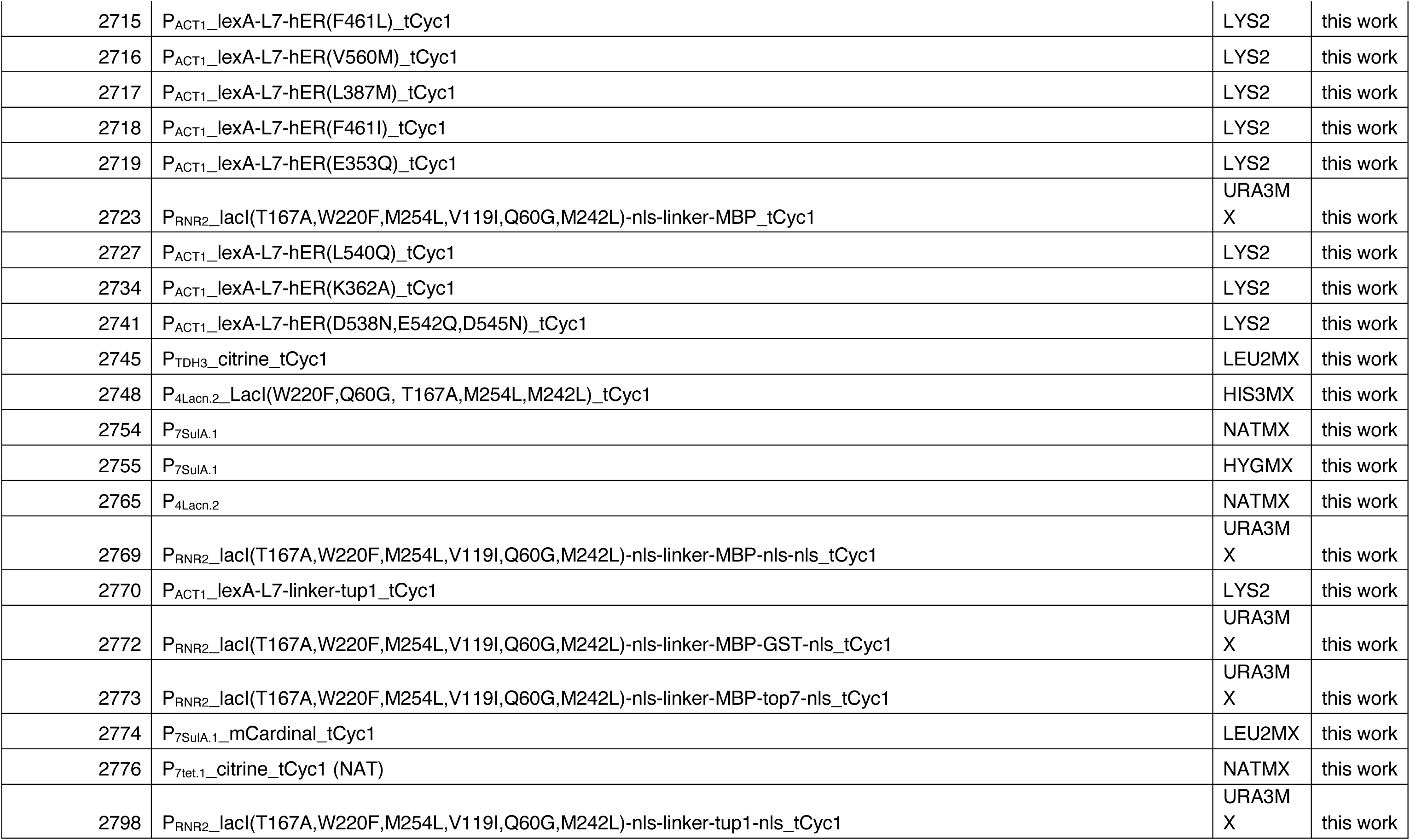
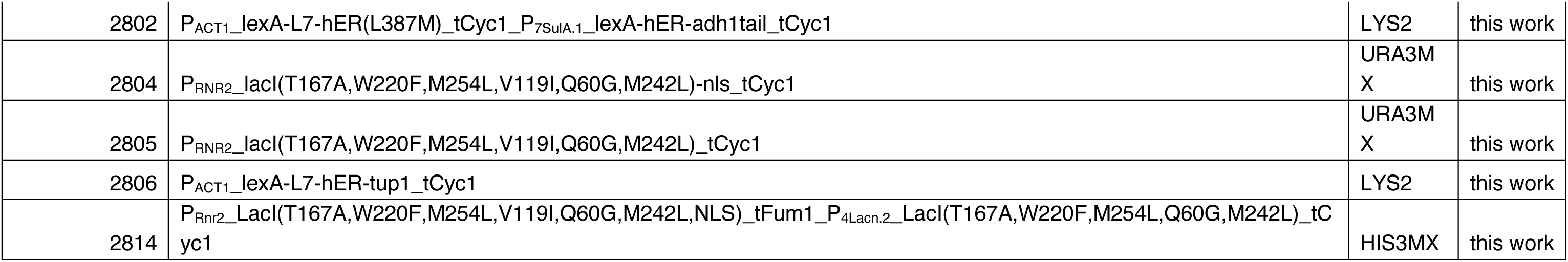
Plasmids used in this study.

**Table 2.**
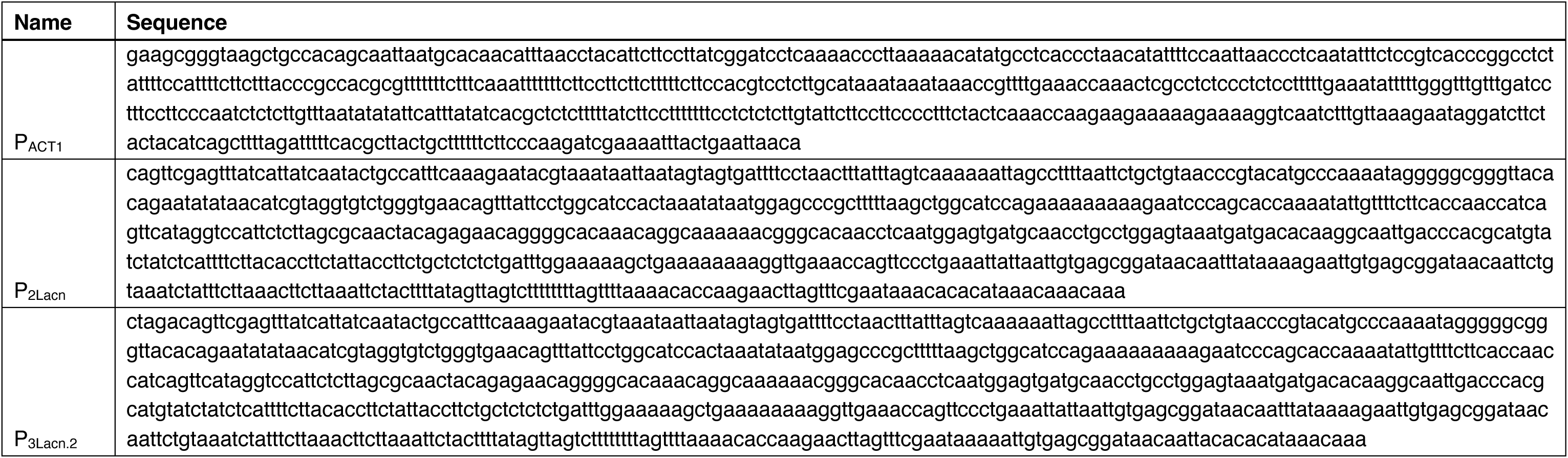
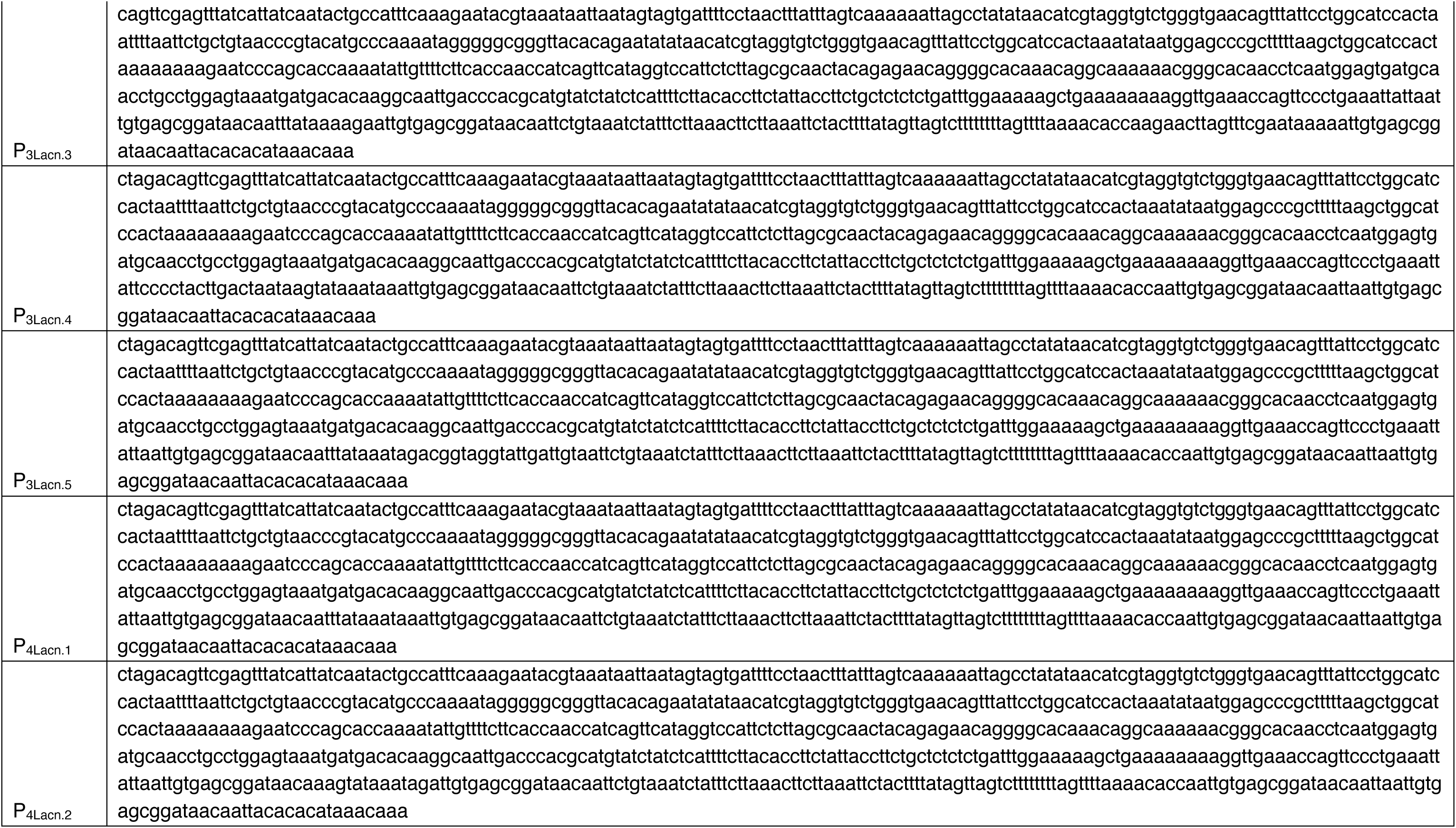
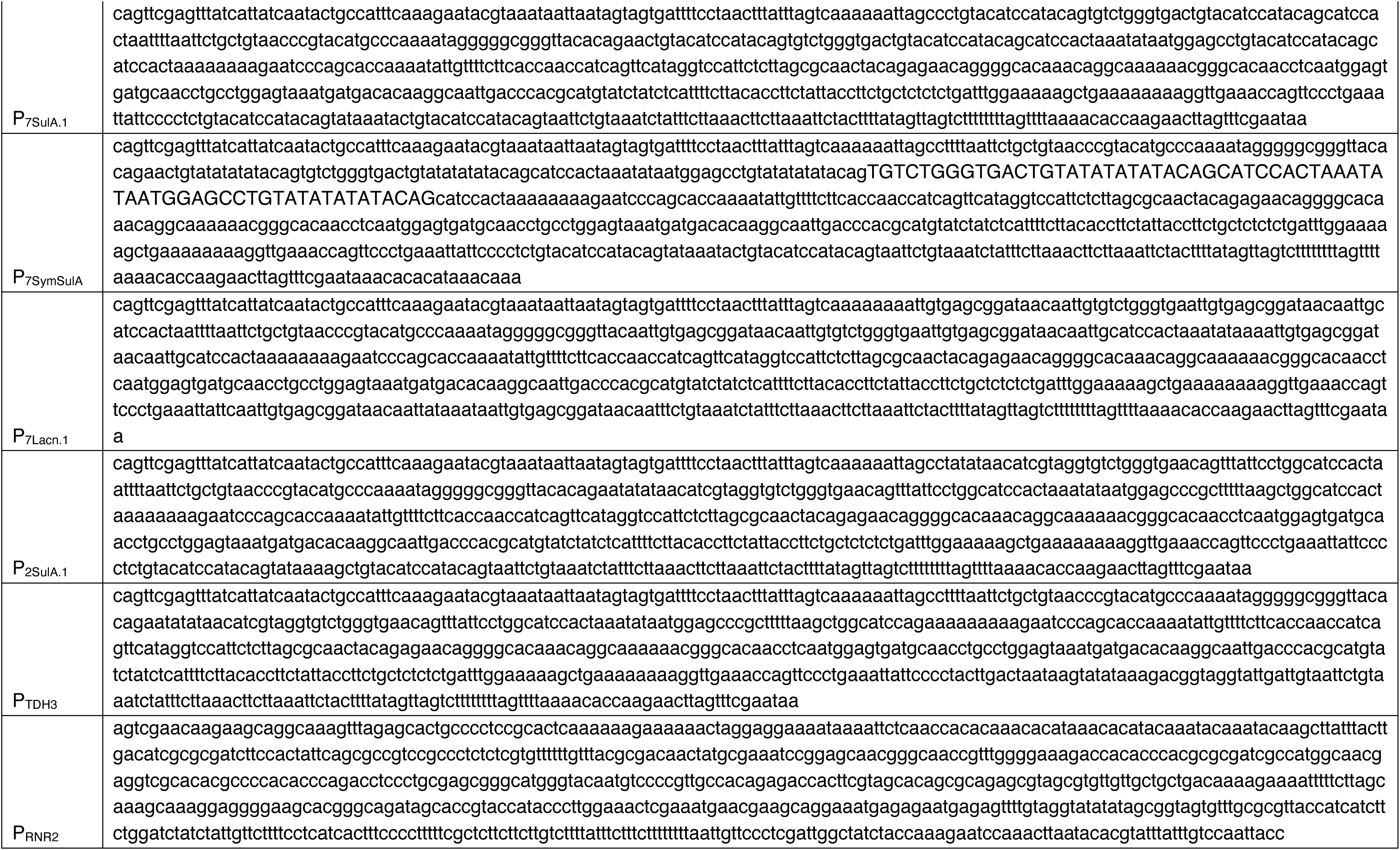
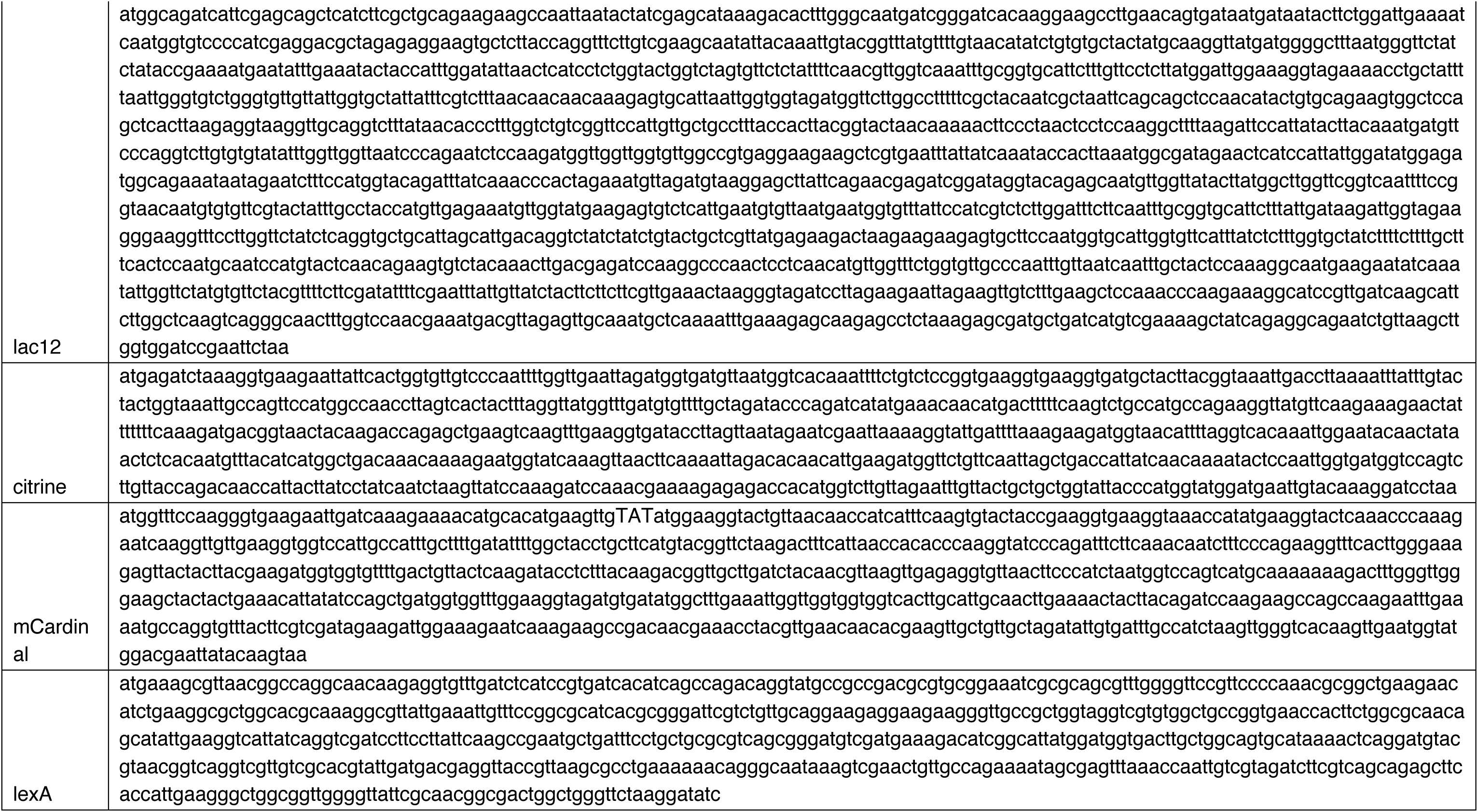
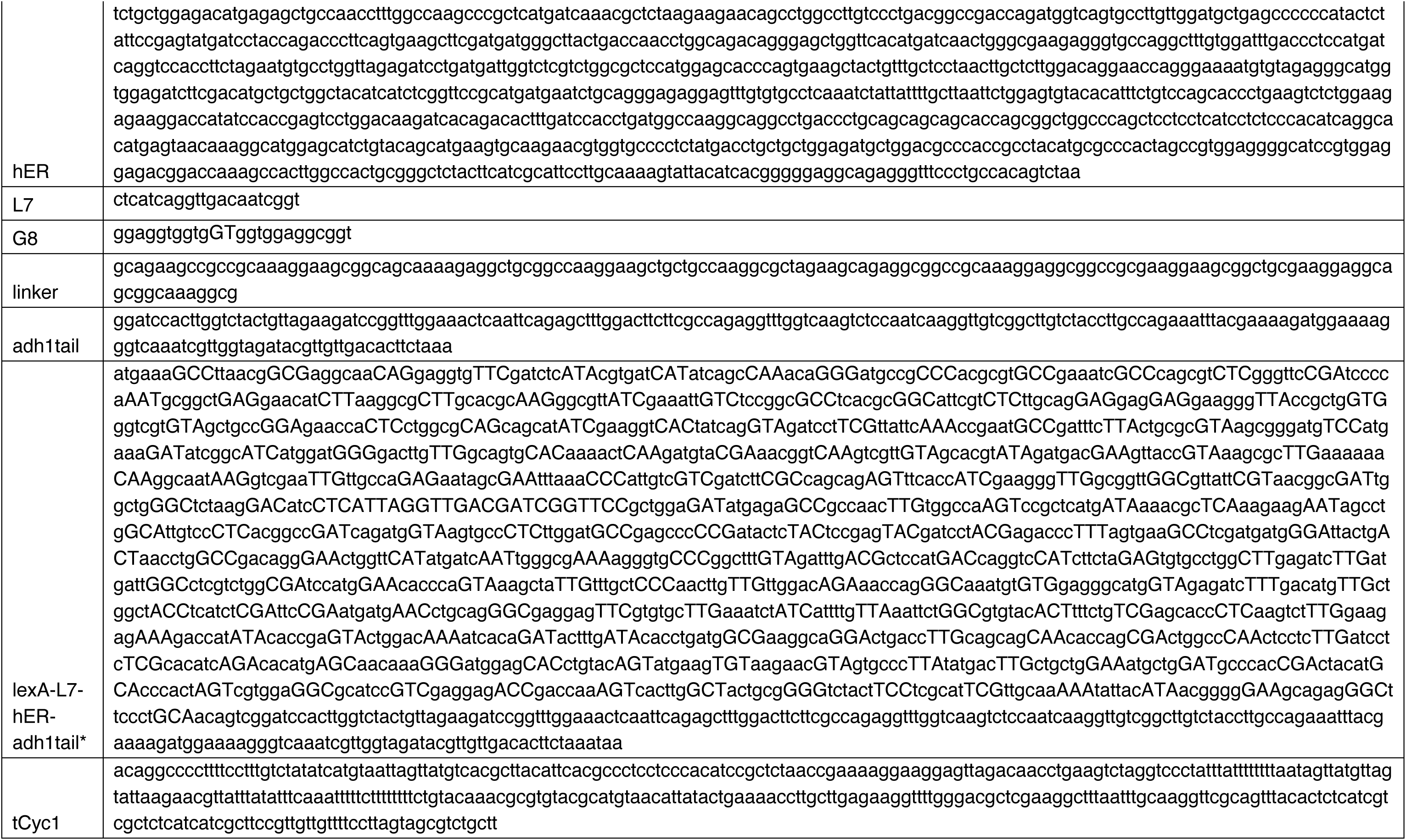
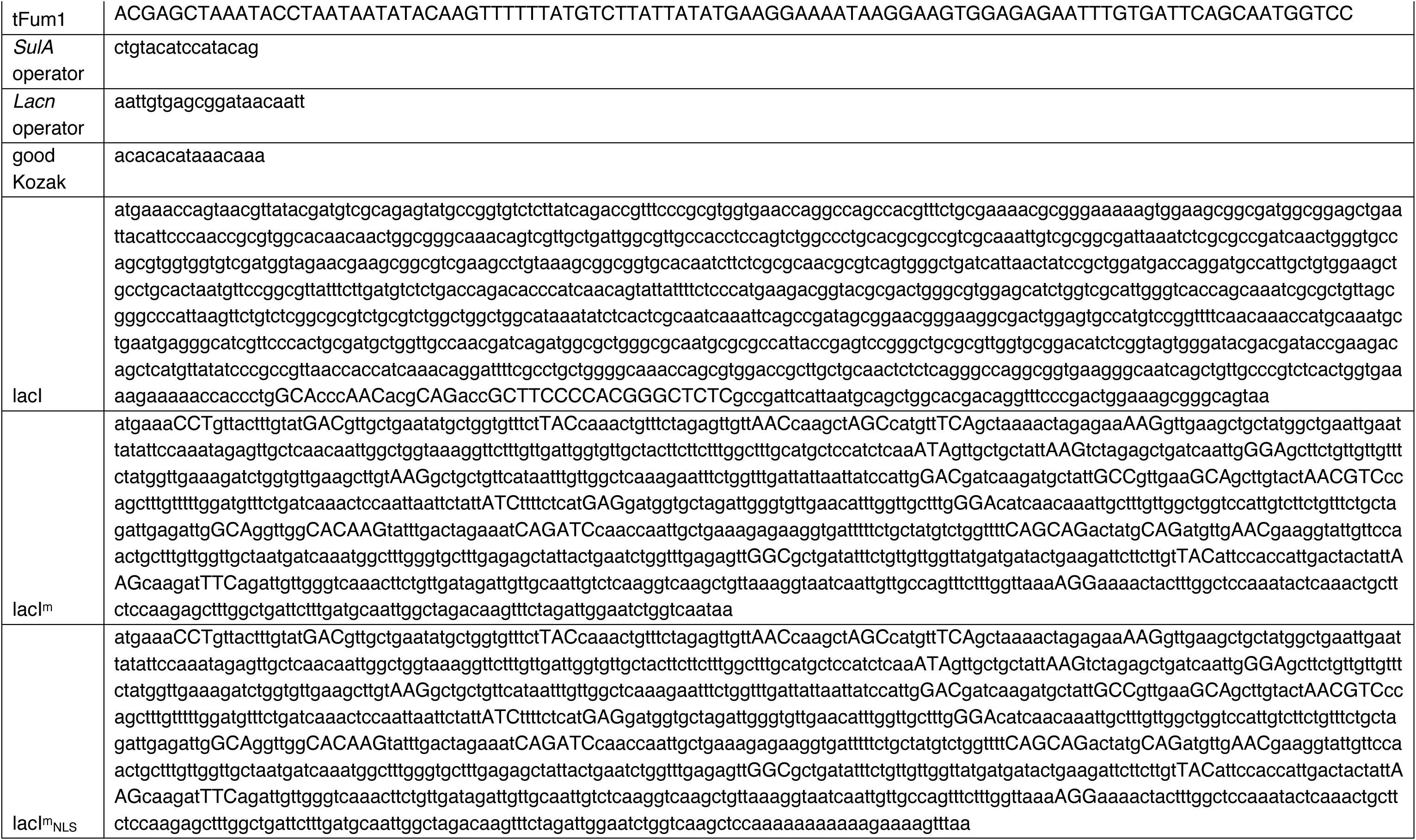
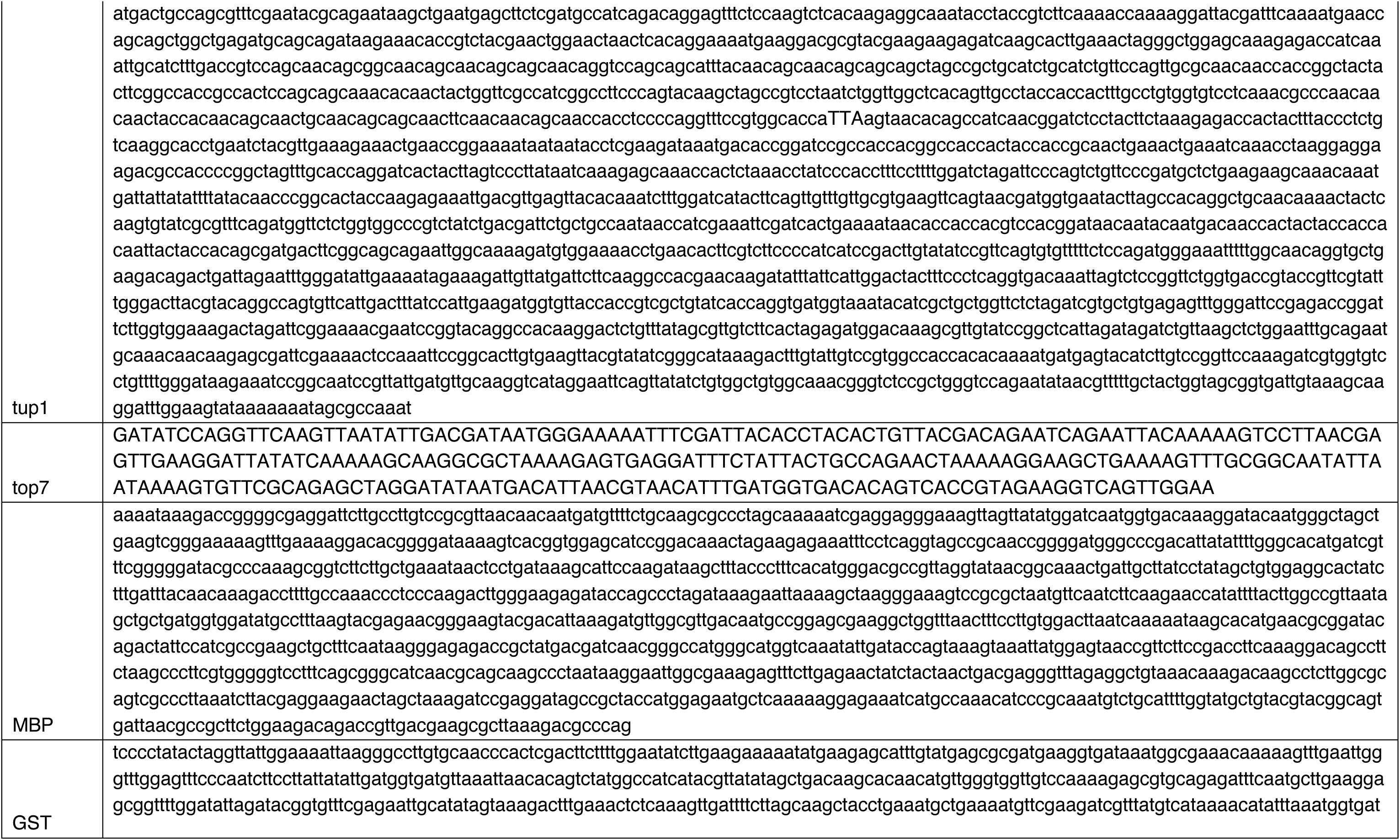
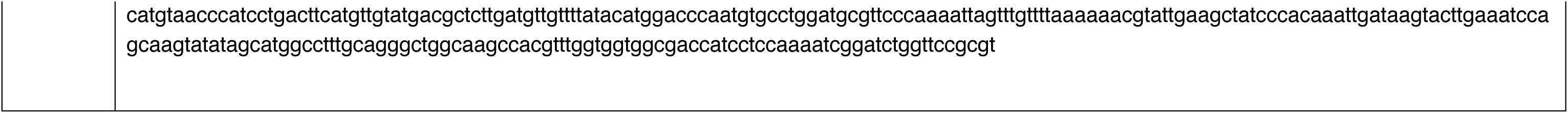
Relevant sequences used in this study. * indicates the lexA-hER-adh1tail construct with silent mutations introduced to avoid homologous recombination within the context of WTC847, where two lexA-hER-adh1tail genes are integrated with only a single amino acid difference between them.

**Table 3.**
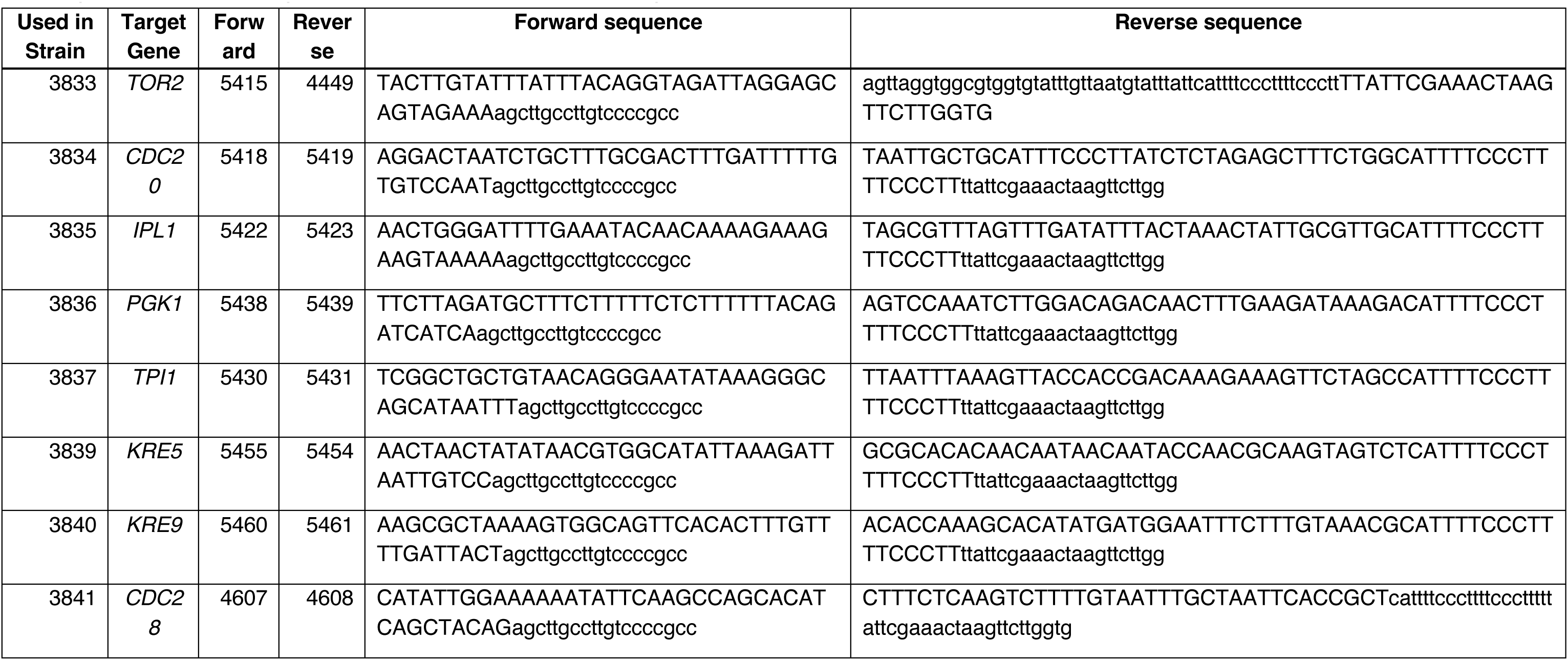
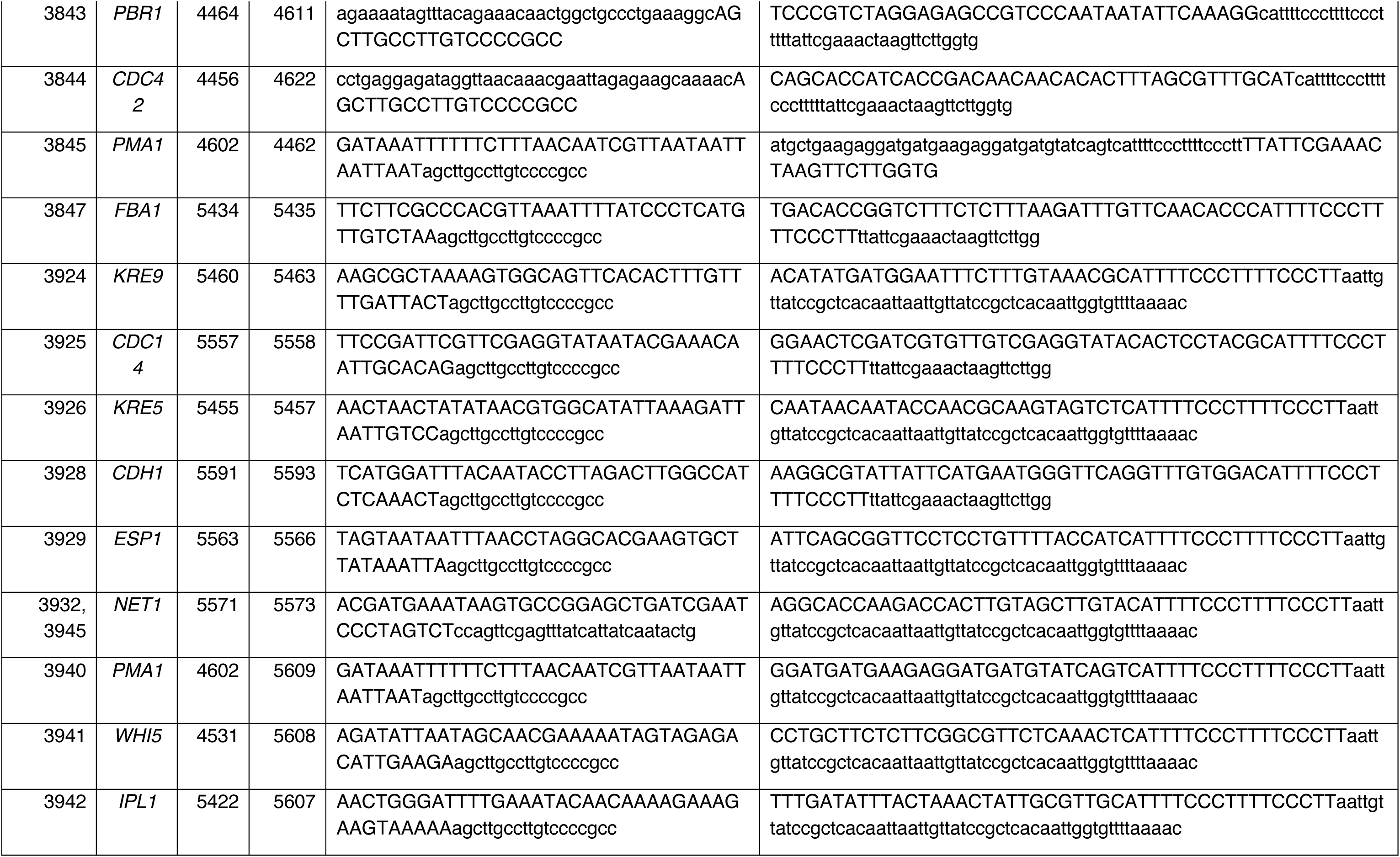
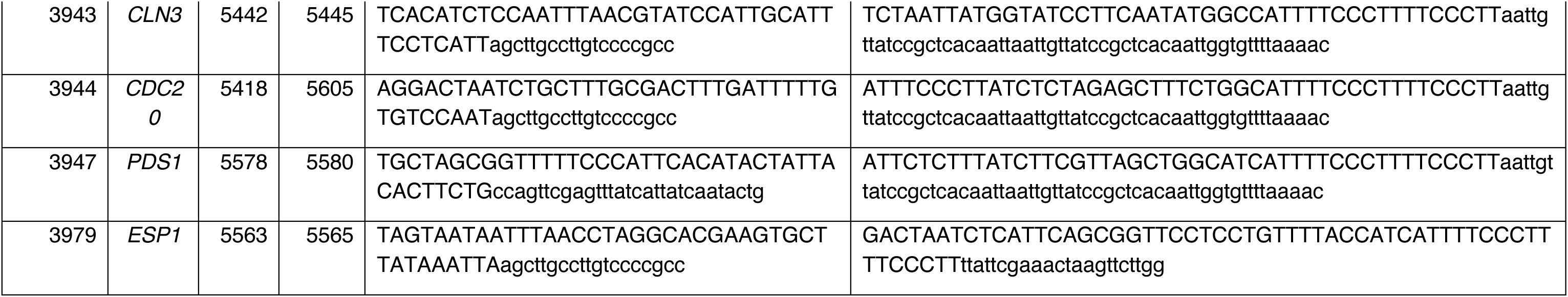
Number and sequences of the forward and reverse oligos used for generating linear PCR products containing either P7SulA.1 or P4Lacn.2, and with or without an antibiotic marker. These linear fragments were used in transformations where endogenous promoters were replaced with one of the two WTC promoters.

There are cases where two plasmids integrated in the same strain contained the same promoter and terminator. In these cases, the inserts were cloned into the plasmid backbone in opposite directions by placing one insert on the sense and the other on the antisense strand, as explained before^19^. This was done to prevent homologous recombination between the inserts during genomic integration.

### Strains

Details on all strains used in this study can be found in Table 4. All strains generated for this study were based on a haploid derivative of the BY4743 background. In strains where WTC promoters P7SulA.1 and P4Lacn.2 replaced endogenous promoters, we used a standard forward oligo annealing to the start of both promoters, and gene specific reverse oligos to confirm correct integration. The sequences of these oligos can be found in Table 5 and a detailed protocol on how to create such strains is given in Appendix 5.

**Table 4.** Strains used in this study. Kozak sequence is the last 15bp before the start codon.

| Y number | Name | Genotype | Kozak Sequence | Yeast Parent | Integrated Plasmid | Source |
| --- | --- | --- | --- | --- | --- | --- |
| 70 | autofluorescence | BY4743 derivative, haploid, MATa *his3Δ1 leu2Δ0 met15Δ0 ura3Δ0 lys2Δ0* |  | BY4743 | - | previous work |
| 2761 | WTC<sub>846</sub>:: | *ura3Δ::P<sub>RNR2</sub>_tetR-NLS-tup1_tCyc1_P<sub>7tet.1</sub>_tetR-NLS_tCyc1-URA3* |  | Y70 | P2365 | this work |
| 3316 | Lac12 | *met15Δ::P<sub>ACT1</sub>_lac12_tCyc1-MET15* |  | Y70 | P2072 | this work |
| 3391 | P<sub>7SulA.1</sub>-const | *met15Δ::P<sub>7SulA.1</sub>_citrine_tCyc1-MET15* |  | Y70 | P2567 | this work |
| 3402 | LexA-hER-MBP | *met15Δ::P<sub>7SulA.1</sub>_citrine_tCyc1-MET15, ura3Δ::P<sub>ACT1</sub>_lexA-hER-MBP_tCyc1-URA3* |  | Y3391 | P2411 | this work |
| 3408 | P<sub>7SulA.1</sub>-SR | *met15Δ::P<sub>7SulA.1</sub>_citrine_tCyc1-MET15, ura3Δ::P<sub>ACT1</sub>_lexA-hER_tCyc1-URA3* |  | Y3391 | P2407 | this work |
| 3412 | P<sub>2Lacn</sub>-const | *met15Δ::P<sub>ACT1</sub>_lac12_tCyc1-MET15, leu2Δ::P<sub>2Lacn</sub>_citrine_tCyc1-LEU2* |  | Y3316 | P2803 | this work |
| 3420 | P<sub>2Lacn</sub>-SR | *met15Δ::P<sub>ACT1</sub>_lac12_tCyc1-MET15, leu2Δ::P<sub>2Lacn</sub>_citrine_tCyc1-LEU2, his3Δ::P<sub>ACT1</sub>_lacI_tCyc1-HIS3* |  | Y3412 | P2601 | this work |
| 3438 | LexA-G8-hER | *met15Δ::P<sub>7SulA.1</sub>_citrine_tCyc1-MET15, lys2Δ::P<sub>ACT1</sub>_lexA-G8-hER_tCyc1-LYS2* |  | Y3391 | P2619 | this work |
| 3441 | LexA(V59I)-G8-hER | *met15Δ::P<sub>7SulA.1</sub>_citrine_tCyc1-MET15, lys2Δ::P<sub>ACT1</sub>_lexA(V59I)-G8-hER_tCyc1-LYS2* |  | Y3391 | P2620 | this work |
| 3442 | LexA(E71K)-G8-hER | *met15Δ::P<sub>7SulA.1</sub>_citrine_tCyc1-MET15, lys2Δ::P<sub>ACT1</sub>_lexA(E71K)-G8-hER_tCyc1-LYS2* |  | Y3391 | P2621 | this work |
| 3463 | P<sub>3Lacn.2</sub>-const | *met15Δ::P<sub>ACT1</sub>_lac12_tCyc1-MET15, leu2Δ::P<sub>3Lacn.2</sub>_citrine_tCyc1-LEU2* |  | Y3316 | P2632 | this work |
| 3464 | P<sub>7Lacn.1</sub>-const | *met15Δ::P<sub>ACT1</sub>_lac12_tCyc1-MET15, leu2Δ::P<sub>7Lacn.1</sub>_citrine_tCyc1-LEU2* |  | Y3316 | P2631 | this work |
| 3470 | P<sub>3Lacn.2</sub>-SR | *met15Δ::P<sub>ACT1</sub>_lac12_tCyc1-MET15, leu2Δ::P<sub>3Lacn.2</sub>_citrine_tCyc1-LEU2, his3Δ::P<sub>ACT1</sub>_lacI_tCyc1-HIS3* |  | Y3463 | P2601 | this work |
| 3497 | P<sub>4Lacn.1</sub>-const | *met15Δ::P<sub>ACT1</sub>_lac12_tCyc1-MET15, leu2Δ::P<sub>4Lacn.1</sub>_citrine_tCyc1-LEU2* |  | Y3316 | P2650 | this work |
| 3498 | P<sub>4Lacn.1</sub>-SR | *met15Δ::P<sub>ACT1</sub>_lac12_tCyc1-MET15, leu2Δ::P<sub>4Lacn.1</sub>_citrine_tCyc1-LEU2, his3Δ::P<sub>ACT1</sub>_lacI_tCyc1-HIS3* |  | Y3497 | P2601 | this work |
| 3516 | P<sub>7SulA.1</sub>-SR-L7 | *met15Δ::P<sub>7SulA.1</sub>_citrine_tCyc1-MET15, lys2Δ::P<sub>ACT1</sub>_lexA-L7-hER_tCyc1-LYS2* |  | Y3391 | P2647 | this work |
| 3518 | P<sub>3Lacn.3</sub>-const | *met15Δ::P<sub>ACT1</sub>_lac12_tCyc1-MET15, leu2Δ::P<sub>3Lacn.3</sub>_citrine_tCyc1-LEU2* |  | Y3316 | P2659 | this work |
| 3539 | P<sub>3Lacn.5</sub>-const | *met15Δ::P<sub>ACT1</sub>_lac12_tCyc1-MET15, leu2Δ::P<sub>3Lacn.5</sub>_citrine_tCyc1-LEU2* |  | Y3316 | P2670 | this work |
| 3540 | P<sub>3Lacn.4</sub>-const | *met15Δ::P<sub>ACT1</sub>_lac12_tCyc1-MET15, leu2Δ::P<sub>3Lacn.4</sub>_citrine_tCyc1-LEU2* |  | Y3316 | P2669 | this work |
| 3542 | P<sub>4Lacn.2</sub>-const | *met15Δ::P<sub>ACT1</sub>_lac12_tCyc1-MET15, leu2Δ::P<sub>4Lacn.2</sub>_citrine_tCyc1-LEU2* |  | Y3316 | P2667 | this work |
| 3544 | P<sub>3Lacn.5</sub>-SR | *met15Δ::P<sub>ACT1</sub>_lac12_tCyc1-MET15, leu2Δ::P<sub>3Lacn.5</sub>_citrine_tCyc1-LEU2, his3Δ::P<sub>ACT1</sub>_lacI_tCyc1-HIS3* |  | Y3539 | P2601 | this work |
| 3545 | P<sub>3Lacn.4</sub>-SR | *met15Δ::P<sub>ACT1</sub>_lac12_tCyc1-MET15, leu2Δ::P<sub>3Lacn.4</sub>_citrine_tCyc1-LEU2, his3Δ::P<sub>ACT1</sub>_lacI_tCyc1-HIS3* |  | Y3540 | P2601 | this work |
| 3547 | P<sub>4Lacn.2</sub>-SR | *met15Δ::P<sub>ACT1</sub>_lac12_tCyc1-MET15, leu2Δ::P<sub>4Lacn.2</sub>_citrine_tCyc1-LEU2, his3Δ::P<sub>ACT1</sub>_lacI_tCyc1-HIS3* |  | Y3542 | P2601 | this work |
| 3554 | LacI(K33E)-Tup1 | *met15Δ::P<sub>ACT1</sub>_lac12_tCyc1-MET15, leu2Δ::P<sub>4Lacn.2</sub>_citrine_tCyc1-LEU2, his3Δ::P<sub>ACT1</sub>_lacI(K33E)-linker-tup1_tCyc1-HIS3* |  | Y3542 | P2678 | this work |
| 3560 | hER(L346I) | *met15Δ::P<sub>7SulA.1</sub>_citrine_tCyc1-MET15, ura3Δ::P<sub>ACT1</sub>_lexA-L7-hER(L346I)_tCyc1-URA3* |  | Y3391 | P2673 | this work |
| 3561 | hER(A350M) | *met15Δ::P<sub>7SulA.1</sub>_citrine_tCyc1-MET15, ura3Δ::P<sub>ACT1</sub>_lexA-L7-hER(A350M)_tCyc1-URA3* |  | Y3391 | P2672 | this work |
| 3570 | LexA-L7-hER | *lys2Δ::P<sub>ACT1</sub>_lexA-L7-hER_tCyc1-LYS2* |  | Y70 | P2647 | this work |
| 3572 | P<sub>7SulA.1</sub>-AR | *met15Δ::P<sub>7SulA.1</sub>_citrine_tCyc1-MET15, P<sub>2SulA.1</sub>_lexA-L7-hER-adh1tail_tCyc1-LYS2* |  | Y3391 | P2680 | this work |
| 3574 | LacI(W220F) | *met15Δ::P<sub>ACT1</sub>_lac12_tCyc1-MET15, leu2Δ::P<sub>4Lacn.2</sub>_citrine_tCyc1-LEU2, his3Δ::P<sub>ACT1</sub>_lacI(W220F)_tCyc1-HIS3* |  | Y3542 | P2681 | this work |
| 3581 | LacI-Tup1 | *met15Δ::P<sub>ACT1</sub>_lac12_tCyc1-MET15, leu2Δ::P<sub>4Lacn.2</sub>_citrine_tCyc1-LEU2, his3Δ::P<sub>ACT1</sub>_lacI-linker-tup1_tCyc1-HIS3* |  | Y3542 | P2642 | this work |
| 3582 | LacI(T167A)-Tup1 | *met15Δ::P<sub>ACT1</sub>_lac12_tCyc1-MET15, leu2Δ::P<sub>4Lacn.2</sub>_citrine_tCyc1-LEU2, his3Δ::P<sub>ACT1</sub>_lacI(T167A)-linker-tup1_tCyc1-HIS3* |  | Y3542 | P2688 | this work |
| 3583 | LacI(M242L)-Tup1 | *met15Δ::P<sub>ACT1</sub>_lac12_tCyc1-MET15, leu2Δ::P<sub>4Lacn.2</sub>_citrine_tCyc1-LEU2, his3Δ::P<sub>ACT1</sub>_lacI(M242L)-linker-tup1_tCyc1-HIS3* |  | Y3542 | P2683 | this work |
| 3584 | LacI(V313A)-Tup1 | *met15Δ::P<sub>ACT1</sub>_lac12_tCyc1-MET15, leu2Δ::P<sub>4Lacn.2</sub>_citrine_tCyc1-LEU2, his3Δ::P<sub>ACT1</sub>_lacI(V313A)-linker-tup1_tCyc1-HIS3* |  | Y3542 | P2684 | this work |
| 3591 | LacI(M254L)-Tup1 | *met15Δ::P<sub>ACT1</sub>_lac12_tCyc1-MET15, leu2Δ::P<sub>4Lacn.2</sub>_citrine_tCyc1-LEU2, ura3Δ::P<sub>ACT1</sub>_lacI(M254L)-linker-tup1_tCyc1-URA3* |  | Y3542 | P2682 | this work |
| 3593 | LacI(W220F,313-317A) | *met15Δ::P<sub>ACT1</sub>_lac12_tCyc1-MET15, leu2Δ::P<sub>4Lacn.2</sub>_citrine_tCyc1-LEU2, his3Δ::P<sub>ACT1</sub>_lacI(W220F,313-317A)_tCyc1-HIS3* |  | Y3542 | P2692 | this work |
| 3603 | LacI(W220F,313-317A,Q60G) | *met15Δ::P<sub>ACT1</sub>_lac12_tCyc1-MET15, leu2Δ::P<sub>4Lacn.2</sub>_citrine_tCyc1-LEU2, his3Δ::P<sub>ACT1</sub>_lacI(W220F,313-317A,Q60G)_tCyc1-HIS3* |  | Y3542 | P2699 | this work |
| 3604 | LacI(W220F,313-317A,P320A) | *met15Δ::P<sub>ACT1</sub>_lac12_tCyc1-MET15, leu2Δ::P<sub>4Lacn.2</sub>_citrine_tCyc1-LEU2, his3Δ::P<sub>ACT1</sub>_lacI(W220F,313-317A,P320A)_tCyc1-HIS3* |  | Y3542 | P2700 | this work |
| 3605 | LacI(W220F,313-317A,K325R) | *met15Δ::P<sub>ACT1</sub>_lac12_tCyc1-MET15, leu2Δ::P<sub>4Lacn.2</sub>_citrine_tCyc1-LEU2, his3Δ::P<sub>ACT1</sub>_lacI(W220F,313-317A,K325R)_tCyc1-HIS3* |  | Y3542 | P2701 | this work |
| 3606 | LacI(W220F,313-317A,F161N) | *met15Δ::P<sub>ACT1</sub>_lac12_tCyc1-MET15, leu2Δ::P<sub>4Lacn.2</sub>_citrine_tCyc1-LEU2, his3Δ::P<sub>ACT1</sub>_lacI(W220F,313-317A,F161N)_tCyc1-HIS3* |  | Y3542 | P2702 | this work |
| 3607 | LacI(W220F,313-317A,Q291I) | *met15Δ::P<sub>ACT1</sub>_lac12_tCyc1-MET15, leu2Δ::P<sub>4Lacn.2</sub>_citrine_tCyc1-LEU2, his3Δ::P<sub>ACT1</sub>_lacI(W220F,313-317A,Q291I)_tCyc1-HIS3* |  | Y3542 | P2703 | this work |
| 3612 | LacI(W220F,313-317A,Q18M,F161Y) | *met15Δ::P<sub>ACT1</sub>_lac12_tCyc1-MET15, leu2Δ::P<sub>4Lacn.2</sub>_citrine_tCyc1-LEU2, his3Δ::P<sub>ACT1</sub>_lacI(W220F,313-317A,Q18M,F161Y)_tCyc1-HIS3* |  | Y3542 | P2704 | this work |
| 3613 | LacI(W220F,313-317A,L296W) | *met15Δ::P<sub>ACT1</sub>_lac12_tCyc1-MET15, leu2Δ::P<sub>4Lacn.2</sub>_citrine_tCyc1-LEU2, his3Δ::P<sub>ACT1</sub>_lacI(W220F,313-317A,L296W)_tCyc1-HIS3* |  | Y3542 | P2705 | this work |
| 3620 | P<sub>7SymSulA</sub>-SR | *lys2Δ::P<sub>ACT1</sub>_lexA-L7-hER_tCyc1-LYS2, leu2Δ::P<sub>7SymSulA</sub>_citrine_tCyc1-LEU2* |  | Y3570 | P2709 | this work |
| 3622 | P<sub>7SymSulA</sub>-const | *leu2Δ::P<sub>7SymSulA</sub>_citrine_tCyc1-LEU2* |  | Y70 | P2709 | this work |
| 3626 | hER(M388Q) | *met15Δ::P<sub>7SulA.1</sub>_citrine_tCyc1-MET15, lys2Δ::P<sub>ACT1</sub>_lexA-L7-hER(M388Q)_tCyc1-LYS2* |  | Y3391 | P2712 | this work |
| 3627 | hER(G521S) | *met15Δ::P<sub>7SulA.1</sub>_citrine_tCyc1-MET15, lys2Δ::P<sub>ACT1</sub>_lexA-L7-hER(G521S)_tCyc1-LYS2* |  | Y3391 | P2713 | this work |
| 3629 | hER(F461L) | *met15Δ::P<sub>7SulA.1</sub>_citrine_tCyc1-MET15, lys2Δ::P<sub>ACT1</sub>_lexA-L7-hER(F461L)_tCyc1-LYS2* |  | Y3391 | P2715 | this work |
| 3630 | hER(V560M) | *met15Δ::P<sub>7SulA.1</sub>_citrine_tCyc1-MET15, lys2Δ::P<sub>ACT1</sub>_lexA-L7-hER(V560M)_tCyc1-LYS2* |  | Y3391 | P2716 | this work |
| 3631 | hER(L387M) | *met15Δ::P<sub>7SulA.1</sub>_citrine_tCyc1-MET15, lys2Δ::P<sub>ACT1</sub>_lexA-L7-hER(L387M)_tCyc1-LYS2* |  | Y3391 | P2717 | this work |
| 3632 | hER(F461I) | *met15Δ::P<sub>7SulA.1</sub>_citrine_tCyc1-MET15, lys2Δ::P<sub>ACT1</sub>_lexA-L7-hER(F461I)_tCyc1-LYS2* |  | Y3391 | P2718 | this work |
| 3633 | hER(E353Q) | *met15Δ::P<sub>7SulA.1</sub>_citrine_tCyc1-MET15, lys2Δ::P<sub>ACT1</sub>_lexA-L7-hER(E353Q)_tCyc1-LYS2* |  | Y3391 | P2719 | this work |
| 3641 | LacIm-MBP | *met15Δ::P<sub>ACT1</sub>_lac12_tCyc1-MET15, leu2Δ::P<sub>4Lacn.2</sub>_citrine_tCyc1-LEU2, ura3Δ::P<sub>RNR2</sub>_lacI(T167A,W220F,M254L,V119I,Q60G,M242L)-nls-linker-MBP_tCyc1-URA3* |  | Y3542 | P2723 | this work |
| 3642 | LexA-L7-hER(L540Q) | *met15Δ::P<sub>7SulA.1</sub>_citrine_tCyc1-MET15, lys2Δ::P<sub>ACT1</sub>_lexA-L7-hER(L540Q)_tCyc1-LYS2* |  | Y3391 | P2727 | this work |
| 3651 | LexA-L7-hER(K362A) | *met15Δ::P<sub>7SulA.1</sub>_citrine_tCyc1-MET15, lys2Δ::P<sub>ACT1</sub>_lexA-L7-hER(K362A)_tCyc1-LYS2* |  | Y3391 | P2734 | this work |
| 3664 | LexA-L7-hER(D538N,E542Q,D545N) | *met15Δ::P<sub>7SulA.1</sub>_citrine_tCyc1-MET15, lys2Δ::P<sub>ACT1</sub>_lexA-L7-hER(D538N,E542Q,D545N)_tCyc1-LYS2* |  | Y3391 | P2741 | this work |
| 3668 | P<sub>TDH3</sub>-const | *leu2Δ::P<sub>TDH3</sub>_citrine_tCyc1-LEU2* |  | Y70 | P2745 | this work |
| 3674 | P<sub>4Lacn.2</sub>-AR | *met15Δ::P<sub>ACT1</sub>_lac12_tCyc1-MET15, leu2Δ::P<sub>4Lacn.2</sub>_citrine_tCyc1-LEU2, his3Δ::P<sub>4Lacn.2</sub>_LacI(W220F,Q60G, T167A,M254L,M242L)_tCyc1-HIS3* |  | Y3542 | P2748 | this work |
| 3718 | LacIm-MBP-nls-nls | *met15Δ::P<sub>ACT1</sub>_lac12_tCyc1-MET15, leu2Δ::P<sub>4Lacn.2</sub>_citrine_tCyc1-LEU2, ura3Δ::P<sub>RNR2</sub>_lacI(T167A,W220F,M254L, V119I,Q60G,M242L)-nls-linker-MBP-nls-nls_tCyc1-URA3* |  | Y3542 | P2769 | this work |
| 3719 | LexA-L7-Tup1 | *met15Δ::P<sub>7SulA.1</sub>_citrine_tCyc1-MET15, lys2Δ::P<sub>ACT1</sub>_lexA-L7-linker-tup1_tCyc1-LYS2* |  | Y3391 | P2770 | this work |
| 3722 | LacIm-MBP-GST-nls | *met15Δ::P<sub>ACT1</sub>_lac12_tCyc1-MET15, leu2Δ::P<sub>4Lacn.2</sub>_citrine_tCyc1-LEU2, ura3Δ::P<sub>RNR2</sub>_lacI(T167A,W220F,M254L, V119I,Q60G,M242L)-nls-linker-MBP-GST-nls_tCyc1-URA3* |  | Y3542 | P2772 | this work |
| 3723 | LacIm-MBP-Top7-nls | *met15Δ::P<sub>ACT1</sub>_lac12_tCyc1-MET15, leu2Δ::P<sub>4Lacn.2</sub>_citrine_tCyc1-LEU2, ura3Δ::P<sub>RNR2</sub>_lacI(T167A,W220F,M254L, V119I,Q60G,M242L)-nls-linker-MBP-top7-nls_tCyc1-URA3* |  | Y3542 | P2773 | this work |
| 3728 | P<sub>7tet.1</sub>_citrine | *P<sub>7tet.1</sub>_citrine-NATMX* |  | Y70 | P2776 | this work |
| 3746 | P<sub>7tet.1</sub>+P<sub>7SulA.1</sub> | *P<sub>7tet.1</sub>_citrine-NATMX, leu2Δ::P<sub>7SulA.1</sub>_mCardinal_tCyc1-LEU2* |  | Y3728 | P2774 | this work |
| 3764 | *WTC<sub>846</sub>::Citrine + P<sub>7SulA.1</sub>* | *P<sub>7tet.1</sub>_citrine-NATMX, leu2Δ::P<sub>7SulA.1</sub>_mCardinal_tCyc1-LEU2, ura3Δ::P<sub>RNR2</sub>_tetR-NLS-tup1_tCyc1_P<sub>7tet.1</sub>_tetR-NLS_tCyc1-URA3* |  | Y3746 | P2365 | this work |
| 3765 | P<sub>3Lacn.3</sub>-SR | *met15Δ::P<sub>ACT1</sub>_lac12_tCyc1-MET15, leu2Δ::P<sub>3Lacn.3</sub>_citrine_tCyc1-LEU2, his3Δ::P<sub>ACT1</sub>_lacI_tCyc1-HIS3* |  | Y3518 | P2601 | this work |
| 3807 | P<sub>4Lacn.2</sub>-LacImutTup1 | *met15Δ::P<sub>ACT1</sub>_lac12_tCyc1-MET15, leu2Δ::P<sub>4Lacn.2</sub>_citrine_tCyc1-LEU2, ura3Δ::P<sub>RNR2</sub>_lacI(T167A,W220F,M254L, V119I,Q60G,M242L)-nls-linker-tup1-nls_tCyc1-URA3* |  | Y3542 | P2798 | this work |
| 3808 | *WTC<sub>847</sub>::Citrine* | *met15Δ::P<sub>7SulA.1</sub>_citrine_tCyc1-MET15, lys2Δ::P<sub>ACT1</sub>_lexA-L7-hER(L387M)_tCyc1_P<sub>7SulA.1</sub>_lexA-hER-adh1tail_tCyc1-LYS2* |  | Y3391 | P2802 | this work |
| 3809 | *WTC<sub>847</sub>::* | *lys2Δ::P<sub>ACT1</sub>_lexA-L7-hER(L387M)_tCyc1_P<sub>7SulA.1</sub>_lexA-hER-adh1tail_tCyc1-LYS2* |  | Y70 | P2802 | this work |
| 3810 | *WTC<sub>846</sub>:: + WTC<sub>847</sub>::* | *ura3Δ::P<sub>RNR2</sub>_tetR-NLS-tup1_tCyc1_P<sub>7tet.1</sub>_tetR-NLS_tCyc1-URA3, lys2Δ::P<sub>ACT1</sub>_lexA-L7-hER(L387M)_tCyc1_P<sub>7SulA.1</sub>_lexA-hER-adh1tail_tCyc1-LYS2* |  | Y2761 | P2802 | this work |
| 3811 | *WTC<sub>846</sub>::Citrine + WTC<sub>846</sub>::mCardinal* | *P<sub>7tet.1</sub>_citrine-NATMX, leu2Δ::P<sub>7SulA.1</sub>_mCardinal_tCyc1-LEU2, ura3Δ::P<sub>RNR2</sub>_tetR-NLS-tup1_tCyc1_P<sub>7tet.1</sub>_tetR-NLS_tCyc1-URA3, lys2Δ::P<sub>ACT1</sub>_lexA-L7-hER(L387M)_tCyc1_P<sub>7SulA.1</sub>_lexA-hER-adh1tail_tCyc1-LYS2* |  | Y3764 | P2802 | this work |
| 3831 | *P<sub>4Lacn.2</sub>-LacImutNLS* | *met15Δ::P<sub>ACT1</sub>_lac12_tCyc1-MET15, leu2Δ::P<sub>4Lacn.2</sub>_citrine_tCyc1-LEU2, ura3Δ::P<sub>RNR2</sub>_lacI(T167A, W220F, M254L, V119I, Q60G, M242L)-nls_tCyc1-URA3* |  | Y3542 | P2804 | this work |
| 3832 | *P<sub>4Lacn.2</sub>-LacImut* | *met15Δ::P<sub>ACT1</sub>_lac12_tCyc1-MET15, leu2Δ::P<sub>4Lacn.2</sub>_citrine_tCyc1-LEU2, ura3Δ::P<sub>RNR2</sub>_lacI(T167A, W220F, M254L, V119I, Q60G, M242L)_tCyc1-URA3* |  | Y3542 | P2805 | this work |
| 3833 | *WTC<sub>847</sub>::TOR2* | *lys2Δ::P<sub>ACT1</sub>_lexA-L7-hER(L387M)_tCyc1_P<sub>7SulA.1</sub>_lexA-hER-adh1tail_tCyc1-LYS2, P<sub>7SulA.1</sub>_tor2_tTor2-NATMX* | AAGGGAA<br>AAGGGAA<br>A | Y3809 | P2754 | this work |
| 3834 | *WTC<sub>847</sub>::CDC20* | *lys2Δ::P<sub>ACT1</sub>_lexA-L7-hER(L387M)_tCyc1_P<sub>7SulA.1</sub>_lexA-hER-adh1tail_tCyc1-LYS2, P<sub>7SulA.1</sub>_cdc20_tCdc20-NATMX* | AAGGGAA<br>AAGGGAA<br>A | Y3809 | P2754 | this work |
| 3835 | *WTC<sub>847</sub>::IPL1* | *lys2Δ::P<sub>ACT1</sub>_lexA-L7-hER(L387M)_tCyc1_P<sub>7SulA.1</sub>_lexA-hER-adh1tail_tCyc1-LYS2, P<sub>7SulA.1</sub>_ipl1_tIpl1-NATMX* | AAGGGAA<br>AAGGGAA<br>A | Y3809 | P2754 | this work |
| 3836 | *WTC<sub>847</sub>::PGK1* | *lys2Δ::P<sub>ACT1</sub>_lexA-L7-hER(L387M)_tCyc1_P<sub>7SulA.1</sub>_lexA-hER-adh1tail_tCyc1-LYS2, P<sub>7SulA.1</sub>_pgk1_tPgk1-NATMX* | AAGGGAA<br>AAGGGAA<br>A | Y3809 | P2754 | this work |
| 3837 | *WTC<sub>847</sub>::TPI1* | *lys2Δ::P<sub>ACT1</sub>_lexA-L7-hER(L387M)_tCyc1_P<sub>7SulA.1</sub>_lexA-hER-adh1tail_tCyc1-LYS2, P<sub>7SulA.1</sub>_tpi1_tTpi1-NATMX* | AAGGGAA<br>AAGGGAA<br>A | Y3809 | P2754 | this work |
| 3839 | *WTC<sub>847</sub>::KRE5* | *lys2Δ::P<sub>ACT1</sub>_lexA-L7-hER(L387M)_tCyc1_P<sub>7SulA.1</sub>_lexA-hER-adh1tail_tCyc1-LYS2, P<sub>7SulA.1</sub>_kre5_tKre5-NATMX* | AAGGGAA<br>AAGGGAA<br>A | Y3809 | P2754 | this work |
| 3840 | *WTC<sub>847</sub>::KRE9* | *lys2Δ::P<sub>ACT1</sub>_lexA-L7-hER(L387M)_tCyc1_P<sub>7SulA.1</sub>_lexA-hER-adh1tail_tCyc1-LYS2, P<sub>7SulA.1</sub>_kre9_tKre9-NATMX* | AAGGGAA<br>AAGGGAA<br>A | Y3809 | P2754 | this work |
| 3841 | WTC<sub>847</sub>::CDC28 | *lys2Δ::P<sub>ACT1</sub>_lexA-L7-hER(L387M)_tCyc1_P<sub>7SulA.1</sub>_lexA-hER-adh1tail_tCyc1-LYS2, P<sub>7SulA.1</sub>_cdc28_tCdc28-NATMX* | AAGGGAA<br>AAGGGAA<br>A | Y3809 | P2754 | this<br>work |
| 3843 | WTC<sub>847</sub>::PBR1 | *lys2Δ::P<sub>ACT1</sub>_lexA-L7-hER(L387M)_tCyc1_P<sub>7SulA.1</sub>_lexA-hER-adh1tail_tCyc1-LYS2, P<sub>7SulA.1</sub>_pbr1_tPbr1-NATMX* | AAGGGAA<br>AAGGGAA<br>A | Y3809 | P2754 | this<br>work |
| 3844 | WTC<sub>847</sub>::CDC42 | *lys2Δ::P<sub>ACT1</sub>_lexA-L7-hER(L387M)_tCyc1_P<sub>7SulA.1</sub>_lexA-hER-adh1tail_tCyc1-LYS2, P<sub>7SulA.1</sub>_cdc42_tCdc42-NATMX* | AAGGGAA<br>AAGGGAA<br>A | Y3809 | P2754 | this<br>work |
| 3845 | WTC<sub>847</sub>::PMA1 | *lys2Δ::P<sub>ACT1</sub>_lexA-L7-hER(L387M)_tCyc1_P<sub>7SulA.1</sub>_lexA-hER-adh1tail_tCyc1-LYS2, P<sub>7SulA.1</sub>_pma1_tPma1-NATMX* | AAGGGAA<br>AAGGGAA<br>A | Y3809 | P2754 | this<br>work |
| 3847 | WTC<sub>847</sub>::FBA1 | *lys2Δ::P<sub>ACT1</sub>_lexA-L7-hER(L387M)_tCyc1_P<sub>7SulA.1</sub>_lexA-hER-adh1tail_tCyc1-LYS2, P<sub>7SulA.1</sub>_fba1_tFba1-NATMX* | AAGGGAA<br>AAGGGAA<br>A | Y3809 | P2754 | this<br>work |
| 3849 | LexA-L7-hER-Tup1 | *met15Δ::P<sub>7SulA.1</sub>_citrine_tCyc1-MET15, lys2Δ::P<sub>ACT1</sub>_lexA-L7-hER-tup1_tCyc1-LYS2* |  | Y3391 | P2806 | this<br>work |
| 3856 | WTC<sub>848</sub>::Citrine | *met15Δ::P<sub>ACT1</sub>_lac12_tCyc1-MET15, leu2Δ::P<sub>4Lacn.2</sub>_citrine_tCyc1-LEU2, his3Δ::P<sub>Rnr2</sub>_LacI(T167A,W220F,M254L,V119I,Q60G,M242L,NLS)_tFum1_P<sub>4Lacn.2</sub>_LacI(T167A,W220F,M254L,Q60G,M242L)_tCyc1* |  | Y3542 | P2814 | this<br>work |
| 3870 | WTC<sub>848</sub>:: | *met15Δ::P<sub>ACT1</sub>_lac12_tCyc1-MET15, his3Δ::P<sub>RNR2</sub>_LacI(T167A,W220F,M254L,V119I,Q60G,M242L,NLS)_tFum1_P<sub>4Lacn.2</sub>_LacI(T167A,W220F,M254L,Q60G,M242L)_tCyc1-HIS3* |  | Y3316 | P2814 | this<br>work |
| 3873 | WTC<sub>846</sub>::Citrine +<br>WTC<sub>846</sub>::mCardinal +<br>Lac12 | *P<sub>7tet.1</sub>_citrine-NATMX, leu2Δ::P<sub>7SulA.1</sub>_mCardinal_tCyc1-LEU2, ura3Δ::P<sub>RNR2</sub>_tetR-NLS-tup1_tCyc1_P<sub>7tet.1</sub>_tetR-NLS_tCyc1-URA3, lys2Δ::P<sub>ACT1</sub>_lexA-L7-hER(L387M)_tCyc1_P<sub>7SulA.1</sub>_lexA-hER-adh1tail_tCyc1-LYS2, met15Δ::P<sub>ACT1</sub>_lac12_tCyc1-MET15* |  | Y3811 | P2072 | this<br>work |
| 3874 | WTC<sub>846</sub>:: + WTC<sub>847</sub>:: +<br>Lac12 | *ura3Δ::P<sub>RNR2</sub>_tetR-NLS-tup1_tCyc1_P<sub>7tet.1</sub>_tetR-NLS_tCyc1-URA3, lys2Δ::P<sub>ACT1</sub>_lexA-L7-hER(L387M)_tCyc1_P<sub>7SulA.1</sub>_lexA-hER-adh1tail_tCyc1-LYS2, met15Δ::P<sub>ACT1</sub>_lac12_tCyc1-MET15* |  | Y3810 | P2072 | this<br>work |
| 3877 | WTC<sub>846</sub>::Citrine +<br>WTC<sub>846</sub>::mCardinal +<br>WTC<sub>848</sub>:: | *P<sub>7tet.1</sub>_citrine-NATMX, leu2Δ::P<sub>7SulA.1</sub>_mCardinal_tCyc1-LEU2, ura3Δ::P<sub>RNR2</sub>_tetR-NLS-tup1_tCyc1_P<sub>7tet.1</sub>_tetR-NLS_tCyc1-URA3,lys2Δ::P<sub>ACT1</sub>_lexA-L7-hER(L387M)_tCyc1_P<sub>7SulA.1</sub>_lexA-hER-adh1tail_tCyc1-LYS2, met15Δ::P<sub>ACT1</sub>_lac12_tCyc1-MET15, his3Δ::P<sub>RNR2</sub>_LacI(T167A,W220F,M254L,V119I,Q60G,M242L,NLS)_tFum1_P<sub>4Lacn.2</sub>_LacI(T167A,W220F,M254L,Q60G,M242L)_tCyc1-HIS3* |  | Y3873 | P2814 | this work |
| 3878 | WTC<sub>846</sub>:: + WTC<sub>847</sub>:: +<br>WTC<sub>848</sub>:: | *ura3Δ::P<sub>RNR2</sub>_tetR-NLS-tup1_tCyc1_P<sub>7tet.1</sub>_tetR-NLS_tCyc1-URA3, lys2Δ::P<sub>ACT1</sub>_lexA-L7-hER(L387M)_tCyc1_P<sub>7SulA.1</sub>_lexA-hER-adh1tail_tCyc1-LYS2, met15Δ::P<sub>ACT1</sub>_lac12_tCyc1-MET15,his3Δ::P<sub>RNR2</sub>_LacI(T167A,W220F,M254L,V119I,Q60G,M242L,NLS)_tFum1_P<sub>4Lacn.2</sub>_LacI(T167A,W220F,M254L,Q60G,M242L)_tCyc1-HIS3* |  | Y3874 | P2814 | this work |
| 3880 | WTC<sub>846</sub>:: + WTC<sub>847</sub>:: +<br>WTC<sub>848</sub>::Citrine | *ura3Δ::P<sub>RNR2</sub>_tetR-NLS-tup1_tCyc1_P<sub>7tet.1</sub>_tetR-NLS_tCyc1-URA3, lys2Δ::P<sub>ACT1</sub>_lexA-L7-hER(L387M)_tCyc1_P<sub>7SulA.1</sub>_lexA-hER-adh1tail_tCyc1-LYS2, met15Δ::P<sub>ACT1</sub>_lac12_tCyc1-MET15,his3Δ::P<sub>RNR2</sub>_LacI(T167A,W220F,M254L,V119I,Q60G,M242L,NLS)_tFum1_P<sub>4Lacn.2</sub>_LacI(T167A,W220F,M254L,Q60G,M242L)_tCyc1-HIS3, leu2Δ::P<sub>4Lacn.2</sub>_citrine_tCyc1-LEU2* |  | Y3878 | P2667 | this work |
| 3924 | WTC<sub>848</sub>::KRE9 | *met15Δ::P<sub>ACT1</sub>_lac12_tCyc1-MET15, his3Δ::P<sub>RNR2</sub>_LacI(T167A,W220F,M254L,V119I,Q60G,M242L,NLS)_tFum1_P<sub>4Lacn.2</sub>_LacI(T167A,W220F,M254L,Q60G,M242L)_tCyc1-HIS3, P<sub>4Lacn.2</sub>_kre9_tKre9-NATMX* | AAGGGAA<br>AAGGGAA<br>A | Y3870 | P2765 | this work |
| 3925 | WTC<sub>846</sub>::CDC14 +<br>WTC<sub>847</sub>:: + WTC<sub>848</sub>:: | *ura3Δ::P<sub>RNR2</sub>_tetR-NLS-tup1_tCyc1_P<sub>7tet.1</sub>_tetR-NLS_tCyc1-URA3, lys2Δ::P<sub>ACT1</sub>_lexA-L7-hER(L387M)_tCyc1_P<sub>7SulA.1</sub>_lexA-hER-adh1tail_tCyc1-LYS2, met15Δ::P<sub>ACT1</sub>_lac12_tCyc1-MET15,his3Δ::P<sub>RNR2</sub>_LacI(T167A,W220F,M254L,V119I,Q60G,M242L,NLS)_tFum1_P<sub>4Lacn.2</sub>_LacI(T167A,W220F,M254L,Q60G,M242L)_tCyc1-HIS3, P<sub>7tet.1</sub>_cdc14_tCdc14-NATMX* | AAGGGAA<br>AAGGGAA<br>A | Y3878 | P2375 | this work |
| 3926 | WTC<sub>848</sub>::KRE5 | *met15Δ::P<sub>ACT1</sub>_lac12_tCyc1-MET15, his3Δ::P<sub>RNR2</sub>_LacI(T167A,W220F,M254L,V119I,Q60G,M242L,NLS)_tFum1_P<sub>4Lacn.2</sub>_LacI(T167A,W220F,M254L,Q60G,M242L)_tCyc1-HIS3, P<sub>4Lacn.2</sub>_kre5_tKre5-NATMX* | AAGGGAA<br>AAGGGAA<br>A | Y3870 | P2765 | this work |
| 3928 | WTC<sub>846</sub>::CDH1 +<br>WTC<sub>847</sub>:: + WTC<sub>848</sub>:: | *ura3Δ::P<sub>RNR2</sub>_tetR-NLS-tup1_tCyc1_P<sub>7tet.1</sub>_tetR-NLS_tCyc1-URA3, lys2Δ::P<sub>ACT1</sub>_lexA-L7-hER(L387M)_tCyc1_P<sub>7SulA.1</sub>_lexA-hER-adh1tail_tCyc1-LYS2, met15Δ::P<sub>ACT1</sub>_lac12_tCyc1-MET15,his3Δ::P<sub>RNR2</sub>_LacI(T167A,W220F,M254L,V119I,Q60G,M242L,NLS)_tFum1_P<sub>4Lacn.2</sub>_LacI(T167A,W220F,M254L,Q60G,M242L)_tCyc1-HIS3, P<sub>7tet.1</sub>_cdh1_tCdh1-NATMX* | AAGGGAA<br>AAGGGAA<br>A | Y3878 | P2375 | this work |
| 3929 | WTC<sub>848</sub>::ESP1 | *met15Δ::P<sub>ACT1</sub>_lac12_tCyc1-MET15, his3Δ::P<sub>RNR2</sub>_LacI(T167A,W220F,M254L,V119I,Q60G,M242L,NLS)_tFum1_P<sub>4Lacn.2</sub>_LacI(T167A,W220F,M254L,Q60G,M242L)_tCyc1-HIS3, P<sub>4Lacn.2</sub>_esp1_tEsp1-NATMX* | AAGGGAA<br>AAGGGAA<br>A | Y3870 | P2765 | this work |
| 3932 | *WTC<sub>846</sub>:: + WTC<sub>847</sub>:: + WTC<sub>848</sub>::NET1* | *ura3Δ::P<sub>RNR2</sub>_tetR-NLS-tup1_tCyc1_P<sub>7tet.1</sub>_tetR-NLS_tCyc1-URA3, lys2Δ::P<sub>ACT1</sub>_lexA-L7-hER(L387M)_tCyc1_P<sub>7SulA.1</sub>_lexA-hER-adh1tail_tCyc1-LYS2, met15Δ::P<sub>ACT1</sub>_lac12_tCyc1-MET15, his3Δ::P<sub>RNR2</sub>_LacI(T167A, W220F, M254L, V119I, Q60G, M242L, NLS)_tFum1_P<sub>4Lacn.2</sub>_LacI(T167A, W220F, M254L, Q60G, M242L)_tCyc1-HIS3, P<sub>4Lacn.2</sub>_net1_tNet1* | AAGGGAA<br>AAGGGAA<br>A | Y3878 | P2765 | this<br>work |
| 3940 | *WTC<sub>848</sub>::PMA1* | *met15Δ::P<sub>ACT1</sub>_lac12_tCyc1-MET15, his3Δ::P<sub>RNR2</sub>_LacI(T167A, W220F, M254L, V119I, Q60G, M242L, NLS)_tFum1_P<sub>4Lacn.2</sub>_LacI(T167A, W220F, M254L, Q60G, M242L)_tCyc1-HIS3, P<sub>4Lacn.2</sub>_pma1_tPma1-NATMX* | AAGGGAA<br>AAGGGAA<br>A | Y3870 | P2765 | this<br>work |
| 3941 | *WTC<sub>848</sub>::WHI5* | *met15Δ::P<sub>ACT1</sub>_lac12_tCyc1-MET15, his3Δ::P<sub>RNR2</sub>_LacI(T167A, W220F, M254L, V119I, Q60G, M242L, NLS)_tFum1_P<sub>4Lacn.2</sub>_LacI(T167A, W220F, M254L, Q60G, M242L)_tCyc1-HIS3, P<sub>4Lacn.2</sub>_whi5_tWhi5-NATMX* | AAGGGAA<br>AAGGGAA<br>A | Y3870 | P2765 | this<br>work |
| 3942 | *WTC<sub>848</sub>::IPL1* | *met15Δ::P<sub>ACT1</sub>_lac12_tCyc1-MET15, his3Δ::P<sub>RNR2</sub>_LacI(T167A, W220F, M254L, V119I, Q60G, M242L, NLS)_tFum1_P<sub>4Lacn.2</sub>_LacI(T167A, W220F, M254L, Q60G, M242L)_tCyc1-HIS3, P<sub>4Lacn.2</sub>_ipl1_tIpl1-NATMX* | AAGGGAA<br>AAGGGAA<br>A | Y3870 | P2765 | this<br>work |
| 3943 | *WTC<sub>848</sub>::CLN3* | *met15Δ::P<sub>ACT1</sub>_lac12_tCyc1-MET15, his3Δ::P<sub>RNR2</sub>_LacI(T167A, W220F, M254L, V119I, Q60G, M242L, NLS)_tFum1_P<sub>4Lacn.2</sub>_LacI(T167A, W220F, M254L, Q60G, M242L)_tCyc1-HIS3, P<sub>4Lacn.2</sub>_cln3_tCln3-NATMX* | AAGGGAA<br>AAGGGAA<br>A | Y3870 | P2765 | this<br>work |
| 3944 | *WTC<sub>848</sub>::CDC20* | *met15Δ::P<sub>ACT1</sub>_lac12_tCyc1-MET15, his3Δ::P<sub>RNR2</sub>_LacI(T167A, W220F, M254L, V119I, Q60G, M242L, NLS)_tFum1_P<sub>4Lacn.2</sub>_LacI(T167A, W220F, M254L, Q60G, M242L)_tCyc1-HIS3, P<sub>4Lacn.2</sub>_cdc20_tCdc20-NATMX* | AAGGGAA<br>AAGGGAA<br>A | Y3870 | P2765 | this<br>work |
| 3945 | *WTC<sub>846</sub>::CDC14 + WTC<sub>847</sub>:: + WTC<sub>848</sub>::NET1* | *ura3Δ::P<sub>RNR2</sub>_tetR-NLS-tup1_tCyc1_P<sub>7tet.1</sub>_tetR-NLS_tCyc1-URA3, lys2Δ::P<sub>ACT1</sub>_lexA-L7-hER(L387M)_tCyc1_P<sub>7SulA.1</sub>_lexA-hER-adh1tail_tCyc1-LYS2, met15Δ::P<sub>ACT1</sub>_lac12_tCyc1-MET15, his3Δ::P<sub>RNR2</sub>_LacI(T167A, W220F, M254L, V119I, Q60G, M242L, NLS)_tFum1_P<sub>4Lacn.2</sub>_LacI(T167A, W220F, M254L, Q60G, M242L)_tCyc1-HIS3, P<sub>7tet.1</sub>_cdc14_tCdc14-NATMX, P<sub>4Lacn.2</sub>_net1_tNet1* | AAGGGAA<br>AAGGGAA<br>A | Y3925 | P2765 | this<br>work |
| 3947 | *WTC<sub>846</sub>::CDH1 + WTC<sub>847</sub>:: + WTC<sub>848</sub>::PDS1* | *ura3Δ::P<sub>RNR2</sub>_tetR-NLS-tup1_tCyc1_P<sub>7tet.1</sub>_tetR-NLS_tCyc1-URA3, lys2Δ::P<sub>ACT1</sub>_lexA-L7-hER(L387M)_tCyc1_P<sub>7SulA.1</sub>_lexA-hER-adh1tail_tCyc1-LYS2, met15Δ::P<sub>ACT1</sub>_lac12_tCyc1-MET15, his3Δ::P<sub>RNR2</sub>_LacI(T167A, W220F, M254L, V119I, Q60G, M242L, NLS)_tFum1_P<sub>4Lacn.2</sub>_LacI(T167A, W220F, M254L, Q60G, M242L)_tCyc1-HIS3, P<sub>7tet.1</sub>_cdh1_tCdh1-NATMX, P<sub>4Lacn.2</sub>_pds1_tPds1* | AAGGGAA<br>AAGGGAA<br>A | Y3928 | P2765 | this<br>work |
| 3979 | WTC<sub>846</sub>::CDH1 +<br>WTC<sub>847</sub>::ESP1 +<br>WTC<sub>848</sub>::PDS1 | ura3Δ::P<sub>RNR2</sub>_tetR-NLS-tup1_tCyc1_P<sub>7tet.1</sub>_tetR-NLS_tCyc1-URA3, lys2Δ::P<sub>ACT1</sub>_lexA-L7-hER(L387M)_tCyc1_P<sub>7SulA.1</sub>_lexA-hER-adh1tail_tCyc1-LYS2, met15Δ::P<sub>ACT1</sub>_lac12_tCyc1-MET15,his3Δ::P<sub>RNR2</sub>_LacI(T167A,W220F,M254L,V119I,Q60G,M242L,NLS)_tFum1_P<sub>4Lacn.2</sub>_LacI(T167A,W220F,M254L,Q60G,M242L)_tCyc1-HIS3, P<sub>7tet.1</sub>_cdh1_tCdh1-NATMX, P<sub>4Lacn.2</sub>_pds1_tPds1, P<sub>7SulA.1</sub>_esp1_tEsp1 | AAGGGAA<br>AAGGGAA<br>A | Y3947 | P2755 | this<br>work |

**Table 5.**
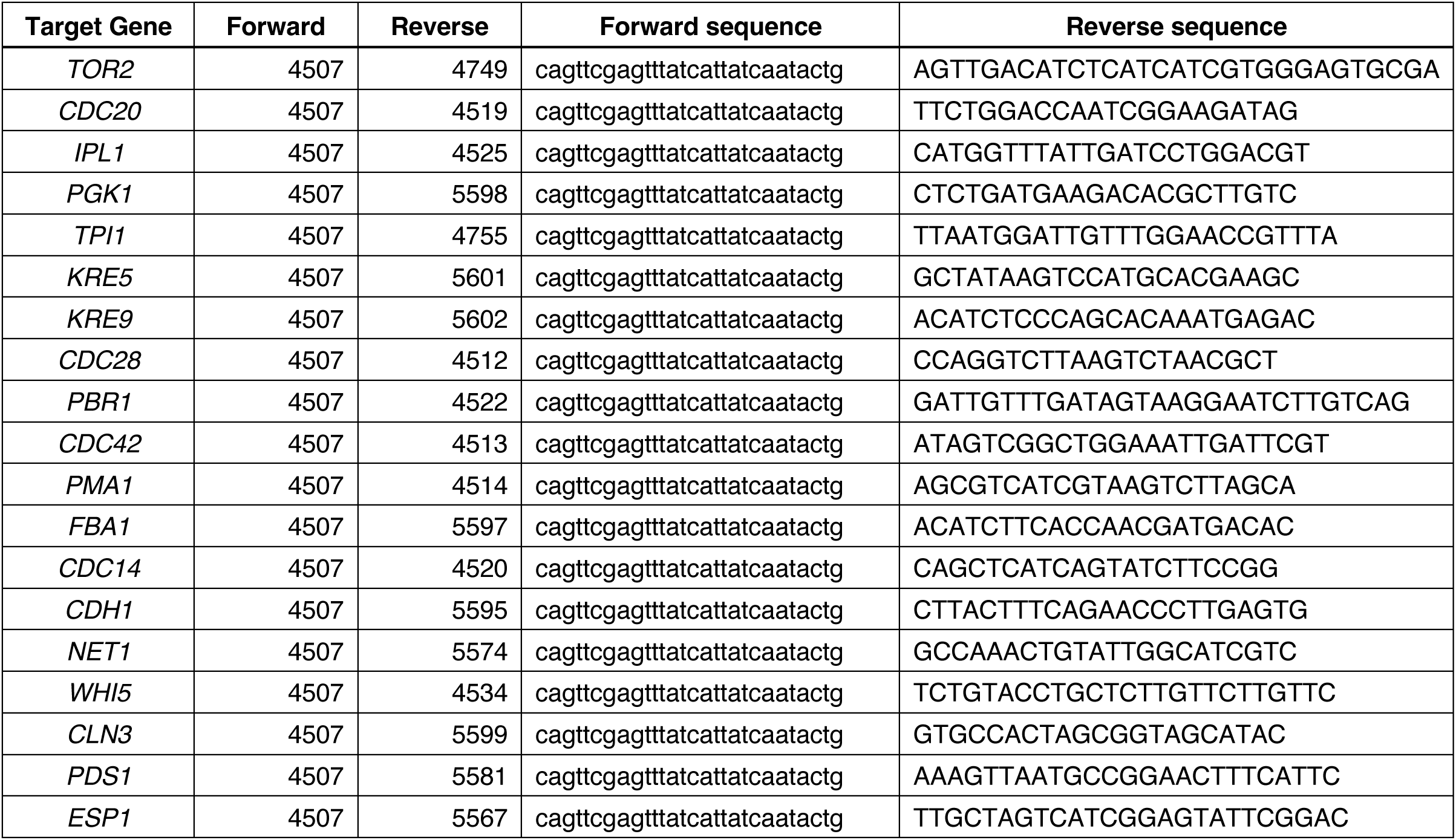
Colony PCR oligos used to check correct replacement of the endogenous promoter with either P7SulA.1 or P4Lacn.2.

### Chemicals and Media

All media compositions as described before^19^. The inducers used were purchased from Cayman Chemicals (aTc, 10009542) or Sigma Aldrich (IPTG I6758, ß-estradiol E2758). aTc was prepared as a 4628.8ng/mL (10mM) stock solution in ethanol. ß-estradiol was prepared as 10mM stock in 70% ethanol. IPTG was prepared as 1M stock in double distilled, sterile water. All were kept at −20°C for long term storage and were diluted in water or media for experiments as necessary.

When constructing strains where WTC promoters replaced endogenous gene promoters, the CRISPR-Cas9 protocol was used as previously explained^52^. In cases where the antibiotic selection marker NATMX was integrated together with the promoter, Nourseothricin (Werner BioAgents, clonNAT) was present in the YPD selection plate, along with the relevant inducer (IPTG or ß-estradiol).

### Spotting Assay

For spotting assays, all strains were cultured overnight in YPD with the lowest amount of IPTG or ß-estradiol possible that still achieved wild type growth rates. In the morning cells were diluted into fresh YPD at a concentration of 0.8 million cells/mL and grown at 30°C for a further 8 hours. The cells were then collected by centrifugation and resuspended in fresh YPD. They were spotted onto prepared agar plates with various inducer concentrations, such that the most concentrated spot contained 100.000 cells per mL and each subsequent spot was a 1:10 dilution. Pictures of the plates were taken after incubation at 30°C for 24 hours.

### Flow cytometry

Cells were grown overnight in rich media and diluted 1:200 into experimental conditions in the afternoon. The next morning, they were again diluted 1:25 into the same conditions and grown for at least a further 6 hours before measurement. Samples were diluted in PBS and measured using a LSRFortessa LSRII equipped with a high-throughput sampler. PMT voltages for the forward and side scatter measurements were set up such that the height of the signal was not saturated.

Citrine fluorescence was quantified using a 488 nm excitation laser and a 530/30 nm emission filter. mCardinal fluorescence was quantified using a 561 nm excitation laser and a 710/50 emission filter. Channel PMT voltages were set up such that a strain expressing Citrine from P*TDH3* did not saturate the device. In cases where basal expression was measured, the PMT voltage was set to maximum to observe small differences in basal expression levels.

### Liquid Growth Assays

Cells were grown overnight at 30°C in the same media the experiment was to be performed in. They were diluted into 96-well plates with transparent bottoms (Enzyscreen, CR1496dg) with various inducer concentrations where appropriate. The total liquid volume in each well was 250µL. Plates were covered and shaken (250rpm) at 30°C by the Growth Profiler 360. Pictures of the bottom of the plate were taken every 20 or 30 minutes depending on the experimental setup. Growth Profiler Viewer 360 software was used to automatically quantify culture density in each well from these pictures. Amplitude values correspond to this calculated value minus the lowest amplitude recorded in a given well. This correction sets the lowest recorded value as 0.

## Data analysis and availability

All analysis of flow cytometry data, dose response curve fittings and calculation of the cell-to-cell variation measure were performed as explained before^19^. Slopes of growth curves were calculated as 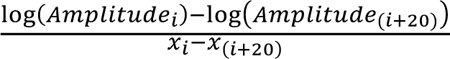 where *x*_i_ is the time at which a measurement was taken. For example, where measurements are 20 minutes apart, these two measurements would be separated by 400 minutes. Maximum slope recorded over the growth curve is reported as the Growth Rate and is used as a proxy for fitness. Key plasmids can be ordered through Addgene. All other plasmids, strains and raw data can be requested through the author.

