## Supplementary Figures for "Orthogonal Inducible Transcriptional Control Systems for Investigation of Gene-Gene Interactions"

Aslı Azizoğlu[1]

[1]D-BSSE and Swiss Institute of Bioinformatics, ETH Zurich, Basel, Switzerland

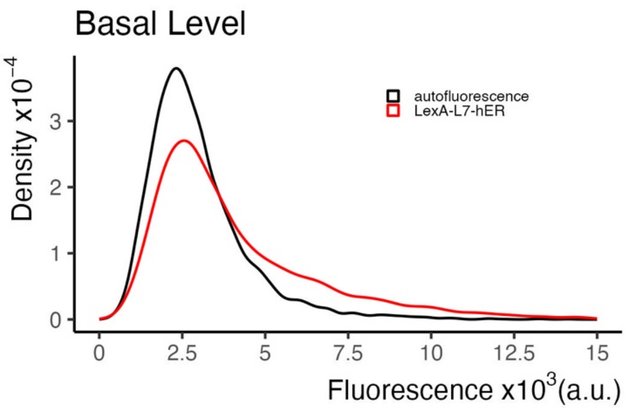


**Figure S1. LexA-hER can repress gene expression from $P_{7SulA.1}$.** A strain bearing $P_{7SulA.1}$ driven Citrine and $P_{ACT1}$ driven LexA-hER was grown to exponential phase in presence of 10 $\mu$M β-estradiol. Citrine fluorescence was measured using flow cytometry (n=5000 cells per measurement). The entire population of measured cells are plotted as a density plot, where the area under the curve equals 1 and the y axis indicates the proportion of cells with a given value of fluorescence. Autofluorescence refers to strain Y70 without any integrated constructs.

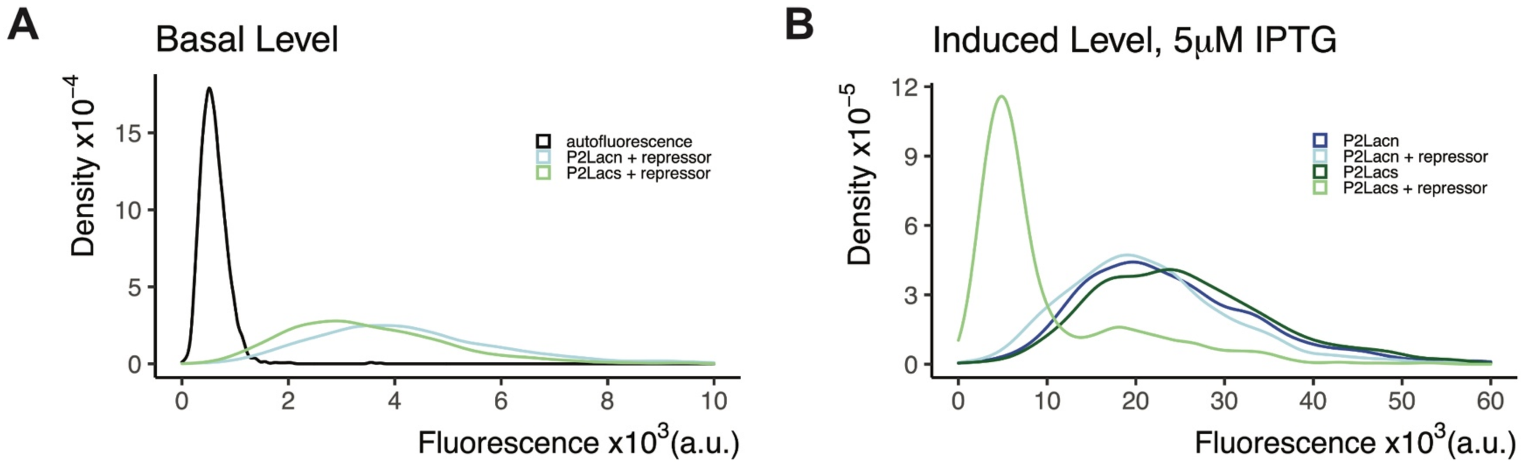


**Figure S2. Effect of different LacI binding sites on basal level and inducibility of the promoter. A)** Two promoters, each with two LacI binding sites flanking their TATA box were genomically integrated to drive Citrine fluorescent protein expression. $P_{2Lacn}$ had *Lacn* sites and $P_{2Lacs}$ had *Lacs* sites, as described in the text. In these strains (Y3420,3421), both the repressor LacI and the membrane IPTG transporter Lac12 were expressed from genomically integrated $P_{ACT1}$. Citrine fluorescence was measured from these strains during exponential phase with flow cytometry (n=5000 cells) and compared to the parent strain Y70 as autofluorescence control to determine basal level. No IPTG was present in these cultures. **B)** Strains Y3420, 3421 from (A) and two strains where the repressor LacI was not present (Y3412,3413) were grown in presence of 5 $\mu$M IPTG and Citrine fluorescence was measured in the same fashion as (A). Plots are density plots, where the area under the curve equals 1 and the y axis indicates the proportion of cells with a given value of fluorescence.

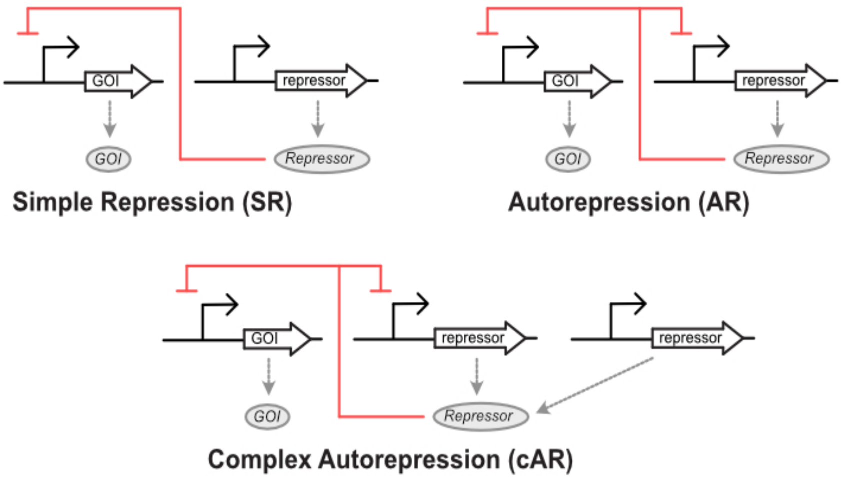


**Figure S3. Schematic representation of the cAR architecture of the WTC systems.** The large empty arrows indicate genes and the region before them represent promoters. Both the Gene of Interest (GOI) and the repressor are transcribed and translated, as represented by the dashed arrows. The red lines represent the relationship between the repressor protein and the promoters within the system. In Simple Repression, the Repressor can repress only the promoter of the GOI. In Autorepression, the repressor represses transcription of both itself and the GOI. These two architectures are combined to make the cAR architecture, in which two separate instances of the repressor gene are present. The repressor protein then represses transcription of the GOI and one of the repressor genes.

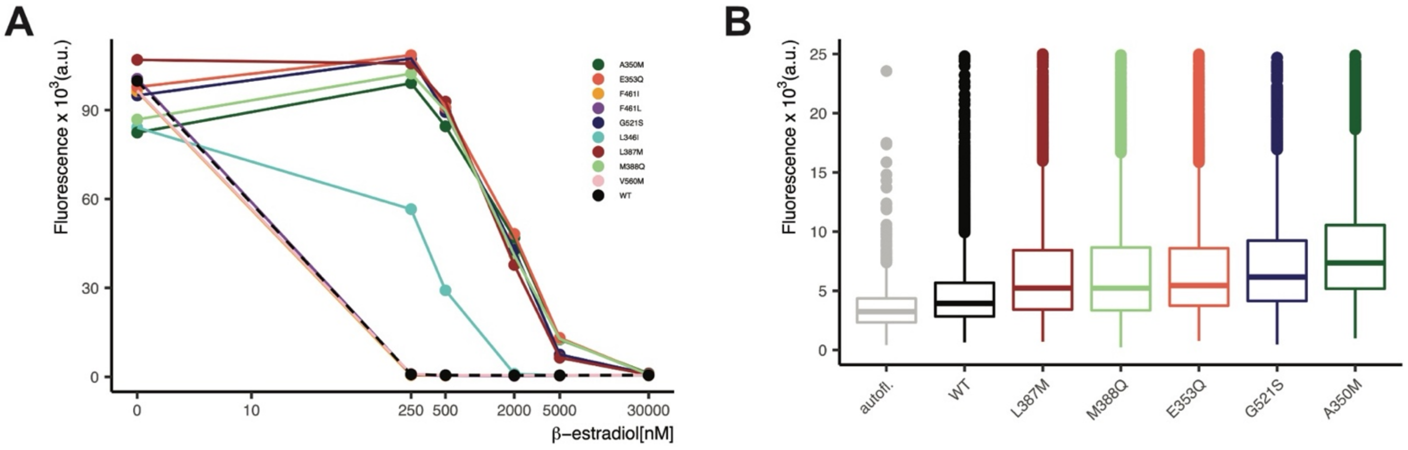


**Figure S4. Reducing β-estradiol sensitivity of LexA-hER through single amino acid mutations. A)** Single amino acid mutations were introduced into hER ligand binding domain (aa 282-595 of full length hER) found in the fusion protein LexA-hER. The mutations are indicated in the legend and the numbers correspond to the amino acid mutated with reference to the full hER sequence. These mutated LexA-hER versions and an unmutated version (WT) were expressed from $P_{ACT1}$. They were genomically integrated into the strain Y3391, which also contains a genomically integrated $P7_{SulA.1}$_citrine construct. These strains (Y3560,3561,3626,3627,3629-33, 3516) were grown overnight in the indicated β-estradiol concentrations, followed by dilution into the same conditions in the morning. Citrine fluorescence was measured using flow cytometry after 6 hours and during exponential phase (n=5000 cells per strain per dose). The symbols indicate median fluorescence measured for each dose. **B)** Strains with right-shifted dose response curves compared to the WT strain were selected and their basal expression with 30$\mu$M β-estradiol was measured with flow cytometry (n=5000 cells per strain). The sensitivity of the flow cytometer was increased to maximum for these measurements. A strain without any integrated constructs was also measured as autofluorescence control (Y70, autofl.). The boxplots span from first to third quartile and the line within the box indicates the median of the measured fluorescence.

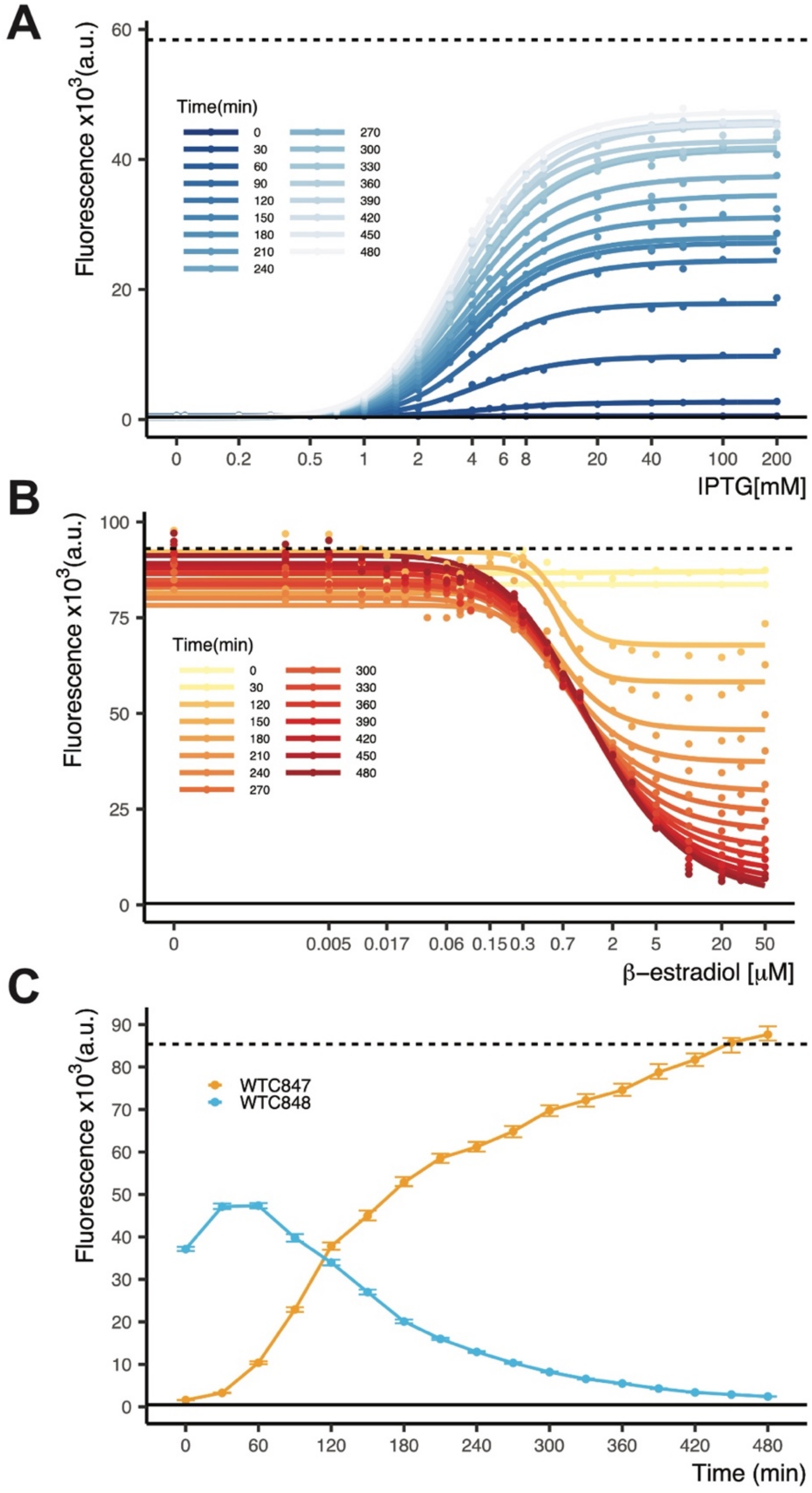


**Figure S5. Time dependent dose response behaviour of $WTC_{847}$ and $WTC_{848}$. A)** A strain bearing $P_{4Lacn.2}$ expressed Citrine and all $WTC_{848}$ components (*$WTC_{848}$::Citrine*, Y3856) was grown in YPD with different IPTG concentrations. Samples were taken right before inducer addition (t=0) and every 30 minutes thereafter. Citrine fluorescence was measured using flow cytometry (n=5000 cells per dose). Symbols indicate the median fluorescence of the measured population at each dose. Lines are fitted using a five-parameter log logistic function as explained in Methods. Dashed line indicates autofluorescence signal measured from the parental strain without Citrine (Y3316). Dot-dashed line indicates fluorescence measured from a strain where $P_{4Lacn.2}$ drove Citrine expression (Y3542). **B)** A strain bearing $P_{7SulA.1}$ expressed Citrine and all $WTC_{847}$ components (*$WTC_{847}$::Citrine*, Y3808) was grown in YPD with different β-estradiol concentrations. Samples were taken right before inducer addition (t=0) and every 30 minutes thereafter apart from 60 and 90 minutes. Citrine fluorescence was measured using flow cytometry (n=5000 cells per dose). Symbols and lines as in (A). Dashed line indicates autofluorescence signal measured from the parental strain without Citrine (Y70). Dot-dashed line indicates fluorescence measured from a strain where $P_{7SulA.1}$ drove Citrine expression (Y3391). **C)** *$WTC_{847}$::Citrine* and *$WTC_{848}$::Citrine* strains were grown in YPD with maximum inducer (50$\mu$M β-estradiol/ 50mM IPTG) overnight. At time 0, all inducer was removed by washing the culture with YPD. Citrine fluorescence was measured every 30 minutes by flow cytometry (n=5000 cells per time point). Dashed line indicates fluorescence measured from a strain where $P_{7SulA.1}$ drove Citrine expression (Y3391) and solid line indicates the parental strain (Y70). Error bars indicate 95% confidence interval calculated using bootstrapping (n=1000) as described in Methods.

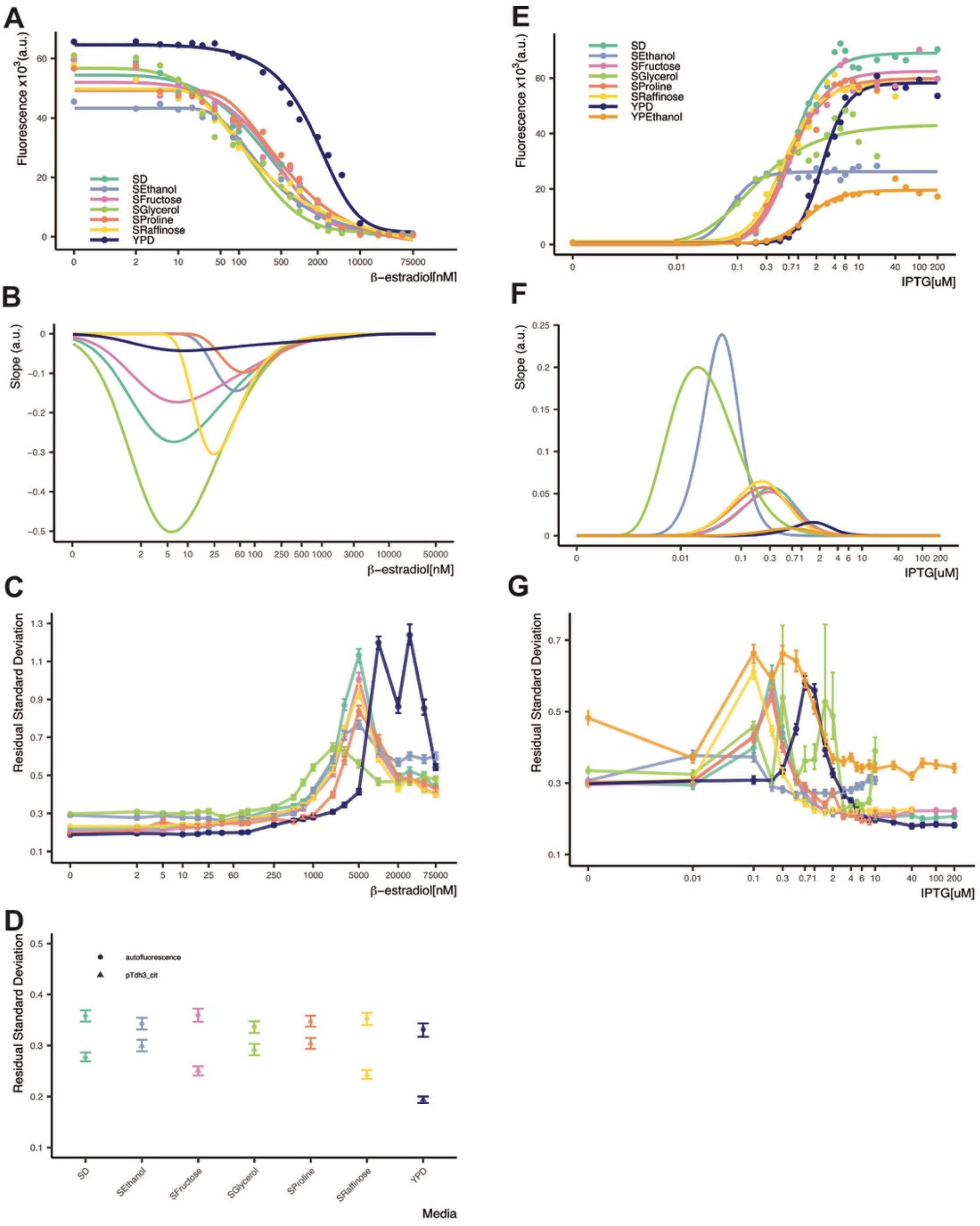


**Figure S6. Behaviour of $WTC_{847}$ and $WTC_{848}$ under different media conditions.** Cells bearing either **A)** $WTC_{847}$ (Y3808) or **E)** $WTC_{848}$ (Y3856) controlled Citrine were grown in different media conditions. Citrine fluorescence was measured with flow cytometry while the cells were in early to mid-exponential phase (n=5000 cells per dose). Symbols indicate the median fluorescence of the measured population at each dose. Lines are fitted using a five-parameter log logistic function as explained in Methods. **B)** Slopes of the dose response curves shown in (A). **C)** Cell-to-cell variation of expression in the strains shown in (A). Higher Residual Standard Deviation (RSD) values (y axis) correspond to greater variation in expression, calculated as explained in the text and previously in [1]. **D)** Cell-to-cell variation seen in two control strains. Autofluorescence refers to a strain without any integrated constructs (Y70) and pTdh3_cit refers to a strain where Citrine is constitutively expressed from the *TDH3* promoter (Y2683). **F)** Slopes of the dose response curves shown in (E). **G)** Cell-to-cell variation of expression in the strains shown in (E). Dot-dash line indicates the variation of the strain where Citrine is constitutively expressed from $P_{7SulA.1}$ and dashed line indicates variation of autofluorescence in the parent strain without Citrine. Error bars indicate 95% confidence interval calculated using bootstrapping (n=1000) as described in Methods.

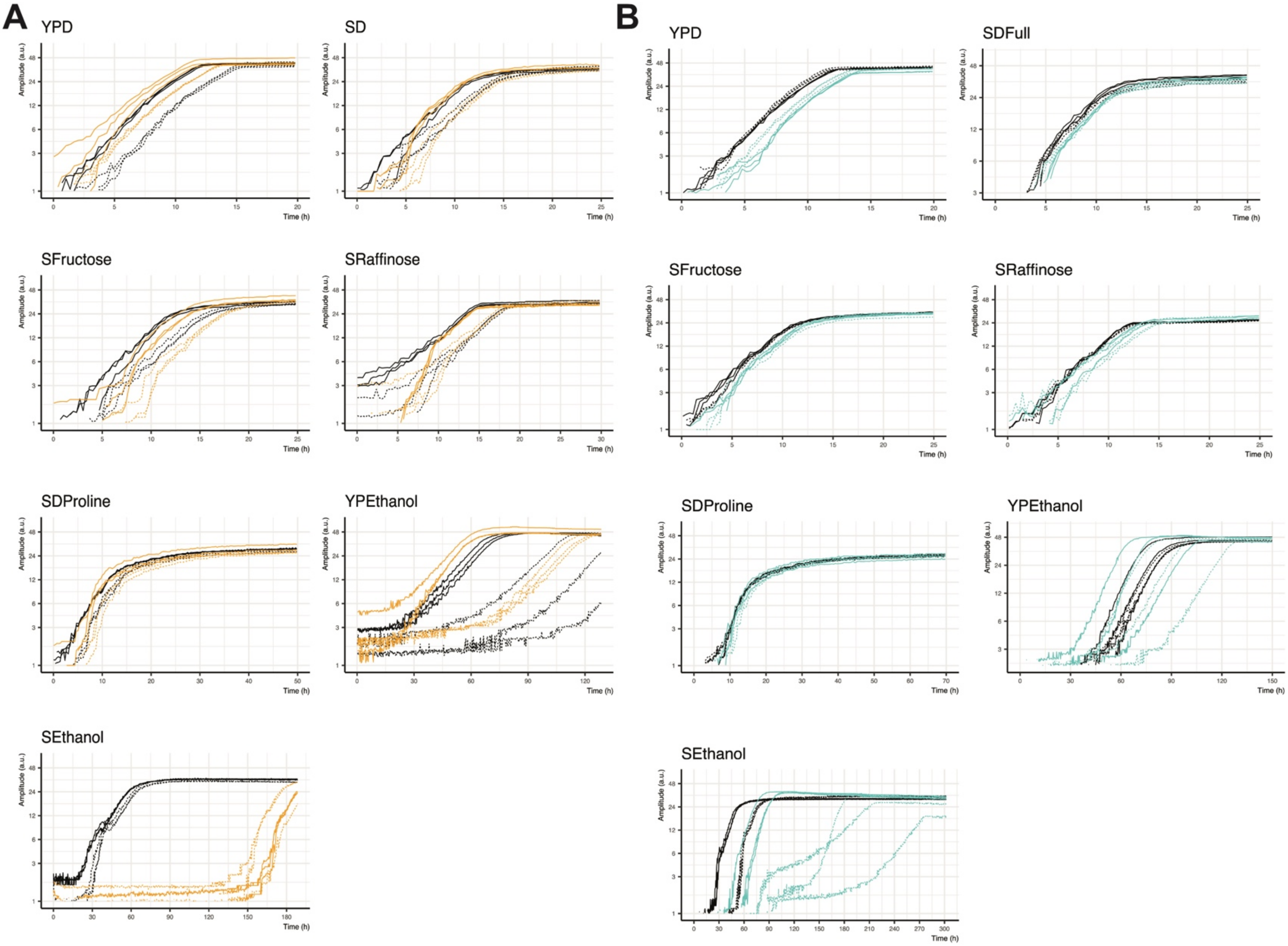


**Figure S7. Growth behaviour of cells bearing WTC$_{847}$ and WTC$_{848}$ under different media conditions.** Cells bearing either **A)** WTC$_{847}$ (Y3809) or **B)** WTC$_{848}$ (Y3870) were grown in different media conditions in a Growth Profiler 360, which photographs transparent-bottom plates and quantifies culture density based on the opacity of the imaged well every 20 minutes. Each well was seeded with 100.000 cells in 250$\mu$L media. Time 0 was arbitrarily chosen to be close to the time point where growth in the first well was observed, hence lag times between media conditions cannot be compared. Black lines indicate parental strain (Y70), turquoise lines indicate cells bearing WTC$_{848}$ and orange lines indicate cells bearing WTC$_{847}$. Solid lines indicate no inducer was added to the culture, dashed lines indicate either 20mM IPTG (WTC$_{848}$) or 30$\mu$M β-estradiol (WTC$_{847}$) was added. Media components are described in Methods.

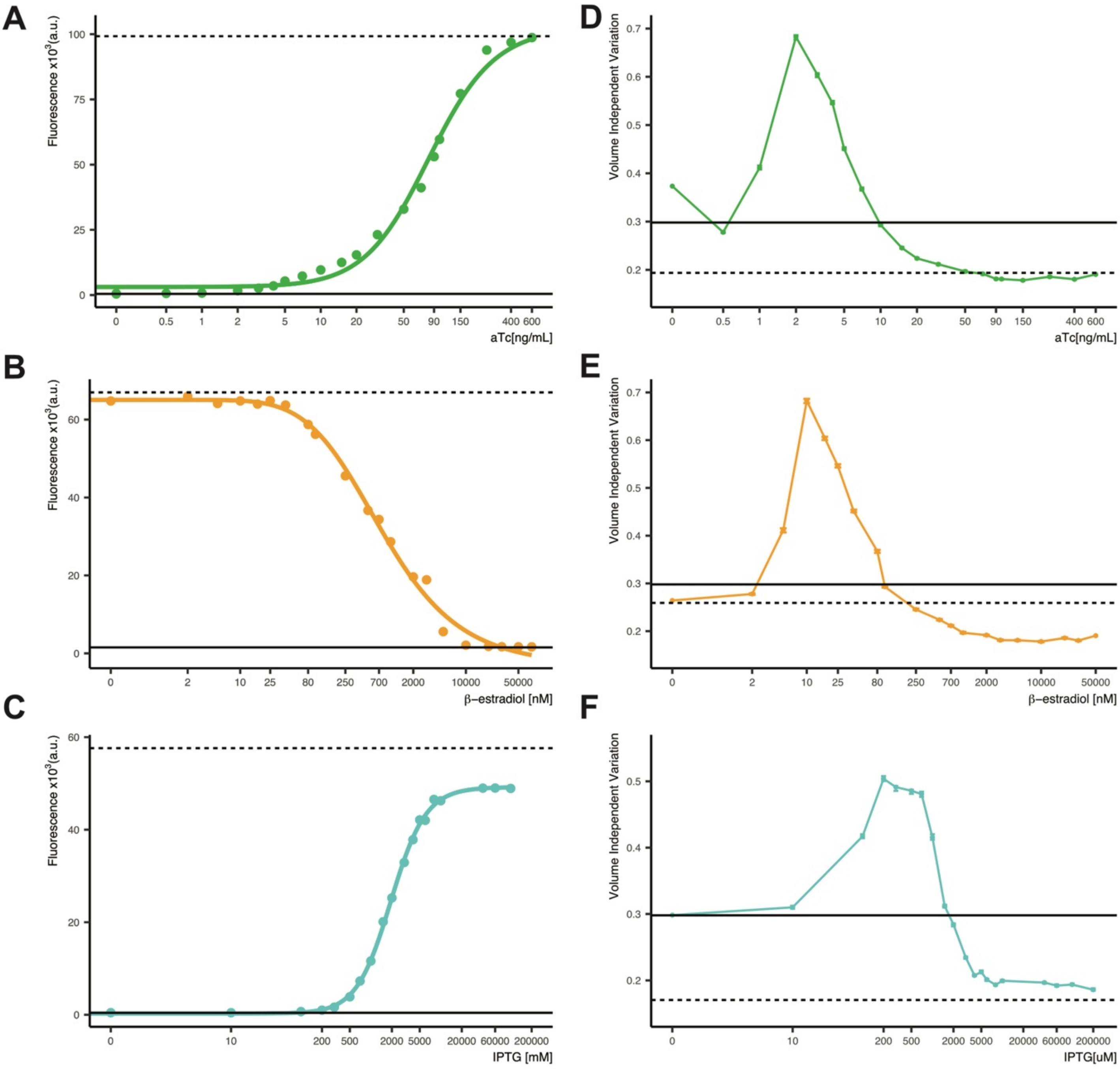


**Figure S8. The three WTC systems behave orthogonally in terms of dose response and cell-to-cell variation. A)** WTC$_{846}$, **B)** WTC$_{847}$ and **C)** WTC$_{848}$ dose response behaviour in YPD using strains where all three WTC systems were genomically incorporated (Y3877, 3880). Citrine (A,C) or mCardinal (B) fluorescence was measured with flow cytometry after 7 hours (n=5000 cells per dose) of growth and during exponential phase. Symbols indicate the median fluorescence of the measured population at each dose. Lines are fitted using a five-parameter log logistic function as explained in Methods. Dashed line indicates autofluorescence signal measured from the parental strain without Citrine (Y70). Dot-dashed line indicates fluorescence measured from a strain where the WTC promoter alone (P$_{7tet.1}$/P$_{7SulA.1}$/P$_{4Lacn.2}$) drove Citrine/mCardinal/Citrine expression (Y3728,3724,3542). **D-F)** Cell-to-cell variation of expression in the strains shown in (A-C). Higher Volume Independent Variation values (y axis) correspond to greater variation in expression, calculated as explained in the text and previously in[1]. Dot-dash line indicates the variation of the strain where Citrine is constitutively expressed from P$_{7SulA.1}$ and dashed line indicates variation of autofluorescence in the parent strain without Citrine. Error bars indicate 95% confidence interval calculated using bootstrapping (n=1000) as described in Methods.

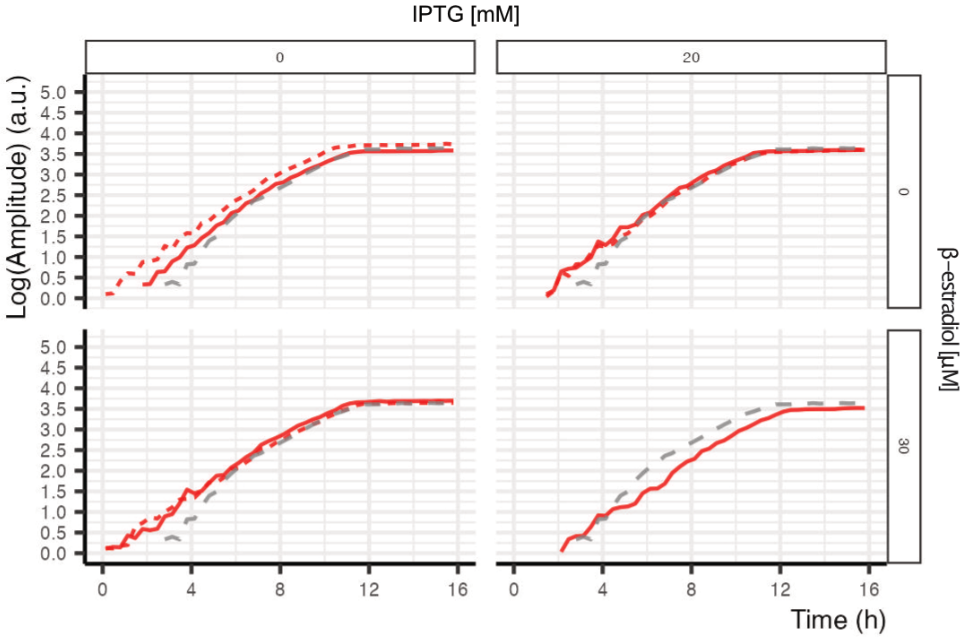


**Figure S9. Simultaneous presence and activation of the three WTC systems do not impair cell growth.** A strain bearing all 3 WTC systems and Citrine under the control of $P_{4Lacn.2}$ (Y3880, red) and the parent strain (Y70, grey) were grown in absence or presence of aTc, β-estradiol and IPTG in YPD. Growth of the culture was recorded using a Growth Profiler 360, which photographs transparent-bottom plates and quantifies culture density based on the opacity of the imaged well every 20 minutes. Growth curves were compared between the two strains to ascertain whether the WTC systems impaired cell physiology and caused growth delays. Solid lines indicate no aTc, dashed lines indicate presence of 600ng/mL aTc.

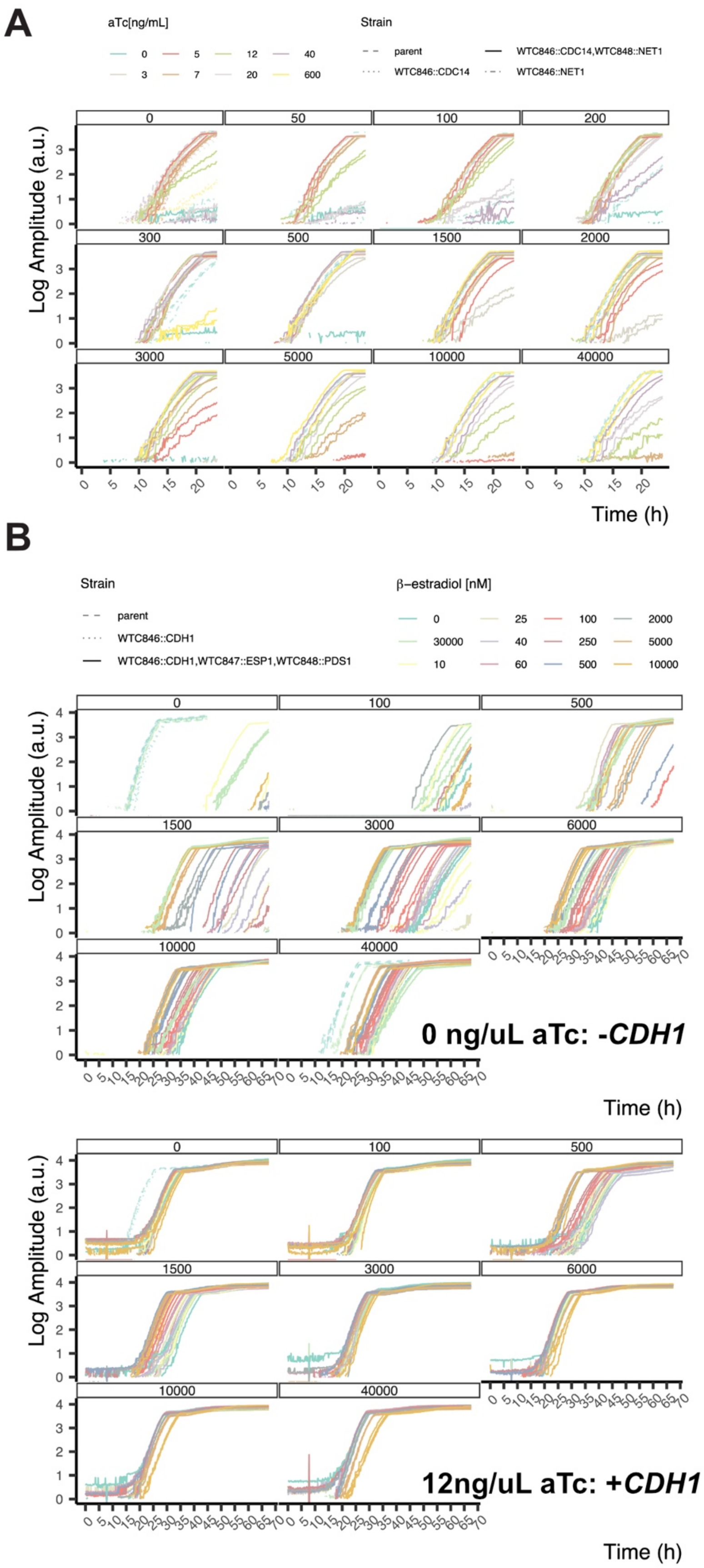


**Figure S10. Growth curves of titration experiments corresponding to Main Figure 3. A)** All cells were grown overnight with low (5ng/mL) aTc. *$CDC14^{846}NET1^{848}$* cells were additionally provided 10$\mu$M IPTG. In the morning cells were spun down and washed twice with YPD, followed by 6 hours of growth in YPD without inducers. Cells were then diluted again into plate reader plates with varying amounts of inducer. Each facet corresponds to one IPTG concentration in $\mu$M. Amplitude refers to measurements of culture density by the Growth Profiler 360 (Y3878,3925,3932,3945). **B)** Same as in A, except cells were grown with 12ng/$\mu$L aTc overnight and *$CDH1^{846}ESP1^{847}PDS1^{848}$*cells were additionally given 30 $\mu$M β-estradiol (Y3878,3928,3979).

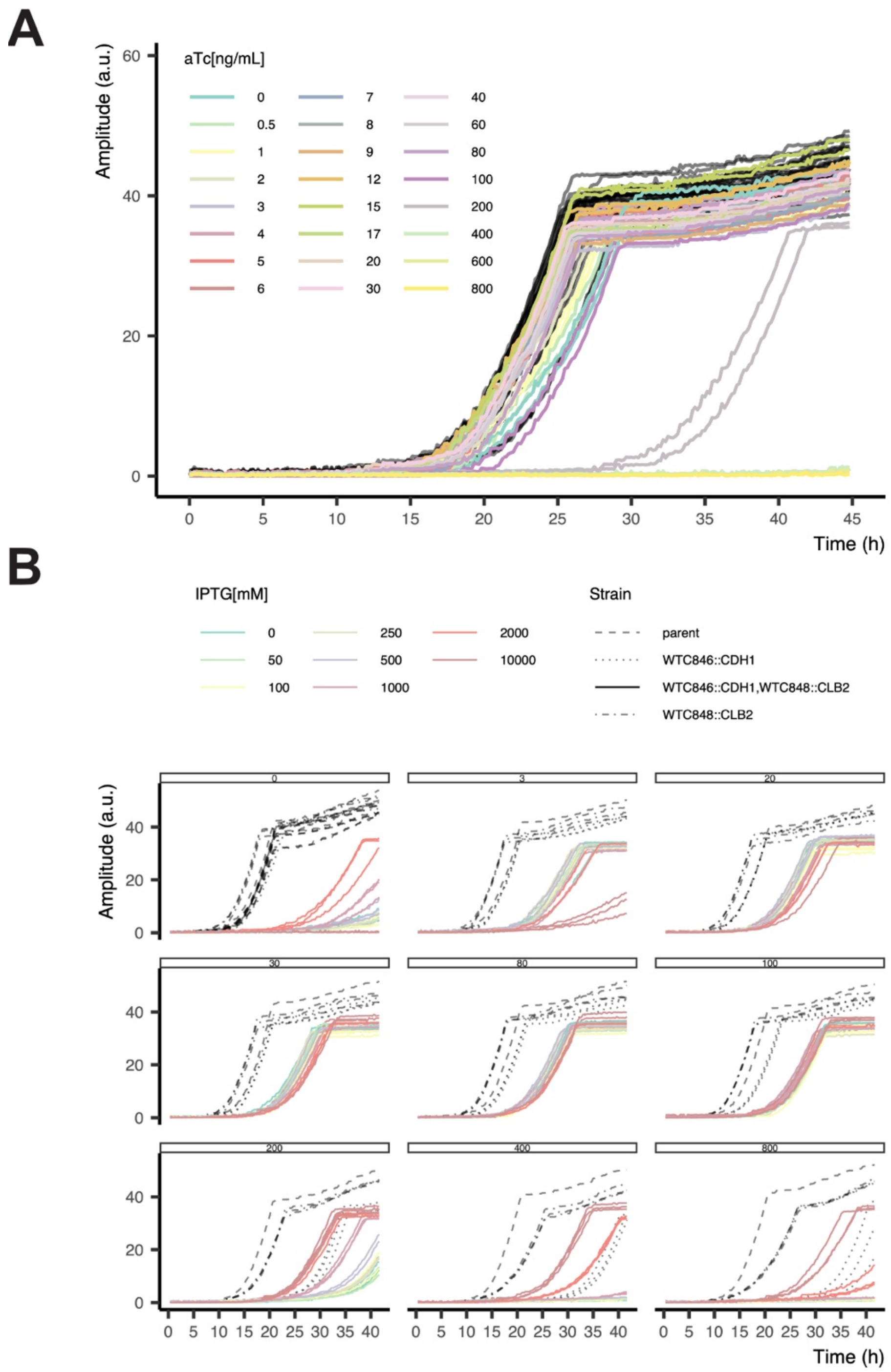


**Figure S11. Growth curves of titration experiments corresponding to Main Figure 4. A)** $\textit{CDH1}^{846}$ cells were grown overnight with low (10ng/mL) aTc. In the morning cells were spun down and washed twice with YPD, followed by 6 hours of growth in YPD without inducers. Cells were then diluted again into plate reader plates with varying amounts of aTc. Amplitude refers to measurements of culture density by the Growth Profiler 360 (Y3928, 3878) . **B)** Same as in A, except for $\textit{CLB2}^{848}$ cells, which were grown with 50$\mu$M IPTG overnight and $\textit{CDH1}^{846}\textit{CLB2}^{848}$ cells, which were grown with 10ng/$\mu$L aTc and 50mM IPTG overnight. (Y3878, 3928, 3923, 4043).

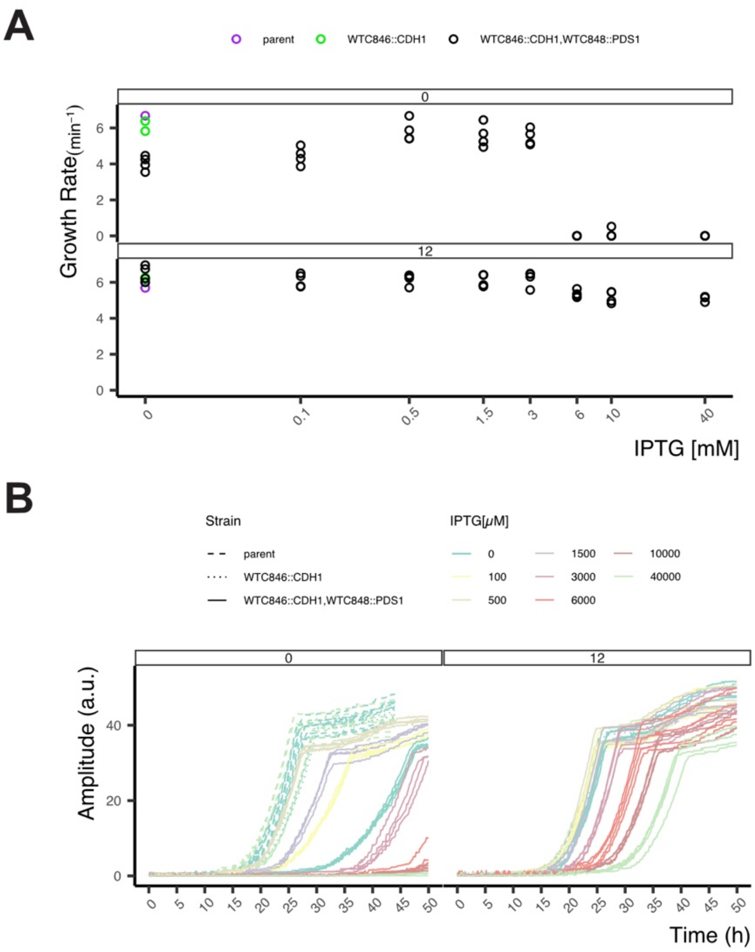


**Figure S12: Pds1 overexpression reduces fitness when Cdh1 is absent and Esp1 is under endogenous control.** Cells were grown overnight with low (10ng/mL) aTc. In the morning cells were spun down and washed twice with YPD, followed by 6 hours of growth in YPD without inducers. Cells were then diluted again into plate reader plates with varying amounts of aTc and IPTG. Maximum slope observed during exponential growth was quantified as a proxy for growth rates and fitness. Each facet indicates one aTc concentration in ng/μL. *$CDH1^{846}$* cells were grown in duplicates, *$CDH1^{846}PDS1^{848}$* in triplicates. Each circle represents an independently growing culture. **B)** Growth curves of the cells seen in (A). Amplitude refers to measurements of culture density by the Growth Profiler 360. (Y3878, 3928, 3947).
